# scDesignPop generates realistic population-scale single-cell RNA-seq for power analysis, benchmarking, and privacy protection

**DOI:** 10.64898/2026.02.23.707578

**Authors:** Chris Y. Dong, Yihui Cen, Dongyuan Song, Jingyi Jessica Li

## Abstract

Single-cell RNA sequencing (scRNA-seq) combined with genotyping in large cohorts has enabled the discovery of genetic associations with molecular traits (e.g., eQTLs) at cell-type resolution. However, generating population-scale data remains cost-prohibitive, selecting appropriate analysis methods lacks consensus, and sharing eQTL results alongside scRNA-seq data raises privacy risks. To address these challenges, we introduce scDesignPop, a flexible statistical simulator for generating realistic population-scale scRNA-seq data with genetic effects. scDesignPop models cell- and individual-level covariates, putative cell-type-specific eQTLs (cts-eQTLs), and either real or synthetic genotypes. We validated scDesignPop using the OneK1K and CLUES cohorts across 4 qualitative and 16 quantitative metrics. Unlike splatPop, the only existing population-scale simulator, scDesignPop better preserves eQTL effects and gene-gene dependencies within cell types, closely recapitulating characteristics of the reference data. Leveraging its generative framework, scDesignPop enables power analysis in cell types under multiple eQTL model specifications to guide experimental design; facilitates benchmarking of single-cell eQTL mapping methods through user-defined ground truths; and mitigates re-identification risk using synthetic data while retaining cts-eQTL effects.

## 1 Introduction

Single-cell RNA sequencing (scRNA-seq) has become a powerful tool for quantifying gene expression and understanding cellular heterogeneity at an unprecedented resolution. Combined with other genomic technologies, scRNA-seq has revealed gene regulatory mechanisms and potential drug target sites in specific cell types [1]. A common approach for studying gene regulation is expression quantitative trait loci (eQTL) analysis, which finds genetic variants linked to gene expression variations at the RNA level. While eQTL studies traditionally use bulk RNA-seq data, advances in scRNA-seq now enable population-scale single-cell eQTL studies that pair scRNA-seq with genotyping, allowing eQTL analysis at cell-type level across individuals [1–4]. Recent population-scale single-cell eQTL (sc-eQTL) studies have revealed both cell-type-specific and context-specific eQTL effects in various human tissues [5–11]. Notably, the OneK1K and CLUES cohorts revealed that many *cis*-eQTLs are highly cell-type-specific in peripheral blood mononuclear cells (PBMCs) [5, 6], with *cis*-eQTL single-nucleotide polymorphisms (*cis*-eSNPs) residing at different loci across cell types. Meanwhile, Natri et al. found most *cis*-eSNPs were shared among cell types in lung tissue [7]. Together, these findings highlight the growing importance of sc-eQTL studies for understanding cell-type-specific genetic regulation.

While population-scale sc-eQTL studies can offer cell-type-level insights that bulk eQTL analyses cannot resolve, there are several challenges. First, sc-eQTL studies are expensive due to the high cost of scRNA-seq. Second, the growing number of sc-eQTL methods makes it difficult for researchers to optimize analysis workflow. Third, sc-eQTL study results raise privacy concerns, as recent works highlighted [12, 13]. We discuss these challenges in detail below.

First, population-scale sc-eQTL studies are cost-prohibitive because detecting cell-type-specific eQTLs (cts-eQTLs), which typically have modest effect sizes, requires sufficient samples (e.g., individuals, cells, and sequencing depth). The sparsity and high-dimensionality of scRNA-seq further compound the difficulty of detecting eQTLs. Power analysis offers a principled approach for estimating the minimum samples needed to achieve a desired power, thereby guiding resource-efficient study design. However, existing power analysis tools for sc-eQTL studies, powerEQTL [14] and scPower [15], have limitations: powerEQTL relies entirely on user-specified parameters rather than being data-driven, reducing realism, while scPower assumes a simple linear model for eQTL mapping and lacks alternative model options. Moreover, neither accounts for covariates such as experimental conditions or cell-type heterogeneity. These limitations underscore the need for a flexible, data-driven power analysis framework that supports different models and real-world experimental complexities.

Second, researchers face numerous method and workflow choices in sc-eQTL analysis, including preprocessing (data quality control, demultiplexing [16–18], doublet removal [19–21], cell-type annotation), batch-effect correction [22], normalization [23–28], eQTL mapping [5, 29–32], and multiple testing correction [33–35]. Different combinations can lead to markedly different eQTL discovery results even on the same dataset [36, 37]. The emergence of sc-eQTL methods that model single-cell counts directly [38–40], rather than pseudobulk expression, further underscores the need for systematic benchmarking [41], yet there is still a lack of gold-standard datasets with known ground truths for cts-eQTLs. Simulated scRNA-seq data with defined eQTL effects, therefore, provide a practical solution for method benchmarking. Among existing scRNA-seq simulators [42–48], muscat [46] and rescueSim [47] explicitly model multiple individuals; however, both are limited to generating synthetic scRNA-seq data within the same individuals as in the reference data and do not incorporate genetic effects such as eQTLs. In contrast, splatPop [48] models eQTL effects but primarily relies on a *de novo* simulation framework that requires numerous user-specified parameters, thus limiting its ability to closely mimic real data even when reference scRNA-seq data are provided as input. As population-scale eQTL studies involve increasingly complex covariates, such as multiple disease statuses and experimental conditions [7, 10, 11], there is a growing need for a flexible simulator that generates realistic scRNA-seq data with interpretable parameters, incorporates eQTL effects, and accounts for both cell- and individual-level covariates.

Third, the genomic data in sc-eQTL studies pose privacy risks due to sensitive genotype information [49–51]. Adversaries can exploit publicly available eQTLs and gene expression data to infer genotypes [12, 13, 52], and then link these inferred genotypes to private databases, thereby re-identifying individuals and revealing sensitive phenotypic information. Mitigation strategies include data encryption for storage and transfer, access control, and data perturbation [49, 53, 54]. However, the first two strategies cannot fully prevent unscrupulous users from re-identifying individuals. Although data perturbation-based approaches such as differential privacy can offer privacy guarantees [55], applying them to scRNA-seq and genotype data while preserving biological signals remains challenging. Although generative models for synthetic data have been proposed to enhance privacy protection [56, 57], none have been explored in the sc-eQTL field.

One solution to address these challenges is to generate realistic, population-scale scRNA-seq data that preserves biological characteristics such as cts-eQTLs. Such simulations can generate synthetic cells either for the real individuals in the reference data or new individuals with real or synthetic genotypes. These synthetic data enable applications such as conducting power analysis, benchmarking methods under realistic population heterogeneity, protecting genomic privacy, and potentially exploring rare individual-level covariate combinations underrepresented in the original data.

Here, we propose **scDesignPop**, a reference-based simulator that generates realistic population-scale scRNA-seq data by learning from paired scRNA-seq and genotype data with putative eQTLs (Fig. 1). Compared to scDesign3 [58], which is well-suited for simulating single-cell and spatial omics data modalities within an individual sample, and existing multi-individual simulators [46–48], scDesignPop distinguishes itself by explicitly modeling the genetic effects present in the population-scale data. Specifically, scDesignPop introduces three key innovations: (1) modeling cts-eQTLs using multiple SNPs, flexible cell- and individual-level covariates, and their interactions; (2) generating realistic new individuals by modeling inter-individual variations and cell-type composition; and (3) integrating eQTL power analysis in cell types as a built-in functionality. Supplementary Tables S1 and S2 summarize the functionalities of scDesignPop in comparison with scDesign3, existing multi-individual scRNA-seq simulators, and current eQTL power analysis tools.

**Fig 1:**
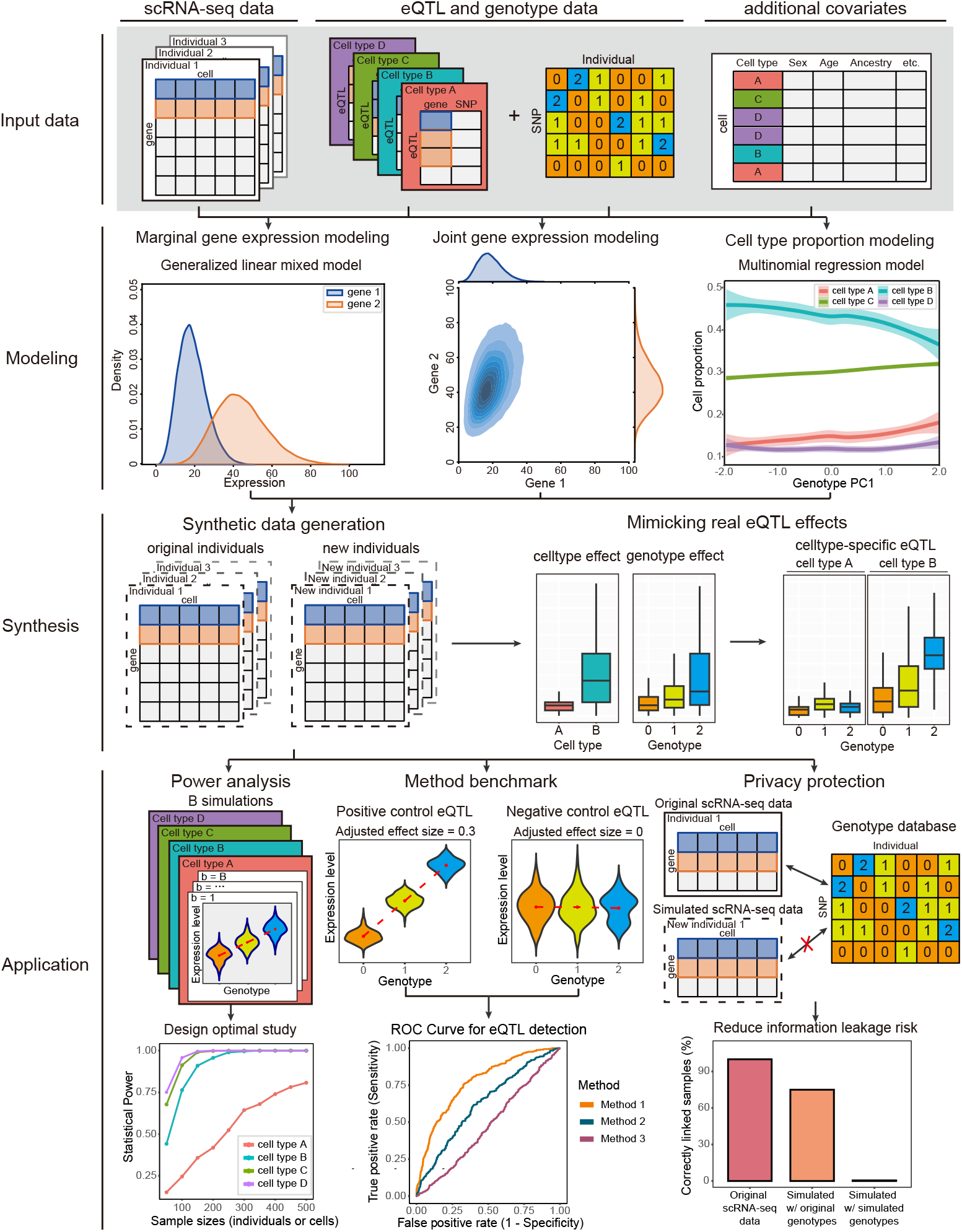
Overview of the scDesignPop framework. scDesignPop requires three input components: population-scale scRNA-seq data, putative eQTL annotations with corresponding genotype matrices (gene-SNP pairs per cell type; genotypes coded as 0, 1, or 2), and additional cell- or individual-level covariates (for example, cell type, sex, and age). The model fitting step comprises three main components: (1) a marginal model for gene expression across cells using a generalized linear mixed model, (2) a joint model for gene expression with gene-gene dependence captured by a Gaussian copula, and (3) a multinomial logistic regression model for simulating cell-type proportions when generating new individuals. The synthesis step generates synthetic scRNA-seq data for both existing and new individuals using either estimated or user-specified eQTL effect sizes. scDesignPop’s downstream applications include (1) cell-type-specific eQTL power analysis for study design, (2) benchmarking of sc-eQTL mapping methods, and (3) privacy-preserving data sharing by releasing simulated datasets that reduce re-identification risk.

scDesignPop flexibly models both count and normalized data, supports user-specified cell- and individual-level covariates, and accommodates both real and synthetic genotypes. The fitted model can also be adjusted with user-specified eQTL effect sizes and used to generate varying numbers of cells and individuals for power analysis and method benchmarking. We first show that scDesignPop preserves realistic cell-type proportions, genes’ marginal and joint expression distributions, and cts-eQTL effects in the OneK1K and CLUES cohorts (Results 2.2, 2.4), justifying scDesignPop’s modeling framework as reasonable and expressive. Next, we show that scDesignPop can also generate realistic dynamic eQTLs along continuous cell trajectories (Results 2.3). Finally, by simulating new individuals, we highlight scDesignPop’s utility for performing eQTL power analysis (Results 2.5), benchmarking sc-eQTL mapping methods (Results 2.6), mitigating genomic privacy risks (Results 2.7), and scaling beyond the original cohorts (Results 2.8). scDesignPop is available as an R package at https://github.com/chrisycd/scDesignPop with tutorials at https://chrisycd.github.io/scDesignPop/docs/index.html.

## 2 Results

### 2.1 Overview of scDesignPop’s framework

scRNA-seq data generated with droplet-based technologies such as 10x Genomics and CITE-seq use unique molecular identifiers (UMIs) to quantify gene expression. Both negative binomial (NB)- and Poisson-based models are commonly used to model UMI counts [59, 60]. In addition, downstream processing steps such as normalization or batch correction may yield non-integer expression values, which are typically modeled by a Gaussian distribution. To accommodate these scenarios, scDesignPop allows users to specify models for either counts (NB- or Poisson-based) or normalized expression levels (Gaussian-based). scDesignPop is a reference-based simulator that takes as input paired scRNA-seq and genotype data, putative cts-eQTLs, and cell- or individual-level covariates such as cell type and ancestry (input data panel, Fig. 1). For putative cts-eQTLs, we recommend using results from prior sc-eQTL studies or databases such as scQTLbase [61]. From these reference data, scDesignPop estimates parameters of flexible models (e.g., allowing interaction effects between genotypes and other covariates), allows users to modify parameters to reflect desired cts-eQTL ground truths, and generates synthetic scRNA-seq data accordingly. The generative framework of scDesignPop consists of three modeling components (modeling panel, Fig. 1) and a data generation component (synthesis panel, Fig. 1): (1) modeling each gene’s marginal distribution across cells given cell- and individual-level covariates, (2) modeling the joint distribution of genes given cell- and individual-level covariates, (3) modeling cell-type proportions within individuals given individual-level covariates, and (4) generating synthetic scRNA-seq data given covariates for same or new individuals. Each component is described in Methods 10.1.

### 2.2 scDesignPop generates realistic population-scale scRNA-seq data in OneK1K and CLUES cohorts

We validated scDesignPop as a realistic simulator using two independent population-scale scRNA-seq datasets, the OneK1K and CLUES cohorts, with distinct data characteristics. The OneK1K cohort comprises 1.27 million peripheral blood mononuclear cells (PBMCs) from 981 individuals across 14 immune cell types as count data [5]. The CLUES cohort comprises 1.23 million PBMCs from 256 individuals of different ancestries and systemic lupus erythematosus (SLE) disease status across 8 cell types, provided as normalized expression data [6]. Because splatPop is the only existing population-scale scRNA-seq simulator [48], we compared it with scDesignPop under both single-SNP and multi-SNP modes (scDesignPop_single and scDesignPop_multi; Methods 10.2) using held-out test data from the OneK1K cohort. As splatPop requires count-based input, it was not evaluated on the normalized CLUES dataset.

Across 16 evaluation metrics, scDesignPop-simulated data more closely resembled real data than splatPop in OneK1K and recapitulated real data well in CLUES (Methods 10.4, 10.5). Using two-dimensional UMAP embeddings (uniform manifold approximation and projection) [62], scDesignPop-simulated data consistently achieved high similarity with real, held-out test data across both cohorts. In OneK1K, scDesignPop attained higher mean local inverse Simpson’s index (mLISI) scores, which measure the resemblance between simulated data and test data, than splatPop at the overall (Fig. 2A), genotype-specific (e.g., *BLK*; Figs. 2B), and individual-specific levels (Fig. 2D). Similarly, in CLUES, scDesignPop achieved high mLISI values (≥ 1.722) across ancestry and disease-status groups relative to test data at group-specific (Fig. 3A), genotype-specific (e.g., *GZMB*; Fig. S7), and joint group-genotype levels (e.g., *GZMB*; Figs. S8). Interestingly, the simulated data for healthy individuals of European or Asian ancestry exhibited higher mLISI values than those for SLE patients with Asian ancestry (Fig. 3D), potentially reflecting greater heterogeneity among SLE patients, whose treatment information was unavailable as model covariates to be included in scDesignPop. Beyond UMAP embedding similarity, scDesignPop also preserved individual-level gene expressions with respect to relevant subgroups (age and sex in OneK1K; Fig. 2C; ancestry and disease status in CLUES; Fig. 3B), closely matched the distributions of the test data for summary statistics at the gene-, cell-, and pseudobulk-levels (Figs. S1, S2), and maintained gene-gene correlations across all cells (“allct”) and multiple representative cell types using their top 100 expressed genes (Figs. 2E, 3C).

**Fig 2:**
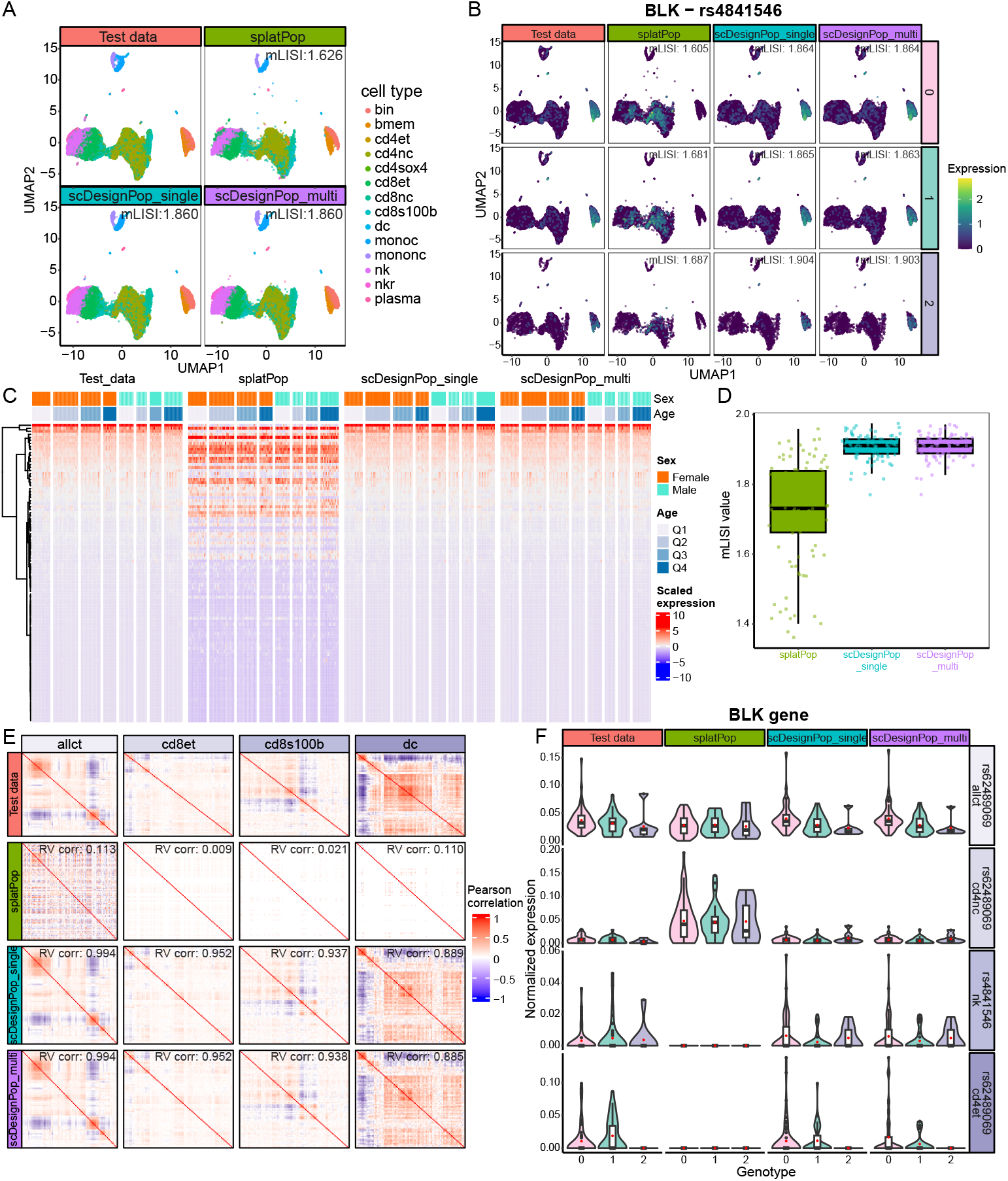
scDesignPop more faithfully recapitulates OneK1K cohort than splatPop. **(A)** UMAP embeddings colored by cell type for held-out test data, splatPop-simulated data (Methods 10.4), and scDesignPop-simulated data under the single-SNP (scDesignPop_single) and multi-SNP (scDesignPop_multi) modes (Methods 10.2). mLISI values quantify how similar each simulated dataset is to the test data. **(B)** UMAP embeddings of *BLK* expression across genotypes (0, 1, 2) of SNP rs4841546 in each dataset; mLISI values are computed within each genotype. **(C)** Heatmaps of scaled pseudobulk gene expression for the top 100 highly variable genes (HVGs; identified by Seurat’s default “vst” method) across individuals, stratified by sex and age quartiles. **(D)** Boxplots of mLISI values computed by comparing cells from each individual in the test data with cells from the same individual in the simulated data. **(E)** Gene-gene correlation matrices for all cells (allct), CD8+ effector memory T cells (cd8et), CD8+ S100B cells (cd8s100b), and dendritic cells (dc), shown for test data and three simulated datasets. Pearson correlations were computed from normalized data for the top 100 expressed genes within each cell type. **(F)** Violin plots of pseudobulk *BLK* expression across eSNP genotypes for selected cts-eQTLs (rs62489069 in all cells (allct); rs62489069 in CD4+ naive/central memory T cells (cd4nc); rs4841546 in natural killer cells (nk); rs62489069 in CD4+ effector memory T cells (cd4et)). Red points indicate mean expression.

**Fig 3:**
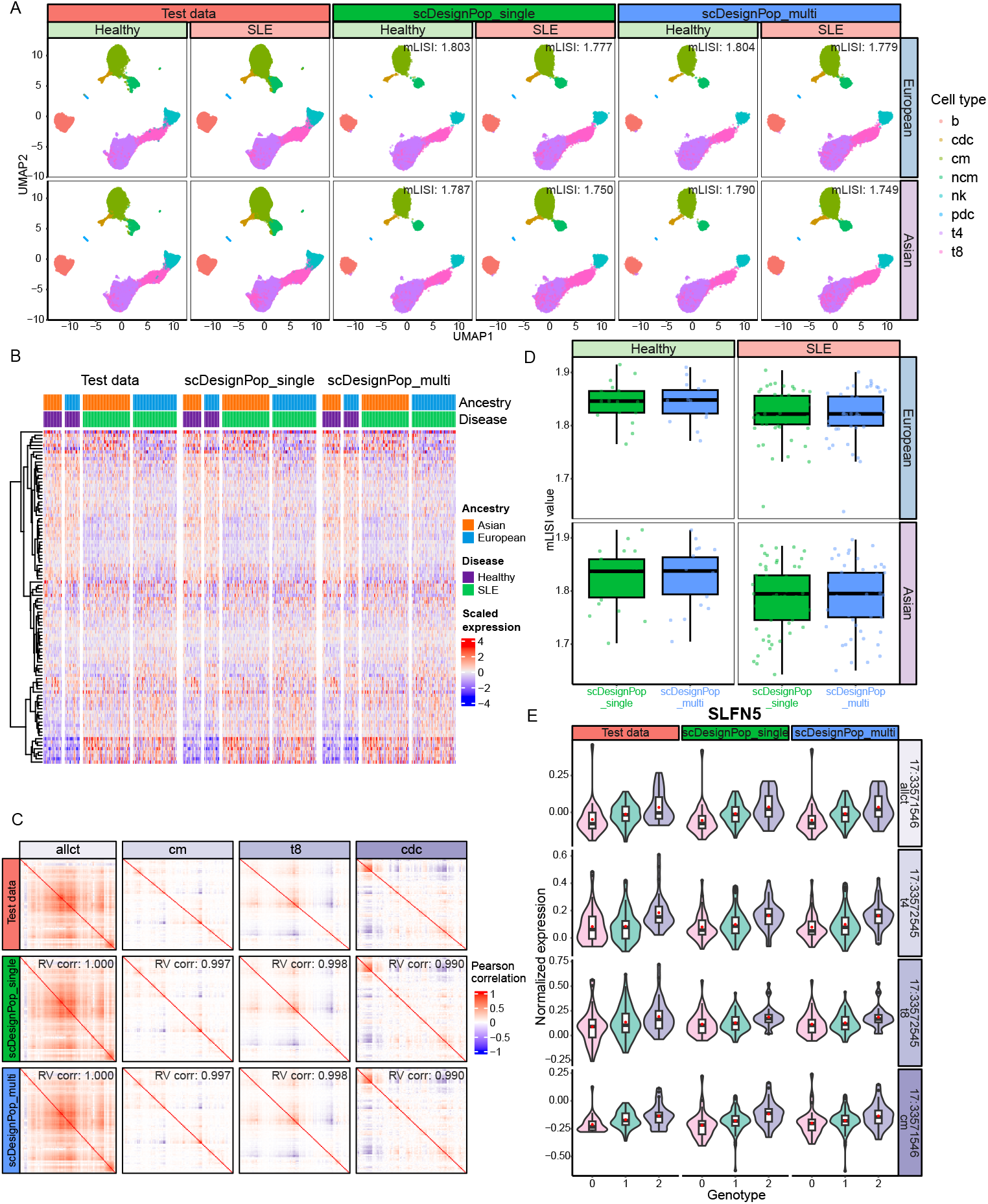
scDesignPop preserves complex ancestry and disease structure in CLUES cohort. **(A)** UMAP embeddings colored by cell type for held-out test data, and scDesignPop-simulated data under both single-SNP (scDesignPop_single) and multi-SNP modes (scDesignPop_multi; details in Methods 10.2)). mLISI values quantify how similar each simulated dataset is to the test data within each disease status and ancestry group. **(B)** Heatmap of scaled pseudobulk gene expression for the top 100 highly variable genes (HVGs) across individuals, stratified by disease statuses and ancestries. The top HVGs were identified using Seurat’s default “vst” method. Gene expression was aggregated at the individual level and scaled prior to visualization. **(C)** Gene-gene correlation matrices for all cells (allct), classical monocytes (cm), CD8+ T cells (t8), and conventional dendritic cells (cdc), shown for test and simulated datasets. Pearson correlation was computed using the normalized test or simulated data. **(D)** Boxplots of mLISI values computed by comparing cells from each individual in the test data with cells from the same individual in the simulated data, stratified by disease status and ancestry. **(E)** Violin plots of pseudobulk *SLFN5* expression across eSNP genotypes for selected cts-eQTLs at their SNP coordinates (17:33571546 in all cells (allct); 17:33572545 in CD4+ T cells (t4); 17:33572545 in CD8+ T cells (t8); 17:33571546 in classical monocytes (cm)). Red points indicate mean expression.

Importantly, scDesignPop more accurately recapitulated cts-eQTL effects at the pseudobulk level compared to the real data in both cohorts. For all cts-eQTLs, Spearman correlations between pseudobulk gene expression and eSNP genotypes more closely matched those in the held-out test data for scDesignPop simulations than for splatPop across all cell types (Fig. S3, S4). In OneK1K, this was reflected by higher coefficients of determination, computed between Spearman correlations from test and simulated data, for scDesignPop (*R*^2^ = 0.36 − 0.75) compared with splatPop (*R*^2^ = 0.00 − 0.05). A likely explanation is that splatPop models the user-specified cts-eQTL effect sizes across genes using a parametric distribution, rather than modeling within each gene with its own distribution. Thus, although we supplied splatPop with Matrix eQTL slope estimates from the OneK1K study [5], its framework does not preserve cts-eQTL effects that are inherently gene-specific (Methods 10.4).

Between the two scDesignPop modes, the multi-SNP mode generally preserved cts-eQTL effects more accurately than the single-SNP mode. Across both cohorts, simulated data under the multi-SNP mode yielded higher *R*^2^ values (defined the same as above) relative to those from the single-SNP mode (Figs. S3, S4), indicating that modeling multiple *cis*-eSNPs better captures the underlying *cis*-regulatory effects of gene expression. In OneK1K, the *R*^2^ ranged from 0.42 − 0.67 for scDesignPop_single and 0.36 − 0.75 for scDesignPop_multi; in CLUES, the *R*^2^ ranged from 0.64 − 0.79 for scDesignPop_single and 0.70 − 0.81 for scDesignPop_multi. Exceptions were observed in plasma, CD4_SOX4_ T, and CD8_S100B_ T cells in OneK1K. The overall higher *R*^2^ values in CLUES likely reflect its 3.7-fold greater number of cells profiled per individual and more individuals in the training data (150 CLUES vs. 100 OneK1K individuals).

scDesignPop’s ability to recapitulate cts-eQTL effects is further illustrated using *BLK* and *SLFN5* genes as examples from the OneK1K and CLUES datasets, respectively. For the *BLK* gene, both scDesignPop modes closely mimicked its expression levels (Fig. 2B, F). In contrast, although splatPop captured the negative association between *BLK* expression and genotype using all cells (“allct”) shown in Fig. 2F, it generated little or no *BLK* expression in memory B cells (bmem) and immature/naive B cells (bin) on UMAP embeddings (Fig. 2B), as well as in natural killer (nk) and CD4_ET_ (cd4et) cells (Fig. 2F). splatPop also overexpressed *BLK* in CD4_NC_ T cells (cd4nc) relative to the test data (Fig. 2F). These instances of under- or over-expression of genes in specific cell types, are consistent with splatPop’s modeling strategy explained above, which does not fully preserve gene-specific cts-eQTL effects. As a result, the simulated expression levels of certain genes in specific cell types may deviate from those observed in the real data. Similarly, for *SLFN5*, a gene implicated in type I interferon activation in SLE [6], scDesignPop again showed close resemblance between simulated and test data at a pseudobulk level for all cells (“allct”), and for several representative cell types (Fig. 3E) in the CLUES cohort.

The reason underlying the above results is that scDesignPop requires only putative cts-eQTL loci, not user-specified effect sizes. scDesignPop infers cts-eQTL effects directly from reference data using a unified marginal model per gene across cell types, thereby preserving empirical effects of cts-eQTLs and generating realistic simulated scRNA-seq data. To assess parameter stability, we subsampled individuals from the OneK1K cohort and refit the marginal model (Methods S2). As expected, both cell-type and cts-eQTL effect estimates, along with their standard errors, stabilized as the number of individuals increased, though this varied across cell types (Fig. S5, S6). Notably, cts-eQTL effect estimates and their uncertainty stabilized with approximately 50 − 100 individuals, suggesting that cohorts of this scale may be sufficient to obtain reliable estimates in similar study settings.

### 2.3 scDesignPop flexibly models dynamic eQTL effects along continuous cell trajectories

scDesignPop also models dynamic eQTL effects along continuous cell trajectories (i.e., pseudotime), allowing genetic effects to vary smoothly across cellular processes such as differentiation or stimulation [63]. Following Yazar et al. [5], we inferred pseudotime for B cells, consisting of immature/naive and memory B cells, then evaluated scDesignPop’s ability to simulate both linear and nonlinear dynamic eQTLs [64], by using pseudotime as a cell-state covariate with a genotype-pseudotime interaction term for linear dynamic eQTLs, and additional quadratic pseudotime for nonlinear dynamic eQTLs (Methods 10.7).

To illustrate scDesignPop’s ability to model dynamic eQTLs, we compared gene expression trends along pseudotime between the reference and simulated data under both linear and nonlinear dynamic eQTL models. Using PHATE (potential of heat-diffusion for affinity-based trajectory embeddings) [65], scDesignPop preserved the manifold structure of the reference data under both linear and nonlinear models (Fig. 4A). For linear dynamic eQTLs, scDesignPop reproduced the linear relationship between *BLK* gene expression and pseudotime observed in the reference data (Fig. 4B). For nonlinear dynamic eQTLs, scDesignPop captured the more complex genotype-pseudotime relationships exhibited by *CD27, SMCO4*, and *TCL1A* genes in the reference data (Fig. 4C, D). Together, these results demonstrate that scDesignPop can realistically model both linear and nonlinear dynamic eQTL effects along continuous cell trajectories.

**Fig 4:**
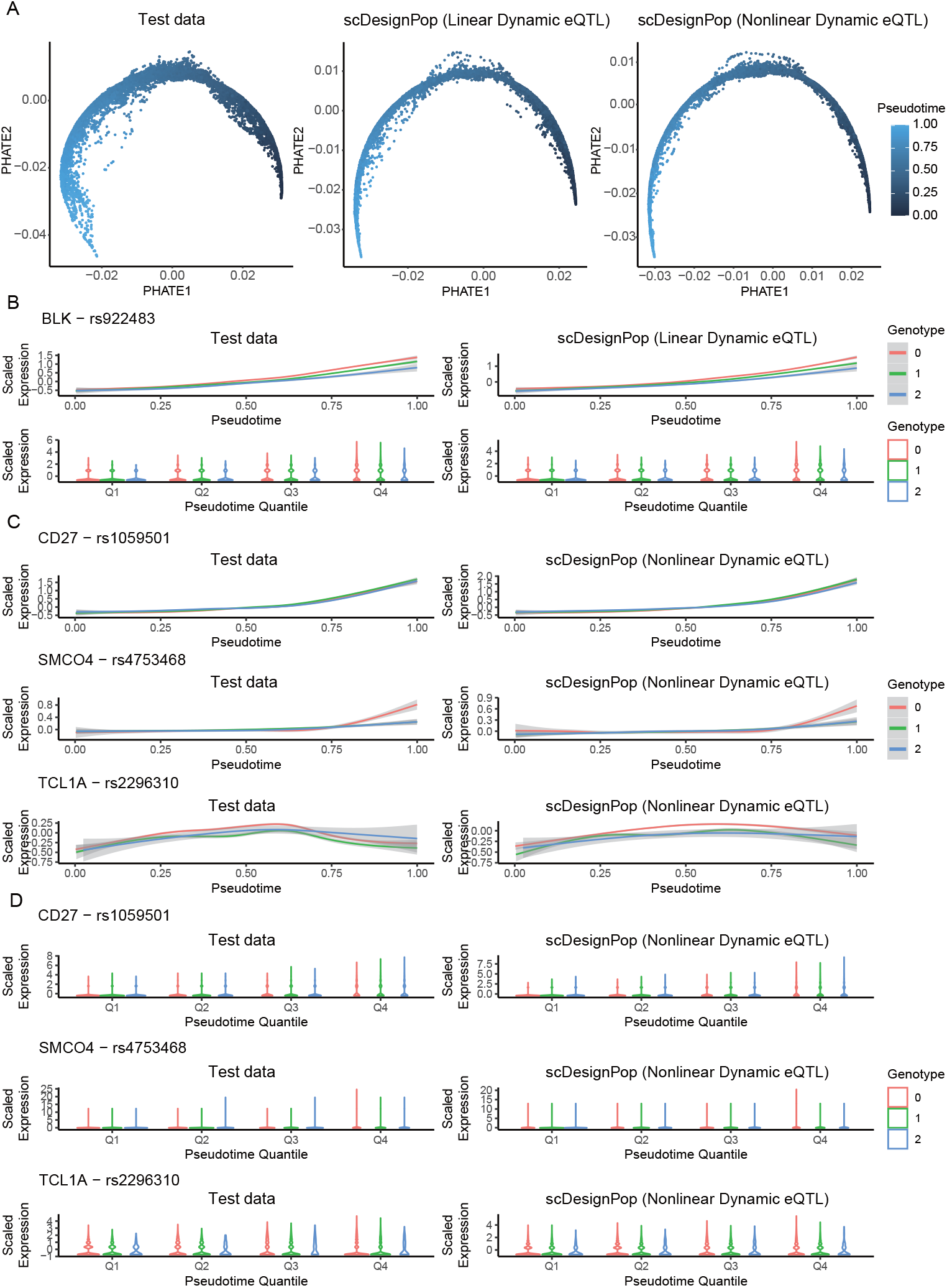
scDesignPop captures linear and nonlinear dynamic eQTL effects. **(A)** PHATE embeddings of the B cell population (immature/naive B cells (bin) and memory B cells (bmem)) from the OneK1K reference data. **(B)** *BLK* expression (scaled log-normalized counts) across rs922483 genotypes along pseudotime in test data and scDesignPop-simulated data under a linear dynamic eQTL model. Smooth curves depict fitted trends; violin plots summarize scaled log-normalized expression within pseudotime quartiles (Q1–Q4). **(C-D)** Gene expressions (scaled log-normalized counts) along pseudotime for representative gene-SNP pairs under a nonlinear dynamic eQTL model, shown for test and simulated data. Smooth curves represent fitted trajectories; violin plots correspond to pseudotime quartiles (Q1–Q4).

### 2.4 scDesignPop simulates scRNA-seq data for new individuals with real or synthetic genotypes

Population-scale sc-eQTL datasets are limited to the individuals originally profiled, which constrains power analysis, restricts method benchmarking, and poses privacy risks when sharing these data. scDesignPop addresses these limitations by simulating *new* individuals, either using real genotypes from held-out test data or fully synthetic genotypes. We show that these simulated new individuals exhibit realistic cell-type compositions and gene expressions compared with the reference data.

Using 70% of individuals in each reference dataset for training (70 individuals in OneK1K and 105 individuals in CLUES), scDesignPop accurately recovered the cell-type proportions of the remaining 30% test individuals (Figs. S9, S10, S11; Methods 10.1.4). Moreover, when provided with 982 synthetic individuals whose genotypes were generated using HAPGEN2 [66] (Methods 10.6), scDesignPop simulated scRNA-seq data in over 1.3 million synthetic cells whose expression closely resembled those from 981 individuals in the OneK1K cohort. This is illustrated by the gene expression distributions of *IL7R* and *SOX4* marker-genes for CD8_S100B_ and CD4_SOX4_ T cells respectively, examined across all cells (“allct”) and within each cell type (Figs. S12, S13).

Together, these results demonstrate scDesignPop’s ability to generate realistic population-scale scRNA-seq data for new individuals beyond the training data. This capability substantially broadens its utility for downstream applications, including power analysis, method benchmarking, and privacy risk mitigation.

### 2.5 scDesignPop provides flexible power analysis for detecting cell-type-specific eQTL effects

scDesignPop provides a flexible, data-driven framework for power analysis of cts-eQTLs across multiple eQTL models. Because closed-form power calculations are generally unavailable for GLMMs [67–69], scDesignPop uses simulation-based power calculations based on real data (Methods 10.8). Compared with powerEQTL [14] and scPower [15], two existing eQTL power calculation tools that rely on user-specified input parameters and simplified modeling assumptions, scDesignPop can incorporate covariates such as sex and age, yielding more realistic power estimates.

scDesignPop’s power analysis framework accounts for both the number of individuals and the number of cells per individual, two key study-design parameters in population-scale sc-eQTL studies [36, 70]. Using cts-eQTLs for *LYZ* (OneK1K) and *GZMB* (CLUES), we observed that power depends strongly on the effect size within each cell type. For *LYZ* in the OneK1K cohort, in classical monocytes, where the effect size is relatively large (−0.321), power increased clearly with more individuals or more cells (Fig. S14). In memory B cells, however, the same eQTL has a much smaller effect size (0.021) and the gene is lowly expressed, resulting in consistently low power even with larger sample sizes (Fig. S16, S17)). Similar trends were seen for *GZMB* in the CLUES cohort: power increased with sample size in CD8+ T cells (effect size 0.075) but not in nonclassical monocytes (effect size 0.002) (Fig. S15, S18). scDesignPop also supports user-specified effect sizes, and as expected, power increased as the effect size grew (Fig. S19, S20). Overall, these relationships among effect size, sample size, and power are consistent with previous findings [71–73].

scDesignPop supports four eQTL models for power analysis: negative binomial (NB) mixed, Poisson mixed, linear mixed models for single-cell-level data, and a pseudobulk linear model for data aggregated at the individual-cell-type level (i.e., one pseudobulk profile per cell type per individual). We did not include a zero-inflated negative binomial (ZINB) model, as for UMI-based scRNA-seq data used in cohort studies, zero inflation is not needed and the NB distribution is sufficient [74]. In the OneK1K cohort, NB mixed and Poisson mixed models yielded higher power than linear mixed and pseudobulk linear models for the highly expressed *LYZ* gene across multiple cell types (Fig. S14). In contrast, for the lowly expressed *S100B* gene, linear mixed and pseudobulk linear models performed comparably or better (Fig. 5), consistent with Zhou et al.’s observations [38]. This highlights that the most appropriate eQTL model depends on both gene expression level and the data representation (single-cell vs. pseudobulk).

**Fig 5:**
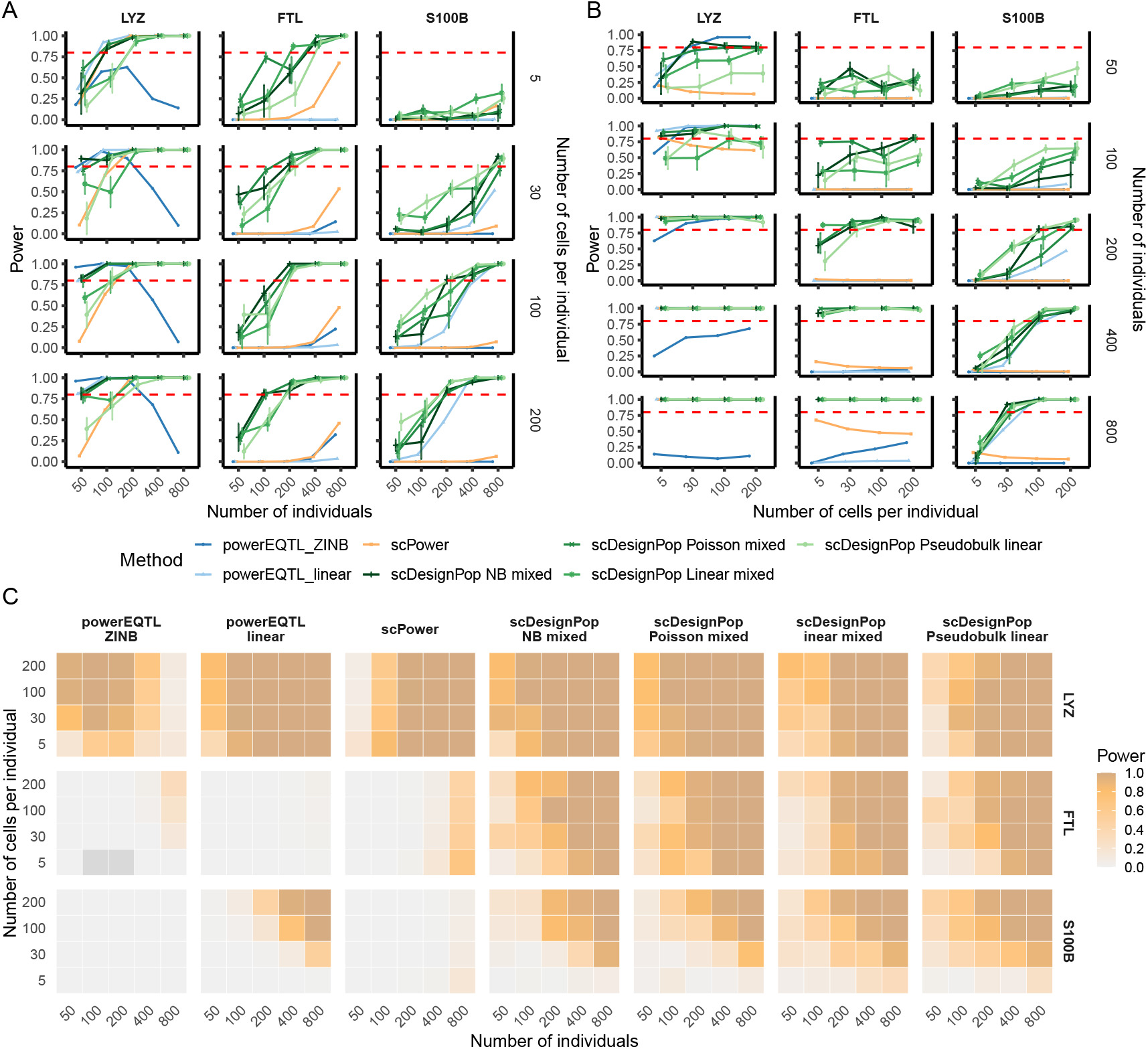
Comparison of eQTL power from scDesignPop, powerEQTL, and scPower in classical monocytes from the OneK1K cohort. **(A)** Line plots showing power as a function of the number of individuals for detecting the eQTL effect of *LYZ* (SNP 12:69732105), *FTL* (SNP 21:48039367), and *S100B* (SNP 19:49540093). The red dashed line denotes 80% power. **(B)** Line plots showing power as a function of the number of cells per individual for the same eQTLs. The red dashed line denotes 80% power. **(C)** Heatmaps summarizing power for detecting eQTL effects for *LYZ* (SNP 12:69732105), *FTL* (SNP 21:48039367), and *S100B* (SNP 19:49540093) changes across the number of individuals or the number of cells per individual changes. Dark grey cells indicate parameter settings for which power was not obtained. For powerEQTL, two options (powerEQTL_ZINB and powerEQTL_linear) were used; for scDesignPop, four eQTL model options were used (Methods 10.9).

Next, we compared scDesignPop’s power analysis with powerEQTL [14] and scPower [15]. Both tools require user-specified parameters, which we estimated from the reference data following their model assumptions (Methods 10.9). For powerEQTL, we evaluated both its simulation-based ZINB mixed model and its analytical linear mixed model. For scPower, we used its analytical power calculation under its assumed simple linear model. As an example, we evaluated eQTL power for *LYZ, FTL*, and *S100B* genes in classical monocytes from the OneK1K cohort. Compared with powerEQTL and scPower, scDesignPop’s power estimates aligned more closely with expected behavior across tested conditions and eQTL model choices. Leveraging real gene expression and genotype data, scDesignPop’s power estimates increased with more individuals or more cells per individual. In contrast, powerEQTL’s ZINB mixed model produced unexpected decreases in power for *LYZ* as the number of individuals increased (Fig. 5A, C), and its linear mixed model showed plateauing power for *FTL* with increasing individuals or cells (Fig. 5A-C), likely due to sensitivity to fixed parameters such as intra-class correlation and residual variance. scPower, although it incorporates real data, relies on parameters derived from a limited bulk eQTL reference and showed a monotonic decrease in overall detection power as the number of cells per individual increased for all three genes (Fig. 5B, C)). This behavior stems from its definition of “overall power” as the product of a “gene expression probability,” which approaches 1 for highly expressed genes, and an “eQTL power” term that does not depend on the number of cells; increasing cells per individual inflates the number of expressed genes, thereby increasing the multiple-testing burden and reducing overall power. Because scPower does not allow users to specify the number of tested genes, this setup diverges from how real eQTL analyses are designed. Overall, scDesignPop avoids these breakdowns and consistently produces the expected increases in power with larger sample sizes across its four model options, enabling realistic and flexible power analysis for cts-eQTL studies.

### 2.6 scDesignPop’s interpretable parameters enable benchmarking of eQTL mapping methods

scDesignPop’s interpretable model parameters enable users to define ground-truth cts-eQTL genes (cts-eGenes), defined as genes with at least one eQTL in a given cell type, and cts-non-eGenes (the complement of cts-eGenes). We demonstrate its application by benchmarking sc-eQTL mapping methods using the OneK1K cohort as reference data. First, we illustrate how ground truth can be constructed from fitted or user-modified effect-size parameters. Then, we benchmark three representative eQTL mapping methods: FastQTL, jaxQTL (in both “permutation” and “ACAT-V” modes), and SAIGE-QTL, using scDesignPop-simulated data.

scDesignPop-simulated scRNA-seq data can either mimic the reference data (using fitted parameters) or incorporate user-defined modifications to weaken or strengthen eQTL effects. cts-eGenes or cts-non-eGenes can be defined based on fitted or modified parameters. Fig. 6A shows four parameter settings for the *BLK* gene and the SNP locus rs62489069 in memory B cells (bmem). Because the OneK1K study did not identify this gene-SNP pair as a putative eQTL in this cell type, *BLK* was defined as a cts-non-eGene by setting its effect size to zero and lowering its mean expression. Conversely, to define it as a cts-eGene, users may amplify the eQTL effect size (e.g., by a log_2_ fold-change increase or to an effect of equal magnitude but with an opposite sign). Importantly, even after these parameter modifications, the simulated B cells remained similar to cells in the reference data at each genotype in UMAP plots (Fig. S22, S23), as only one gene was altered in a cell-type-specific manner. scDesignPop’s modifyMarginalModels function supports batch editing of effect sizes across multiple genes.

**Fig 6:**
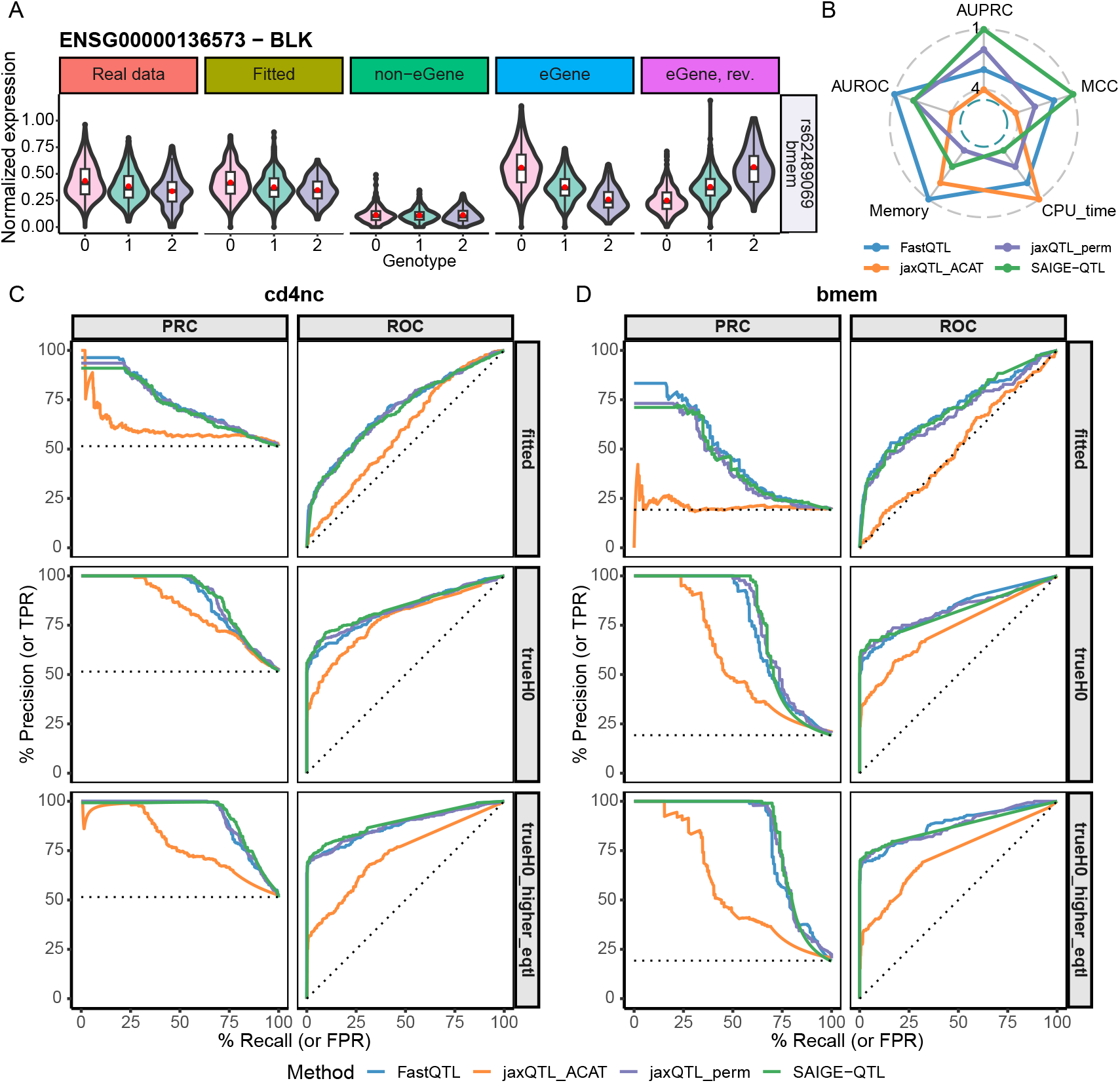
Benchmarking eQTL mapping methods using scDesignPop under different simulation settings. **(A)** Violin plots of pseudobulk *BLK* expression across SNP (rs62489069) genotypes in memory B cells (bmem) comparing real data to scDesignPop-simulated data under fitted, or modified parameters to define a non-eGene or eGene. For the non-eGene, the eQTL effect size was set to zero and its mean expression was lowered by − 2 log_2_ fold-change. For the eGene, the eQTL effect size was amplified by 2 log_2_ fold-change, with its direction either matching the same sign as the fitted or reversed (rev.). **(B)** Radar plot showing the ranks of five performance metrics for eQTL mapping methods. CPU_time represents the total computational time in hours, and memory represents the peak memory usage in gibibytes. **(C-D)** Performance of eQTL mapping methods in CD4_NC_ T cells (**C**) and memory B cells (**D**) using scDesignPop-simulated data generated under three parameter settings: fitted parameters (fitted), zero effect sizes for cts-non-eGenes (`trueH0`), and zero effect sizes for cts-non-eGenes with increased effect sizes for cts-eGenes (`trueH0_higher_eqtl`) (10.10). Precision-recall (left) and receiver operating characteristic (right) curves are shown for each method in each cell type. The dotted line denotes random (no-skill) guessing. *Abbreviations*: non-eGene (non-eQTL gene); eGene (eQTL gene); memory B cells (bmem); CD4_NC_ T cells (cd4nc); Area under the receiver operating characteristic curve (AUROC); area under the precision-recall curve (AUPRC); Matthew’s correlation coefficient (MCC); false positive rate (FPR); true positive rate (TPR); jaxQTL with set-based aggregated Cauchy association test (jaxQTL_ACAT); jaxQTL with beta-approximation using permutations (jaxQTL_perm).

Next, we benchmarked three eQTL mapping methods that represent the two major analysis paradigms: FastQTL and jaxQTL (permutation and ACAT-V modes), which operate on pseudobulk gene expression, and SAIGE-QTL, which directly models single-cell counts. FastQTL has been widely used in eQTL studies, whereas jaxQTL and SAIGE-QTL were recently developed, underscoring the need for a fair comparison for the field. We compared these three methods using scDesignPop-simulated data from the OneK1K CD4_NC_ T cells (473 average cells per individual) and memory B cells (49 average cells per individual). Three simulation settings were used: (1) fitted parameters (fitted), (2) zero effect sizes for cts-non-eGenes (trueH0), and (3) zero effect sizes for cts-non-eGenes plus a 0.5 log_2_ fold-change increased effect sizes for cts-eGenes (trueH0_higher_eqtl). These settings create progressively greater separation between the effect sizes of cts-non-eGenes (null hypothesis: H0) and cts-eGenes (alternative hypothesis). For each simulation setting, we generated approximately 1.2 million cells in 981 individuals for 810 genes using real genotypes and cell type labels (Methods 10.10).

Based on the ranks of five performance metrics we evaluated, FastQTL and SAIGE-QTL outperformed jaxQTL overall. FastQTL achieved the lowest computational time and higher area under the ROC curve (AUROC) (Fig. 6B, Supplementary Table S4), whereas SAIGE-QTL attained higher area under the precision-recall curve (AUPRC), and Matthew’s correlation coefficient (MCC) at the 5% target FDR threshold. Under the fitted simulation setting, FastQTL outperformed the other methods in AUROC, AUPRC, and MCC across both cell types (Fig. 6C, D). In contrast, under trueH0 and trueH0_higher_eqtl settings, SAIGE-QTL achieved the highest AUPRC in CD4_NC_ cells and the highest MCC in both cell types. Notably, jaxQTL_ACAT performed much worse than jaxQTL_perm across both cell types (Fig. 6C, D), possibly reflecting implementation differences between the two modes, although jaxQTL_ACAT had the shortest computational time among all methods (Supplementary Table S4). Overall, these results illustrate scDesignPop’s flexibility in benchmarking both pseudobulk-based and single-cell-based eQTL mapping methods under user-defined ground-truth effect sizes.

### 2.7 scDesignPop protects genomic information leakage by mitigating eQTL-based linking attack

Individuals in eQTL studies are vulnerable to re-identification because eQTLs can be used to link scRNA-seq profiles to genotype data. PrivaSeq, introduced by Harmanci and Gerstein [12], demonstrated this risk using bulk RNA-seq, and recent work by Walker et al. [13] showed that PrivaSeq’s eQTL-based linking attacks remain feasible for scRNA-seq data, even when using only CD4_NC_ T cells. However, whether simulation of scRNA-seq data from population-scale eQTL studies can effectively mitigate these privacy risks remains unknown. We therefore applied scDesignPop to generate realistic synthetic scRNA-seq data and tested its ability to mitigate eQTL-based linking attacks.

Across a range of gap statistic thresholds, which define the minimum reliability required for a correct link, scDesignPop-simulated data substantially reduced the success of linking attacks when paired with reference genotypes (OneK1K) and nearly eliminated it when paired with synthetic genotypes (Methods 10.11). With a gap statistic threshold of zero, 98.1% of the 981 individuals were correctly linked between the reference scRNA-seq and genotype data (solid red line, Fig. 7A, B), consistent with Walker et al.’s results for CD4_NC_ T cells [13]. This high success rate highlights strong re-identification risk when releasing real scRNA-seq data. In contrast, the correct linking proportion was reduced to 78.7% when using scDesignPop-simulated scRNA-seq paired with reference genotypes (dashed teal line, Fig. 7B) and further down to 0.0% when using scDesignPop-simulated scRNA-seq paired with synthetic genotypes generated by HAPGEN2 [66] (dashed purple line, Fig. 7B). Hence, using simulated data with synthetic genotypes yielded a correct linking proportion comparable to the 0.2% obtained when permuting pseudobulk expression across individuals in the reference data (solid green line, Fig. 7B), which removes all eQTL effects and represents a best-case scenario for minimizing linking attack risk. Figure 7C further shows this reduction in linking attack risk, where the gap statistic distributions for both scDesignPop-simulated datasets are shifted lower than the reference data (two-sample Wilcoxon *p*-values of 1.17 × 10^−194^ and 3.29 × 10^−295^).

**Fig 7:**
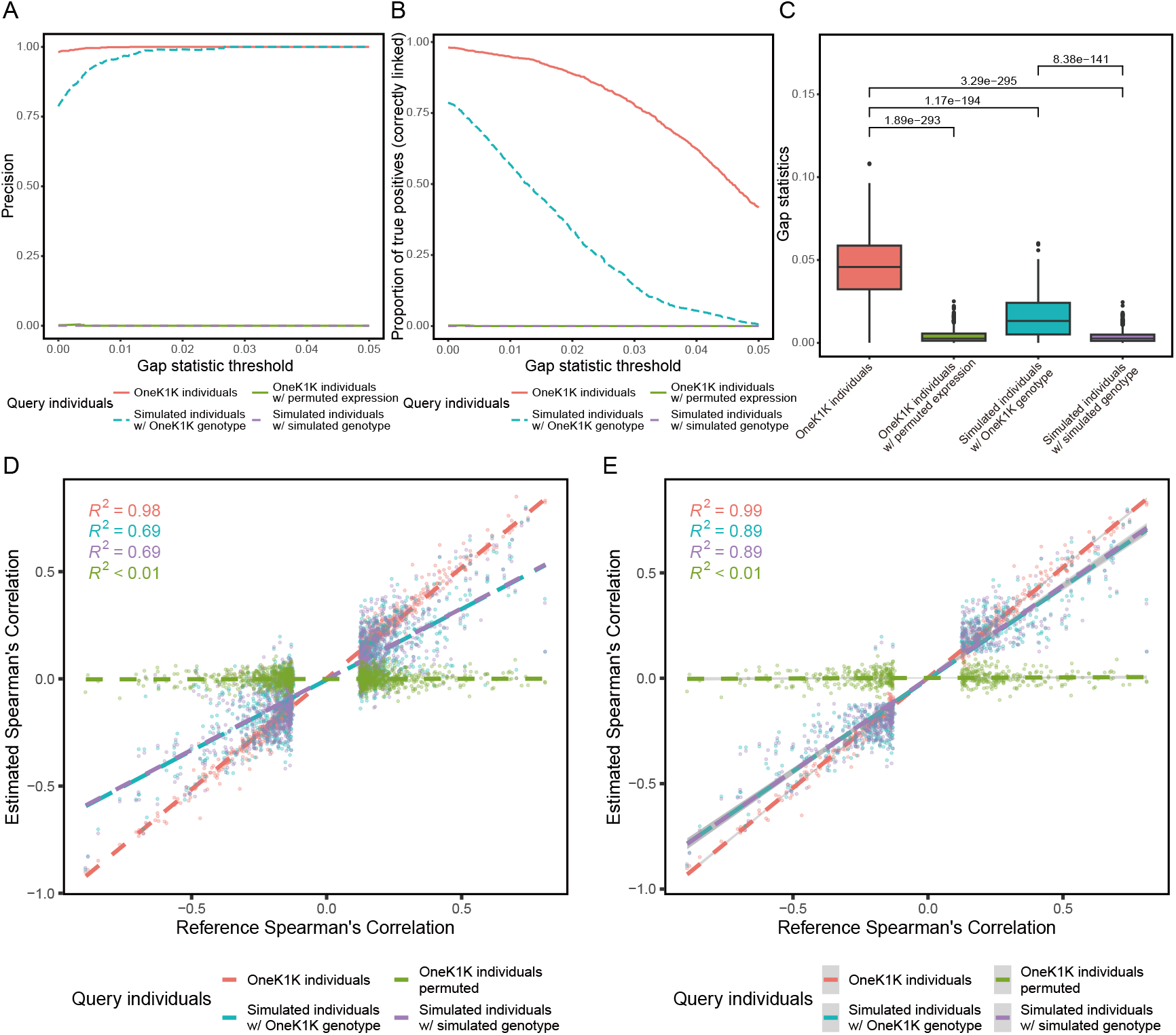
scDesignPop mitigates eQTL-based linking attacks while preserving eQTL effects in simulated data. Linking attack performance was evaluated using four datasets: (1) original OneK1K individuals; (2) OneK1K individuals with permuted pseudobulk expression; (3) scDesignPop-simulated individuals paired with OneK1K genotypes; and (4) scDesignPop-simulated individuals paired with synthetic genotypes generated by HAPGEN2. **(A)** Precision (ratio of true positives among all positive links) as a function of the gap statistic threshold chosen by the attacker. **(B)** Proportion of correctly linked individuals across gap statistic thresholds. **(C)** Distributions of gap statistics for the four datasets. Statistical significance was assessed using a two-sided Wilcoxon rank-sum test. **(D-E)** Scatterplots showing the relationship between estimated Spearman correlations for each query dataset and reference Spearman correlations obtained from the OneK1K’s publicly available eQTL data for all 1,140 gene-SNP pairs **(D)**, and a subset of 530 gene-SNP pairs after filtering for those with estimated Spearman correlation above the minimum observed in the reference eQTL data **(E)**. Spearman correlation was computed between pseudobulk expression and the corresponding genotype. Dashed lines indicate least-squares linear regressions fitted to each query dataset with *R*^2^ values shown in corresponding colors. All panels use CD4_NC_ T cells, the most abundant cell type in the OneK1K dataset, as the representative example.

Importantly, unlike permuted data, scDesignPop-simulated data retained cts-eQTL effects, showing strong concordance with the cts-eQTL effects observed in the reference data. For each gene-SNP pair, we quantified the cts-eQTLs as the Spearman correlation between pseudobulk gene expression and genotype of putative eQTLs across individuals in both simulated and reference data from OneK1K. Across 1,140 pairs, simulated and reference data showed moderate concordance of *R*^2^ = 0.69 for both scDesignPop simulations, one with reference genotypes and the other with synthetic genotypes (dashed teal and purple lines, Fig. 7D). This moderate concordance likely reflects uncertainty in cts-eQTL estimation arising from the 100 sampled individuals used to train scDesignPop, compounded by a large portion of modest cts-eQTL effects. Also, using synthetic genotypes versus real genotypes had minimal impact, since both had nearly identical concordance. Consistent with this interpretation, the concordance increased to *R*^2^ = 0.89 after we focused on 530 gene-SNP pairs with absolute Spearman correlations in the simulated data above 0.121, which was the minimum correlation required to call eGenes at a 0.05 FDR in the reference data (dashed teal and purple lines, Fig. 7E). At the same time, permuting pseudobulk gene expression in the reference data removed all eQTL effects, yielding very low concordance of *R*^2^ < 0.01 with the reference (dashed green line, Fig. 7D, E), as expected. Together, these results demonstrate that scDesignPop-simulated scRNA-seq paired with synthetic genotypes preserves cts-eQTL effect signals while mitigating eQTL-based linking attack risks, thereby enabling privacy-conscious data sharing and supporting downstream analyses, particularly the development and benchmarking of eQTL mapping methods.

### 2.8 scDesignPop’s computation time and memory costs for simulation at population-scale and beyond

As population-scale scRNA-seq data size continues to grow, with the TenK10K project recently having profiled over 5 million cells from nearly 2,000 individuals [75], simulator scalability is essential for it to be of practical use. We therefore evaluated scDesignPop’s scalability with varying training data sizes, different modeling specifications, and for large synthetic data generation.

scDesignPop’s total computation time and memory usage increased linearly with the number of cells, *I*, exhibiting *𝒪* (*I*) complexity throughout model training and data generation (Fig. S24). We assessed scDesignPop’s performance from loading input data to generating synthetic data for varying numbers of subsampled individuals from OneK1K cohort to measure the total computation time and peak memory usage (Methods 10.12). To further assess the computational cost under different modeling specifications and data regimes, we measured the average time in gene marginal modeling, which is typically the rate-limiting step. Across datasets and marginal model configurations, per-gene computation time increased with both the number of cells and model complexity (e.g., inclusion of additional covariate terms), as expected. Average computation time ranged from 6.4 seconds per gene for approximately 92,000 cells to 17.9 seconds per gene with approximately 473,000 cells (Supplementary Table S6).

Crucially, scDesignPop’s generative framework supports biobank-scale simulation designs, whereby a model trained on a moderate cohort size can generate millions of synthetic cells for thousands of new individuals. As our validation results show, scDesignPop retained realistic cts-eQTL effects, despite using a relatively modest training subset (e.g., 100 individuals). Thus, to demonstrate scDesignPop’s scalability beyond reference cohort size, we generated up to 6.33 million synthetic cells and 4,905 individuals, which took under 18.9 CPU hours and utilized 597 GiB of peak memory (Supplementary Table S7), using a model trained on 70 individuals from the OneK1K cohort. These correspond to only approximately 1.4-fold and 2.2-fold greater CPU computation and peak memory cost, respectively, than for training and simulating the same number of 261,000 cells from 200 individuals on average. Thus, scDesignPop’s simulation flexibility enables single-cell data generation at population- and biobank-scale computationally feasible.

Finally, scDesignPop leverages parallel computing for both model fitting and synthetic data generation to alleviate the computational burden. Specifically, CPU parallelization is implemented in marginal and joint model fitting, parameter estimation, synthetic data generation, and power analysis modules. To maximize cross-operating-system compatibility and user flexibility, scDesignPop supports four parallelization backend options (Methods 10.12).

## 3 Discussion

scDesignPop addresses three critical bottlenecks in population-scale single-cell genomics: uncertainty in optimal study design for adequate power, proliferation of analysis methods lacking systematic benchmarking, and genomic privacy concerns limiting data sharing. By providing an interpretable, realistic simulation framework validated by both OneK1K and CLUES cohorts, scDesignPop enables researchers to perform cell-type-specific power analysis to guide study design, benchmark eQTL mapping methods under user-specified cts-eQTL ground truths, and mitigate genomic privacy risks through the synthetic data generation.

Our results underscore the generalizability of scDesignPop for single-cell eQTL studies in two directions: applicability to diverse tissues and support for complex study designs. First, as a reference-based simulator, scDesignPop is readily applicable to any tissue with available population-scale single-cell eQTL data. Although validated here using PBMCs, the same modeling framework naturally extends to population-scale single-cell datasets from diverse tissues, including lung [7], brain [10], and liver [76]. Second, scDesignPop’s flexibility in modeling through cell- and individual-level covariates enables a broad range of experimental designs such as longitudinal studies, imbalanced case-control settings, and multi-ancestry cohorts [77]. Diverse population cohorts can be modeled using genotype principal components (PCs) as individual-level covariates to account for fine-scale population structure.

scDesignPop’s interpretable framework further enables simulation of single-cell molecular QTL data beyond *cis*-eQTLs. For example, *trans*-eQTLs can be specified using putative *trans*-eQTL loci. More broadly, scDesignPop naturally extends to other molecular QTLs, including chromatin-accessibility QTLs [78–81] and protein QTLs [82], requiring only appropriate data modalities and corresponding distributional families. Future extensions may further incorporate transcript-level count models to support splicing and isoform QTLs, as well as multimodal frameworks that jointly model gene expression and chromatin accessibility at the single-cell level.

Importantly, our results highlight that synthetic genotypes are effective for mitigating privacy risks, but are insufficient on their own to ensure biological realism in simulated scRNA-seq data, which also depends on the simulation framework. Although synthetic genotypes could be incorporated into other population-scale scRNA-seq simulators, such as splatPop, to mitigate the risk of re-identifying real individuals, scDesignPop more faithfully preserves biological signals, such as gene-gene correlations and eQTL effects. This emphasizes the importance of jointly considering privacy protection and biological realism when using simulated data, which depends on the user’s specific use case and scientific objectives.

Despite the advantages of scDesignPop, several limitations remain, pointing to future directions. First, scDesignPop is a reference-based simulator and therefore depends on the availability of population-scale single-cell datasets for the tissue of interest. For tissues lacking such data, incorporating bulk eQTL summary statistics to guide parameter estimation may be a promising extension. Second, population-scale sc-eQTL studies frequently use multiplexing across batches. While scDesignPop can model batch effects in the reference data as fixed effects, adding support for simulating entirely new batches containing multiple individuals in each (e.g., through random effects) would further broaden its applicability. Third, although we showed that scDesignPop-simulated data reduced eQTL-based linking attack risks, we did not evaluate other forms of privacy attacks [49, 52] or other privacy metrics [50, 56]. Providing rigorous guarantees for privacy-utility tradeoffs remains an open challenge. Finally, memory overhead can become substantial when simulating biobank-scale datasets with millions of cells, highlighting opportunities for further optimization, such as on-disk data structures or GPU acceleration, without compromising simulation realism. Similarly, although scDesignPop can be seamlessly paired with HAPGEN2 to generate synthetic genotypes that retain realistic LD patterns, broader software support for additional genetic simulators, such as HAPNEST ([66, 83–85]), would further enhance flexibility.

## 4 Data and software resources

### Data resources

OneK1K eQTL data were obtained from the OneK1K website (https://onek1k.org). Pre-processed OneK1K scRNA-seq, Matrix eQTL, and genotype data was provided by the authors. CLUES scRNA-seq data and eQTL data were obtained under Gene Expression Omnibus (GEO) accession number GSE174188, and from the Supplementary Materials in [6], respectively. CLUES genotype data was obtained under dbGaP accession phs002812.v1.p1. Genetic maps for hg19 were downloaded from Broad Institute (https://alkesgroup.broadinstitute.org/Eagle/downloads/tables). PrivaSeq was downloaded and installed from http://privaseq.gersteinlab.org. scPrivacy scripts were downloaded from https://github.com/G2Lab/scPrivacy. Gene annotations files, gencode.v19, were downloaded from https://www.gencodegenes.org.

### Software resources

scDesignPop v0.0.0.9: https://github.com/chrisycd/scDesignPop

bcftools v1.20: https://github.com/samtools/bcftools [accessed July 8, 2024]

PLINK v1.90_beta 7.2: https://www.cog-genomics.org/plink/ [accessed July 26, 2024]

HAPGEN2 v2.2.0: https://mathgen.stats.ox.ac.uk/genetics_software/hapgen/hapgen2.html [accessed November 12, 2024]

FastQTL: https://github.com/francois-a/fastqtl [accessed March 7, 2025]

SAIGE-QTL: https://github.com/weizhou0/SAIGEQTL [accessed February 10, 2025]

jaxQTL: https://github.com/mancusolab/jaxqtl [acccessed June 1, 2025]

PrivaSeq: http://privaseq.gersteinlab.org [accessed December 3, 2024]

scPrivacy: https://github.com/G2Lab/scPrivacy [accessed December 2, 2024]

precrec R package v0.14.4: https://github.com/evalclass/precrec

Seurat R package v4.4.0: https://github.com/satijalab/seurat

SeuratDisk R package v0.0.0.9021: https://github.com/mojaveazure/seurat-disk

scDesign3 R package v0.99.7: https://github.com/SONGDONGYUAN1994/scDesign3

qvalue R package v2.30.0: https://github.com/StoreyLab/qvalue

ggplot2 R package v3.5.1: https://github.com/tidyverse/ggplot2

ggrastr R package v1.0.2: https://github.com/VPetukhov/ggrastr

ComplexHeatMap R package v2.14.0: https://github.com/jokergoo/ComplexHeatmap

gridExtra R package v2.3: 3https://github.com/cran/gridExtra/blob/master/R/gridExtra-package.r

CellMixS R package v1.14.0: https://github.com/almutlue/CellMixS

irlba R package v2.3.5.1: https://github.com/bwlewis/irlba

coop R package v0.6-3: https://github.com/wrathematics/coop

Rfast R package v2.1.0: https://github.com/RfastOfficial/Rfast

FactoMineR R package v2.11: http://factominer.free.fr/index.html

glmmTMB R package v1.1.9: https://github.com/glmmTMB/glmmTMB

MASS R package v7.3-58.2: https://cran.r-project.org/web/packages/MASS/index.html

MGLM R package v0.2.1: https://cran.r-project.org/web/packages/MGLM/index.html

matrixStats R package v1.3.0: https://github.com/HenrikBengtsson/matrixStats

splatter R package v1.22.1: https://github.com/Oshlack/splatter

powerEQTL R package v0.3.4: https://cran.r-project.org/src/contrib/Archive/powerEQTL/

scPower R package v1.0.4: https://github.com/heiniglab/scPower

phateR R package v1.0.7: https://github.com/KrishnaswamyLab/phateR

PHATE R package v1.0.11: https://github.com/KrishnaswamyLab/PHATE

## 5 Code availability

The scDesignPop R package is available on GitHub at https://github.com/chrisycd/scDesignPop with completed tutorials at https://chrisycd.github.io/scDesignPop/docs/index.html. The code used to reproduce our analysis is available at https://github.com/chrisycd/scEQTLsim.

## 6 Acknowledgements

We thank the group members of the Junction of Statistics and Biology at UCLA (http://jsb.ucla.edu), and the anonymous reviewers from Research in Computational Biology (RECOMB) for their comments and feedback. Additionally, we thank Dr. Powell and their lab for providing the OneK1K data.

## 10 Methods

### 10.1 scDesignPop’s generative modeling framework

#### 10.1.1 Modeling notations

Let *i* = 1, …, *I* index cells, *j* = 1, …,*J* genes, *k* = 1, …, *K* individuals (samples), *l* = 1, …, *L* SNPs, and *s* = 1, …, *S* cell states. scDesignPop is trained using the following five matrices:

- **Y** = [*Y*_*i j*_] ∈ ℝ^*I*×*J*^ : a cell-by-gene expression matrix with *I* cells and*J* genes. For scRNA-seq data, **Y** is typically a non-negative count matrix (i.e., 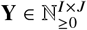). For example, in 10x Genomics scRNA-seq data, *Y*_*i j*_ represents the UMI-count of gene *j* in cell *i*. On the other hand, **Y** ∈ ℝ^*I*×*J*^ when normalized scRNA-seq data is used.
- **G** = [*G*_*kl*_] ∈ {0, 1, 2}^*K*×*L*^ : an individual-by-SNP genotype matrix with *K* individuals and *L* SNPs. Here, *G*_*kl*_ denotes the genotype dosage of the number of alternative alleles for SNP *l* in individual *k*.
- **Z** = [**z**_1_, …, **z**_*I*_]^T^ ∈ {0, 1}^*I*×*K*^ : a cell-by-individual indicator matrix with *I* cells and *K* individuals. Each vector **z**_*i*_ encodes cell *i*’s membership to individual *k* (i.e., **z**_*i*_ is one-hot encoded, with 𝓏_*ik*_ := **1**{*i* ∈ *k* }). This information is obtained by mapping each cell’s barcode to an individual via demultiplexing.
- **S** = [**s**_1_, …, **s**_*I*_]^T^ ∈ ℝ^*I*×*S*^ : a cell-state matrix with *I* cells and *S* dimensions representing either discrete cell types or continuous cell states. If cells are annotated as belonging to *H* discrete cell types, then we set *S* = *H* − 1 and treat cell type as a categorical covariate one-hot encoded as **s**_*i*_ ∈ {0, 1}^*H*−1^ using *H* − 1 dummy variables (with one cell type as the baseline level); if instead cell states are continuous (e.g., pseudotime), then **s**_*i*_ ∈ ℝ^*S*^ is a continuous vector representing *S* lineages.
- **C** = [**x a b e**] = [**c**_1_, · · ·, **c**_*I*_]^T^ ∈ ℝ^*I*×q^ : an (optional) cell-by-covariate design matrix with *I* cells and q total dimensions corresponding to individual-level covariates, cell-level covariates, batch covariate, and experiment condition covariate. The individual-level covariates **x** = [**x**_1_, …, **x**_*I*_]^T^ (i.e., age, gender, genotype principal components), are mapped to cells using **Z**, so cells from the same individual share identical individual-level covariates. The cell-level covariates **a** = [**a**_1_, …, **a**_*I*_]^T^ can correspond to mitochondrial percentage, sequencing pool, etc. The batch covariate **b** = [**b**_1_, …, **b**_*I*_]^T^ is one-hot encoded as **b**_*i*_ ∈ {0, 1}^*B*−1^ using *B* − 1 dummy variables (with one batch as the baseline level), if there are *B* total batches. Similarly, experiment condition covariate **e** = [**e**_1_, …, **e**_*I*_]^T^ is one-hot encoded as **e**_*i*_ ∈ {0, 1}^*E*−1^ using *E* − 1 dummy variables (with one condition as the baseline level), if there are *E* conditions. We refer to vector **c**_*i*_ as the *design covariates* for cell *i* henceforth.

#### 10.1.2 Modeling each gene’s marginal distribution

For each gene *j*, scDesignPop fits a generalized linear mixed model (GLMM) to flexibly capture the relationship between the response variable, whose value is a gene’s expression value in a cell, and its predictors. The predictors include the genotypes of putative eQTLs, a cell-individual indicator, a cell-type/state covariate, and additional cell- and individual-level covariates. Additionally, a random intercept specific for each individual is included to model the inter-individual variability. The parametric family *F*_*j*_ (·) is chosen according to the data modality of the response variable. scDesignPop currently supports Gaussian, Poisson, NB, and Bernoulli distributions, whose forms and link functions are summarized in Supplementary Table S3. Optional individual-level covariates and genotype-covariate interactions can be included as fixed effects.

Let *Y*_*i j*_ denote the expression value of gene *j* in cell *i*. To model the inter-individual variability, we introduce a random intercept *u*_*k j*_ for individual *k* independently generated from a Gaussian distribution 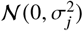 with zero mean and gene-specific variance 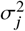. Conditional on *u*_*k j*_, the genotype vector **G**_*k,l*(*j*)_ for putative eQTLs (where *l* (*j*) denotes the putative eSNP indices for gene *j* in an individual-by-SNP genotype matrix **G**), the cell-individual indicator vector **z**_*i*_, the cell-state vector **s**_*i*_, and the cell design covariate vector **c**_*i*_, the marginal model for gene *j* is

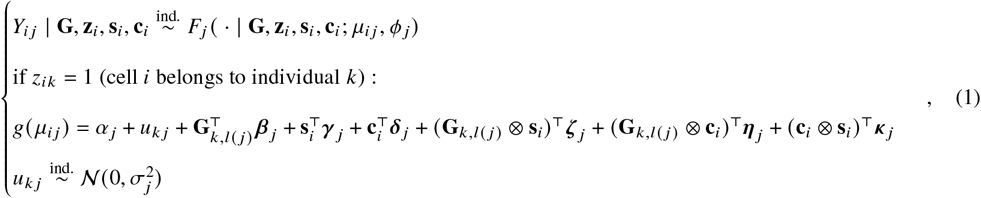

where *g*(·) denotes the link function, and **G**_*k,l*(*j*)_ ⊗ **s**_*i*_, **G**_*k,l*(*j*)_ ⊗ **c**_*i*_, and **c**_*i*_ ⊗ **s**_*i*_ are the genotypes-cell-state, genotypes-covariates, and cell-state-covariates interactions, respectively. The distribution *F*_*j*_ (·) is parameterized by location parameter 𝜇_*i j*_ and scale parameter 𝜙 _*j*_ . The model parameters estimated for each gene *j* include 𝜙 _*j*_, 𝛼 _*j*_, 𝜷 _*j*_, 𝜸 _*j*_, 𝜹 _*j*_, 𝜻 _*j*_, 𝜼 _*j*_, *κ* _*j*_, and 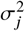. For model identifiability, given **s**_*i*_ is a categorical variable (e.g., cell type) and additional categorical variables in the design covariates **c**_*i*_ (such as batch or condition) are one-hot encoded, the first category of each variable is used as a baseline.

Besides the design covariate **c**_*i*_ that may be different between cells, scDesignPop explicitly models the genetic effects stemming from putative eSNPs using **G**_*k,l*(*j*)_ . Therefore, each gene has distinct eQTL genotype covariates, a key difference from scDesign3. Furthermore, each gene may have one or more putative eSNPs. If there is only one putative eSNP for a gene, then the genotype and its effect size parameter are scalars, that is, *G*_*k,l*(*j*)_ and 𝜷 _*j*_ . Motivated by findings from Yazar et al. and Natri et al. [5, 7], users may either specify a single SNP or multiple SNPs per gene using the `snp_mode` option (`“single”` or `“multi”`; Methods 10.2) for the putative eSNPs. If using the multi-SNP mode, the number of SNPs is automatically selected by scDesignPop using a correlation threshold (Methods 10.2).

We assume all cells are labeled with cell types or states and assigned to individuals. These assumptions are reasonable because they are naturally part of upstream data preprocessing via cell clustering/annotation and demultiplexing, respectively. Furthermore, we assume the scale parameter 𝜙 _*j*_ is gene-specific but constant across cells, and only the location parameter 𝜇_*i j*_ depends on eQTL, cell states, and design covariates. This balances model flexibility with computational tractability for population-scale datasets, where there may be millions of cells. Following the principle of model parsimony and to avoid overfitting, we assume there are only first-order interactions between each SNP and cell state pair in **G**_*k,l*(*j*)_ ⊗ **s**_*i*_, between each SNP and design covariate pair in **G**_*k,l*(*j*)_ ⊗ **c**_*i*_, and between each design covariate and cell state pair in **c**_*i*_ ⊗ **s**_*i*_. Additional assumptions are provided below.

##### Additional assumptions of the marginal model terms

- For the genotype vector **G**_*k,l*(*j*)_, we assume cells from the same individual have the same genotype. Also, eQTL effect sizes are assumed to be additive in the link function (e.g., log scale if *F*_*j*_ (·) is Poisson or NB). Under this assumption, the eQTL effect size is defined as a *per-allele eQTL effect size*.
- For the random intercept *u*_*k j*_ for individual *k* and gene *j*, we assume there are no dependencies between random effects *u*_*k j*_ among the *K* individuals. However, population structure of the underlying cohort and relatedness between individuals can be accounted for as fixed effects using genotype principal components (PCs) as covariates in the design matrix **C**. Lastly, we assume there are no interactions between the random effects and any fixed effects.

In practice, users may specify a simpler marginal model for Equation 1 depending on available design covariates, interaction terms, and input data size. The specific marginal models used by scDesignPop for the OneK1K and CLUES cohorts are described in Methods S1.

The marginal distribution *F*_*j*_ (· | **G, z**_*i*_, **s**_*i*_, **c**_*i*_) in Equation 1 is fitted using the R package `glmmTMB`. The fitted marginal distribution is denoted by 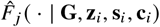.

#### 10.1.3 Modeling the joint distribution of genes

scDesignPop models the joint distribution of gene expression using a Gaussian copula framework, which allows flexible gene marginal distributions while capturing gene-gene rank dependence.

Let *y*_*i j*_ be a realization (observed expression) from a random variable *Y*_*i j*_ for gene *j* . We denote the joint cumulative distribution function (CDF) as *F*_*C*_ (· | **G, z**_*i*_, **s**_*i*_, **c**_*i*_), given the genotype matrix **G**, cell-individual membership **z**_*i*_, cell-state covariate **s**_*i*_, and design covariates **c**_*i*_ from the training data.

Under a copula, the joint distribution of*J* genes in cell *i* is constructed using the marginal CDFs *F*_1_(· | **G, z**_*i*_, **s**_*i*_, **c**_*i*_), …, *F*_*J*_ (· | **G, z**_*i*_, **s**_*i*_, **c**_*i*_). For *y*_*i j*_, we compute its marginal CDF value *w*_*i j*_ as

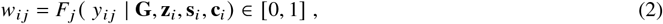

and transform it using the inverse standard Gaussian CDF to obtain 𝜓_*i j*_ = Φ^−1^(*w*_*i j*_). For a Gaussian copula, the transformed vector of marginal CDFs (𝜓_*i*1_, …, 𝜓_*iJ*_)^T^ follow a*J*-dimensional Gaussian distribution with zero mean and cell-state-specific covariance matrix 𝚺(**s**_*i*_), denoted as Φ_*J*_ (· ; 𝚺(**s**_*i*_)) such that [0, 1] ^*J*^ → [0, 1]. Because all transformed marginals are standard Gaussian with variances equal 1, 𝚺(**s**_*i*_) is equivalently a correlation matrix.

The resulting joint CDF of gene expression for cell *i* is

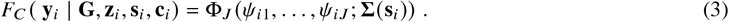

where **y**_*i*_ = (*y*_*i*1_, …, *y*_*iJ*_)^T^ are realizations from a random vector **Y**_*i*_ = (*Y*_*i*1_, …, *Y*_*iJ*_)^T^ for*J* genes.

The Gaussian copula is estimated using a plug-in estimate for the covariance matrix (i.e., the sample correlation matrix 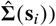 along with the fitted marginal distributions 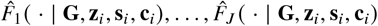 from Methods 10.1.2. When **s**_*i*_ encodes a discrete cell type, scDesignPop estimates 𝚺(**s**_*i*_) as the sample correlation matrix computed from all cells of that type. When **s**_*i*_ is continuous, scDesignPop estimates 𝚺(**s**_*i*_) by pooling cells with similar values of **s**. Also, when 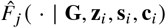 is a continuous distribution, each realization *y*_*i j*_ is transformed using Equation 2 as 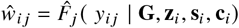 . However, when 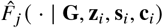 is a discrete distribution with support on non-negative integers (i.e., a NB distribution), we apply a distributional transformation as in [58, 86]:

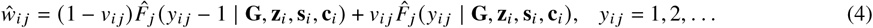

where *v*_*i j*_ ‘s are sampled independently from Uniform[0, 1] ∀ *i* = 1, …, *I*; *j* = 1, …,*J*.

Given 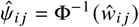, the sample correlation matrix 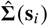 is estimated with 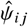 using the R package `matrixStats`.

Therefore, the estimated joint distribution is

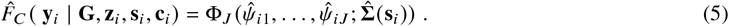

#### 10.1.4 Modeling the cell-type proportions

In addition to marginal and copula modeling, which enable generation of synthetic cells within the same individuals, scDesignPop incorporates cell-type proportion modeling as a third component to simulate cells for new individuals. Cell-type composition is modeled using a two-layer hierarchical framework, whereby the total number of cells per individual is first sampled, then cells are subsequently allocated to cell types via a multinomial distribution.

Formally, let there be *H* cell types (or discrete cell states) indexed by *h*. For each individual *k*, the total number of cells *n*_*k*,total_ follows a log-normal distribution (ℒ 𝒩) with mean 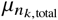 and variance 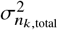as

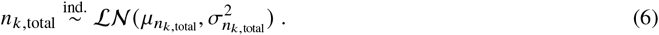

Conditional on *n*_*k*,total_, a vector of cell-type counts **n**_*k*_ = (*n*_*k*1_, …, *n*_*kH*_)^T^ is modeled by a multinomial distribution

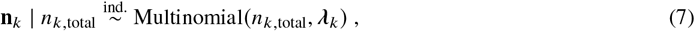

where 𝜆_*k*_ = (𝜆_*k*1_, …, 𝜆_*kH*_)^T^ is an individual-specific cell-type composition vector satisfying 0 < 𝜆_*kh*_ < 1 and 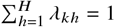.

To allow cell-type composiTo allow cell-type composition to vary across individuals, we link 𝜆_*k*_ to optional individual-level covariates. Let **X** = [**x**_1_, …, **x**_*K*_]^T^ ∈ ℝ^*K*×*p*^denote a *K* × *p*individual-level design matrix that is a subset of the **C** cell-level design matrix (Methods 10.1.1). Covariates **x**_*k*_ may comprise of genotype principal components, ancestry, disease status, sex, age, or other individual-level predictors. Note that we previously used **x**_*i*_ to denote the individual-level covariates of cell *i*, given the cell-to-individual mapping indicator **z**_*i*_. Here, we use the individual index *k* for notation convenience, so **x**_*k*_ = **x**_*i*_ if cell *i* belongs to individual *k* (i.e., 𝓏_*ik*_ = 1). Then, the composition vector 𝜆_*k*_ is linked to individual-level covariates **x**_*k*_ via multinomial logit regression as

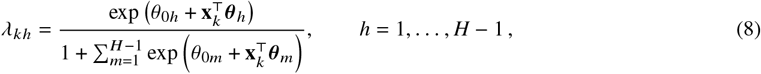

where 𝜃_0*h*_ and 𝜽 _*h*_ ∈ ℝ^*p*^are the intercept and slope coefficients for cell type *h*, respectively. To satisfy the coefficients 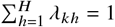, there are *H* − 1 sets of estimable regression coefficients, each of length *p*+ 1. To simplify notations, let 𝚯 ∈ ℝ^(*p*+1)×(*H*−1)^ be a parameter matrix collecting these sets of regression coefficients.

scDesignPop fits the univariate log-normal distribution using the R package `MASS`, and fits the multinomial regression model using the R package `MGLM`.

#### 10.1.5 Synthetic data generation

scDesignPop generates a synthetic cell-by-gene expression matrix 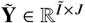 for Ĩ synthetic cells in 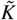 individuals using the same*J* genes and corresponding putative eSNPs as in the training data, given the fitted marginal models, Gaussian copula model, and cell-type proportion model from Methods 10.1.2, 10.1.3, 10.1.4.

Users may provide or simulate the following inputs for the synthetic cohort: an individual-level design matrix 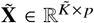, genotype matrix 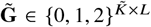, cell-level design matrix 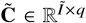, cell-state matrix 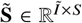, and cell-individual indicator matrix 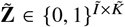 . These matrices may be (1) user-specified (e.g., a held-out cohort), (2) resampled from the training data, or (3) externally simulated [85]. When simulating new individuals without pre-specified cell labels, 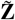 and 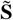 are generated using the fitted cell-type proportion model described below. For 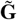, users can use existing genotype simulators, such as HAPGEN2, HAPNEST, and others [66, 83–85]); for 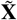 and 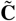, generative models are needed to simulate individual-level and cell-level design covariates, respectively.

The synthetic data generation proceeds in three steps as follows.

##### Step 1. Generating cell numbers and cell-type composition

For each new individual 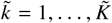, the total number of cells is sampled from the fitted log-normal distribution 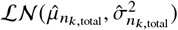 from Equation 6. Because the distribution is continuous, whereas total cell count must be discrete, we set the sampled value to the nearest integer, denoted as 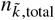. This discretization has a negligible impact in practice, since population-scale scRNA-seq studies typically sequence hundreds to thousands of cells per individual.

Then, conditional on the individual-level covariates 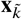 and 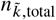, the vector of cell type counts 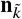 is sampled independently from the fitted multinomial logit model

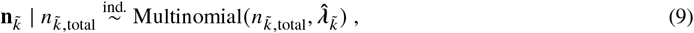

where 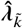 is computed using the estimated regression coefficients 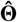 from Equation 8.

Thus, the sampled counts 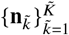 determine the synthetic cell-state matrix 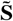 and cell-individual indicator matrix 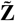. If synthetic cells are instead generated for the same *K* training individuals with provided cell type labels, then this step is omitted.

##### Step 2: Generating individual-specific random effects and copula variables

For each gene *j* and individual 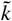, the individual-specific random intercept is generated as

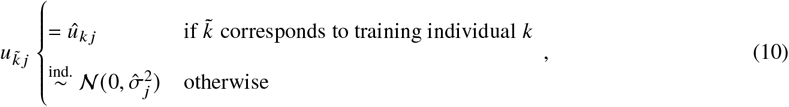

where 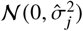 is the fitted Gaussian from Equation 1.

Then, for each synthetic cell 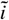 belonging to individual 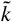 with cell-state 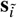, we sample a vector of latent values in ℝ^*J*^ from a multivariate Gaussian distribution (MVN),

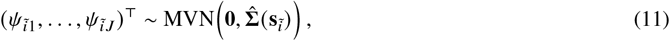

where 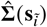 is the estimated cell-state-specific covariance matrix from Equation 5.

These latent values are transformed using the standard Gaussian CDF to obtain 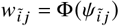 for *j* = 1, …,*J*, so that 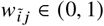 and the joint dependence structure is preserved through the Gaussian copula.

##### Step 3: Generating gene expression values

For each synthetic cell 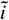 from individual 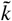, the conditional mean of gene *j*, denoted 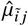, is computed using the fitted GLMM from Equation 1, individual-specific intercept 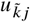, individual’s eQTL genotypes 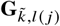, cell-indicator 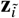, cell-state covariate 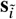, and design covariates 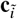

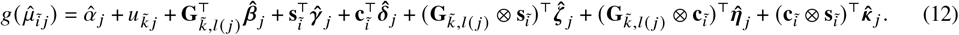

Finally, given the estimated conditional mean 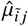, the estimated dispersion 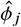, and CDF value 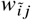, the gene expression vector for synthetic cell 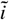, denoted as 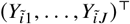, is obtained by its inverse CDF value from the fitted marginal parametric family

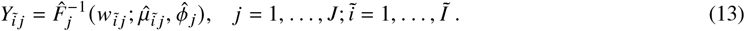

Together, these three generative components enable scDesignPop to have interpretable parameters and modeling flexibility, thus enabling several applications.

### 10.2 Multi-SNP and single-SNP eQTL modeling

scDesignPop has the option to either use a single-SNP mode (`snp_mode = “single”`) or a multi-SNP mode (`snp_mode = “multi”`). The former assumes only one eQTL SNP across cell types per gene, while the latter assumes at least one eQTL SNP. Specifically, **G**_*k,l*(*j*)_ is a scalar in Equation 1 when using single-SNP mode, and a vector when using multi-SNP mode. Depending on the data size and desired application, users may opt to use a single-SNP marginal model per gene for a simpler marginal model.

On the other hand, scDesignPop can also model multiple SNPs in the marginal model (Equation 1) since multiple population-scale studies have revealed cell-type-specific eQTLs, where there are different putative eQTLs in different cell types [5, 6]. However, SNPs at the loci of these putative eQTLs may be highly correlated with one another (i.e., in linkage disequilibrium). To ensure the marginal model remains tractable, scDesignPop prunes these SNPs based on their estimated pairwise Pearson correlations of the SNPs’ genotype for a given gene across all individuals. SNP pruning has been employed in popular genomic analysis tools such as PLINK [87]. A correlation threshold of 0.9 was chosen as a sensible default value when using the multi-SNP mode in our analyses. scDesignPop also filters out SNPs whose genotypes have zero variance (e.g., all 0’s), which translates to filtering out SNPs with 0 minor allele frequency (MAF).

### 10.3 Data preprocessing

The scRNA-seq, genotype, eQTL data, and gene selection from OneK1K and CLUES cohorts were each preprocessed accordingly before being modeled using scDesignPop.

#### Cell type filtering and genotype preprocessing

For the OneK1K dataset, we first filtered out erythrocytes and platelets as done so by Yazar et al. [5], resulting in 1,267,768 remaining cells from 981 individuals spanning 14 cell types. Then, we subsampled 100 individuals and split the cells into 70:30 training and testing sets by: 1) each cell type for the 100 individuals, and 2) individuals into 70 and 30 individuals, respectively.

For the CLUES dataset, we focused on Asian and European ancestries as done so by Perez et al. [6], and because these subpopulations had adequate individuals with healthy and diseased states. Next, we filtered out plasmablasts (“Pb”), proliferating lymphocytes (“Prolif”), and CD34+ progenitors (“Progen”), as well as individuals whose genotypes were greater than 95% missing, resulting in 1,232,696 remaining cells from 256 individuals spanning 8 cell types. Then, we subsampled 150 individuals and split the cells into 70:30 training and testing sets either by: 1) each cell type within the 150 individuals, and 2) across individuals into 105 and 45 individuals, respectively.

#### eQTL preprocessing

For OneK1K cohort we constructed the eQTL-genotype input data for scDesignPop as follows:

1. We first merged significant (≤ 0.05 FDR) cell-type-specific eQTL results from https://onek1k.org with the Matrix eQTL’s *cis*-eQTL results for all cell-types containing nominal *p*-value, slope (beta), *t*-statistic, and adjusted FDR values by matching the SNP id, gene, cell type and chromosome to obtain eQTL annotations. eQTLs with missing slope values were filtered out.
2. Next, we filtered for the first putative eSNPs found in each gene and each cell-type corresponding to `ROUND = 1` as found by Yazar et al. [5].
3. Then, we used the distinct SNP loci to extract the genotype data for selected individuals from the VCF files using `bcftools` [88], filtering out multi-allelic and non-SNP variants.
4. Finally, cell-type-specific eQTL annotations were merged with extracted genotype data after converting them to 0, 1, or 2 as the number of alternative alleles. SNPs with missing genotypes were filtered out. These eQTL annotations combined with eQTL genotype data (eQTL-genotype) were used as input in scDesignPop.
5. Steps 3 − 4 were conducted separately for training and testing data based on respective individuals’ sample ids.

For CLUES cohort we constructed the eQTL-genotype input data for scDesignPop as follows:

1. We first merged cell-type-specific eQTLs and cell-type specific eGenes from Supplementary Table S6 and Table S7 in Perez et al. [6] and retained cell type, gene-snp pair, nominal *p*-values, slope, adjusted *p*-value, and FDR results to obtain eQTL annotations. Distinct cell-type-specific eQTLs were kept and eQTLs corresponding to `“pbmc”` cell types were filtered out, since these correspond to eQTLs across all cell types.
2. Next, we filtered the eQTL annotations using an FDR threshold of 0.40.
3. Then, we used the distinct SNP loci obtained from the gene-SNP pairs to extract the genotype data for selected individuals from the VCF files using `bcftools`, filtering out multi-allelic and non-SNP variants.
4. Finally, cell-type-specific eQTL annotations were merged with extracted genotype data after converting them to 0, 1, or 2 as the number alternative alleles. SNPs with missing genotypes were filtered out. These eQTL annotations combined with eQTL genotype data (eQTL-genotype) was used as input in scDesignPop.
5. Steps 3 − 4 were conducted separately for training and testing data based on respective individuals’ sample ids.

#### Gene selection

For OneK1K cohort, we used R package Seurat (v4.4.0) to find the top 2000 highly variable genes (HVG). Using the scRNA-seq training data stratified by cell type in **Cell type filtering and genotype preprocessing**, we first normalized with default normalization settings `normalization.method = “LogNormalize”` and `scale.factor = 10000`, before using `FindVariableFeatures` with `selection.method = “vst”`. Next, to ensure consistency with downstream eQTL modeling, we found the intersection of genes with those in the eQTL-genotype data detailed in **eQTL preprocessing** (Methods 10.3), resulting in 817 HVGs used for marginal modeling. Applying the same procedure to the training data stratified by individuals yielded 810 overlapping HVGs.

For CLUES cohort, normalization was not performed since the provided scRNA-seq data had already been batch-corrected and normalized. We followed the same Seurat workflow as for OneK1K; however, since there were only 1999 genes in the scRNA-seq data, all genes were retained. After intersecting with the eQTL-genotype data detailed in **eQTL preprocessing** (Methods 10.3), 798 HVGs remained and were used for marginal modeling in both the individual-stratified and cell-type-stratified training data.

### 10.4 Simulator comparison

splatPop [48] is the only existing population-scale scRNA-seq simulator that incorporates eQTL effects to the best of our knowledge. For this reason, we did not compare with other multi-individual scRNA-seq simulators such as muscat or rescueSim [46, 47].

#### splatPop simulation

We compared scDesignPop’s performance to splatPop by following their tutorial to simulate the test data directly. In brief, we used splatPop’s `splatPopEstimate(), setParams(), splatPopSimulateMeans()`, and `splatPopSimulateSC()` functions from the R package `splatter` (v1.22.1) with genotype, gene annotation, eQTL and scRNA-seq count data as input to simulate the test data for OneK1K cohort. Since splatPop requires eQTL slopes, we used the Matrix eQTL *cis*-eQTL results for each cell-type provided by the OneK1K authors (Methods 10.3). Detailed steps are outlined below.

1. First, we extracted relevant gene annotations from GENCODE v19 gtf file [89] and filtered for the same highly variable genes as in the training data.
2. Second, we computed the individual-level means for the training data by computing the mean of counts for each cell type and individual.
3. Next, we subsetted the scRNA-seq training data, the scRNA-seq testing data, gene annotation, and eQTL data into each cell type. We also filtered the VCF file for individuals in the training data, testing data, and for the SNPs corresponding to the eQTL data.
4. Then, we used splatter’s `splatPopEstimate` to estimate the individual-level means with the scRNA-seq count matrix and eQTL data as input for each cell type.
5. We used `setParams` to use the estimated individual-level means and set `eqtl.maf.min = 0`.
6. Next, we simulated the means by using `splatPopSimulateMeans`, with individual-filtered VCF, gene annotation, eQTL, and individual-level means for each cell type.
7. Next, we used `splatPopSimulateSC` with simulated individual-level means and estimated parameters from Step 6 to simulate for each cell type in the test data. We set the `batchCells` parameter equal to the number of cells of an individual and cell type in the test data.
8. Finally, we merged the simulated files from all cell types together and converted this into a `SingleCellExperiment` object.

Since splatPop [48] is restricted to count-based scRNA-seq data as input, we were unable to use splatPop to simulate normalized data in the CLUES cohort.

#### scDesignPop simulation

For scDesignPop, we used two different modes (single-SNP and multi-SNP modes), denoted as scDesignPop_single and scDesignPop_multi (e.g., in Figs 2, 3), respectively. The primary difference between these two is that the latter assumes ≥ 1 eQTL per gene while the former assumes only 1 eQTL per gene. Details of terms assumed in the models used to simulate OneK1K and CLUES cohort data are in Methods S1.

### 10.5 Evaluation metrics

We evaluated scDesignPop using both qualitative and quantitative metrics. For qualitative metrics, we visualized the simulated data on a 2D low-dimensional embedding using a UMAP plot [62], the gene-gene correlation matrices with heatmaps [90], and the pseudobulk expression across genotypes of a putative eQTL for a gene. For quantitative metrics, we used the mean local inverse Simpson’s index (mLISI) of cells [91] to compare the reference test data and each simulated data on the 2D UMAP plot, as previously done so in [58, 86]. We used the RV correlation [92] to compare the correlation matrices. We evaluated the similarities between test and simulated scRNA-seq data by computing twelve summary statistics spanning gene-, cell-, and pseudobulk-levels and visualized their distributions using violin plots and compared with the Kolmogorov-Smirnov test statistic.

#### UMAP visualization

Overplotting can be an issue when visualizing a large number of cells in an overall UMAP plot. Thus, we also generated finer-scaled UMAP plots by further grouping cells by the individuals, by the genotypes of an eQTL, or by a combination of design covariates if there are any.

#### Gene correlation matrices

Gene-gene correlation matrices were also used to qualitatively compare the gene expression between genes. For this, we first normalized the data using Seurat, then subsetted the data by specified cell type (if any) and then found the top 100 expressed genes by ranking each gene’s average expression, then computed the Pearson correlation using using R package `coop` (v0.6-3). Hierarchical clustering with complete linkage was applied on 1 – *r* values (where *r* is the sample Pearson correlation) for the test data, and then the same gene ordering was applied to the simulated data. We visualized the correlation matrices using R packages `ComplexHeatMap` (v2.14.0) and `gridExtra` (v2.3).

#### RV correlation

We computed the RV correlation [92] using R package `FactoMineR` (v2.11) as a metric to quantify how close the computed gene-gene correlation matrix of the simulated data resembles that of the test data, denoted as 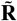 and **R**, respectively:

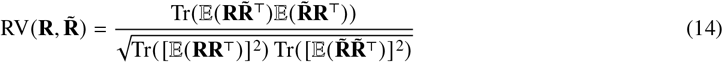

#### Gene expression heatmaps

To show the expression pattern across individual-level covariates (e.g., age, sex, ancestry, and disease status), we extracted the top 100 HVGs using Seurat’s default “vst” method, and then calculated the pseudobulk expression of these genes for each individual by summing the log-normalized (log1p()) counts. The pseudobulk expression values were then scaled within each individual to better visualize the expression pattern in the heatmap with group labels (e.g., ancestry and disease status)

#### Spearman correlations of cts-eQTLs

First we normalized the count data using log1p. Then, we obtained the pseudobulk gene expression for individuals per cell type by averaging the expression for each gene. Spearman correlations between genotypes and pseudobulk expression levels of genes for putative eQTLs were then estimated within each cell type. By comparing these Spearman correlations of the same putative eQTLs between the test and synthetic data, we evaluated the eQTL effects preserved in the synthetic data.

#### mLISI

We used the R package `CellMixS` (v1.14.0) to compute the mean local inverse Simpson’s index (mLISI) as a metric to measure the similarity between the test and synthetic data on 2D UMAP plot. Here, an mLISI value closer to 2 implies that cells between the test data and synthetic data are well mixed. Whereas, an mLISI value of 1 means that the two groups are more distinct.

#### Summary statistics

Twelve summary statistics spanning across gene-level, cell-level, and pseudobulk-level were computed and their distributions were compared. A two-sample Kolmogorov-Smirnov test was computed using `ks.test()` function in R. Pearson correlations were computed using R package coop (v0.6-3). Principal component analysis (PCA) was performed using R package `irlba` (v2.3.5.1) on centered and scaled data. Unless indicated otherwise, we normalized the counts for OneK1K data using default Seurat settings, and we directly used the normalized test and simulated data for CLUES data without any further normalization. To compute the pseudobulk matrix (gene-by-individual), we summed all the counts across cells within an individual for OneK1K data. Unless indicated otherwise, we normalized the pseudobulk counts for OneK1K data using log(pseudob. expr. + 1), and we directly used the pseudobulk expression for CLUES data without any further normalization.

##### Gene-level summary statistics

1. Mean of normalized gene expression: a gene’s mean of normalized expression across all cells.
2. Variance normalized gene expression: a gene’s variance of normalized expression across all cells.
3. Gene non-zero proportion: a gene’s proportion of nonzero counts across all cells.
4. Gene-gene correlation: the Pearson correlations between two genes’ normalized expression across all cells.

##### Cell-level summary statistics

1. Cell non-zero proportion: a cell’s proportion of nonzero counts across all genes.
2. Cell library size: a cell’s total counts across all genes (log scale for OneK1K data, 10^−4^ scale for CLUES data).
3. Cell-cell correlation: the Pearson correlations between two cells’ normalized expression across all genes.
4. Cell-cell distance: the Euclidean distance between two cells in the 50-dimensional principal component space.

##### Pseudobulk-level summary statistics

1. Mean of normalized pseudobulk gene expression: a gene’s mean of log(pseudob. expr. + 1) across all individuals (no log transformation for CLUES data).
2. Variance of normalized pseudobulk gene expression: a gene’s variance of log(pseudob. expr. + 1) across all individuals (no log transformation for CLUES data).
3. Individual library size: an individual’s pseudobulk expression across all genes (log scale for OneK1K data, 10^−2^ scale for CLUES data).
4. Individual-individual distance: the Euclidean distance between two individuals in the 30-dimensional principal component space.

For cell-cell correlations, we first normalized the test data using Seurat’s default settings (`normalization.method = “LogNormalize”` and `scale.factor = 10000`), then randomly sampled 5,000 cells and computed their pair-wise correlations using R package coop (v0.6-3). For cell-cell distance, we first normalized the test data using Seurat’s default settings, then found the top 50 PCS on the centered and scaled data using R package `irlba` (v2.3.5.1). Next, we normalized the simulated data and projected all cells onto the same PC space using `predict()` function. Lastly, we randomly sampled 5,000 cells and computed every pairwise Euclidean distance using R package `Rfast` (v2.1.0) with `method = “euclidean”` option. For CLUES data, we skipped data normalization since the data representation is already normalized.

For individual-individual distance, we first normalized the pseudobulk expression matrix using `log1p()`, then found the top 30 PCS on the centered, scaled, and normalized test data using R package `irlba` (v2.3.5.1). Next, we projected the centered, scaled, and normalized simulated data onto the same PC space using `predict()` function, and computed every pairwise Euclidean distance using R package `Rfast` (v2.1.0) with `method = “euclidean”` option. For CLUES data, we skipped data normalization since the data representation is already normalized.

Non-zero proportion of genes and non-zero proportion of cells were not plotted for the CLUES data (Fig. S2) since the expression are normalized and thus not informative as summary statistics.

### 10.6 Generating synthetic genotypes for new individuals

To simulate scRNA-seq data for new individuals not present in scDesignPop’s training data, genotypes at putative eQTL loci are required. scDesignPop can incorporate synthetic genotypes generated by tools such as HAPGEN2 [66], a reference-based genotype simulator.

We simulated genotypes for 982 synthetic individuals across all autosomes using HAPGEN2 (v2.2.0) with phased genotypes from the OneK1K cohort as the reference panel. HAPGEN2 was selected because it preserves the linkage disequilibrium (LD) structure of the underlying cohort [85], and scales to genome-wide data with millions of SNPs. All file processing and format conversions were performed using `bcftools` (v1.2.0). The synthetic genotypes were used in simulating scRNA-seq data for new individuals (Results 2.4) and in eQTL-based linking attacks (Methods 10.11).

Genotypes were simulated in three steps, processed per chromosome: 1) conversion of variant call format (VCF) files to IMPUTE2 format, 2) haplotype simulation with HAPGEN2, and 3) conversion back to VCF format with variant annotation. First, phased OneK1K genotypes in VCF format were converted to IMPUTE2 format using `bcftools convert –haplegendsample`, producing triplet files (`.haps, .leg, .samp`) for each chromosome. Because the OneK1K genotypes were already phased (i.e., 0|0, 0|1, 0|1, 1|1), no additional phasing was required. Genetic maps, required by HAPGEN2, containing the recombination frequencies for hg19 build were obtained from the Broad Institute, split into files for per chromosome, and supplied to HAPGEN2.

Second, we simulated phased haplotypes for 982 synthetic individuals per chromosome using HAPGEN2 with the -n 982 0 argument. Since HAPGEN2 required specification of a disease locus even when simulating a synthetic cohort that mimics the reference, we selected the genomic position of the first variant as a placeholder SNP, denoted as `DUMMY_SNP`, and supplied it using the `-dl DUMMY_SNP 0 0 0` argument.

Finally, simulated haplotype outputs were converted back to VCF format using `bcftools convert –haplegendsample2vcf`. Variant-level metadata (ID, QUAL, FILTER, and INFO/AF fields) were transferred from the original OneK1K VCF files to the simulated VCFs or annotated accordingly using `bcftools annotate`.

### 10.7 Dynamic eQTL effect modeling

To prepare the data for modeling, we extracted count data for the 12,420 B cells (immature/naive B cells (bin) and memory B cells (bmem)) from 100 individuals from the OneK1K cohort [5]. From these individuals, 817 highly variable genes (HVGs) were selected following Methods 10.3. We then applied PHATE for dimensionality reduction using the `phateR` (v1.0.7) interface (PHATE (v1.0.11) in Python). Pseudotime was inferred on the reference data using `slingshot` (v2.7.0). The dataset was then randomly partitioned into a 70% training set and a 30% test set at the individual level. The training set was used for model fitting and data simulation, and the resulting simulated data were evaluated by comparison with the held-out test set. For visualization and comparative analysis, PHATE dimensionality reduction was applied independently to both the simulated data and the held-out test set.

Consistent with Methods 10.1.2, a negative binomial (NB) mixed model was used as the parametric form in the gene marginal model for dynamic eQTL effects, using pseudotime to represent the continuous cell-states *s*_*i*_ for each cell *i*. An interaction effect between cell-state (pseudotime) and eQTL effects was included. Here, *l* (*j*) represents the index of the single SNP that is used for each gene *j* .

To model linear dynamic eQTL effects, we used

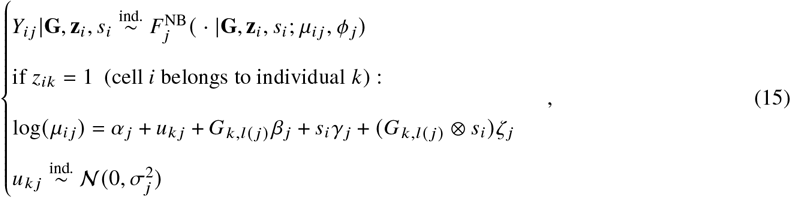

where we only used continuous cell states (pseudotime) to replace both the effect from the discrete cell types and their corresponding interaction effect with eQTL effects in the gene marginal model from Methods 10.1.2.

To model nonlinear dynamic eQTL effects, we used

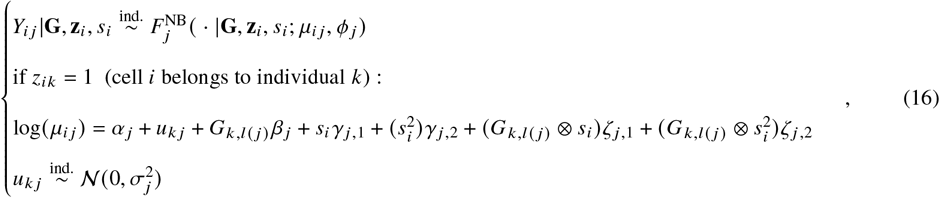

where we included both the linear and quadratic terms of the cell states (pseudotime) in the gene marginal model, including their corresponding interaction effects with eQTL effects.

For both scenarios, we then followed the same joint gene modeling and synthetic data generation procedure to generate simulated data.

When further visualizing the data in smooth curves and violin plots, the count data was log-normalized first (denoted 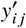, using natural log), then scaled as 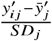 in each gene *j* to get the Z-scores. The smoothing was performed by `ggplot2 default in geom_smooth()` with a cubic-spline GAM (Generalized Additive Model).

### 10.8 Power analysis functionality

Since scDesignPop generates synthetic data that closely matches real data with realistic cts-eQTL effect sizes, scDesignPop can perform simulation-based power analysis by treating its estimated cts-eQTL effects as ground truth. For each gene, scDesignPop fits a marginal model containing the eQTL term (the alternative hypothesis) and creates a null model with the eQTL term removed. Then scDesignPop generates paired alternative (nonzero-effect) and null (zero-effect) datasets containing individuals and cells. For power analysis, an eQTL mapping model is applied to each simulated dataset to re-estimate the effect size. Currently, four model options are included: 1) negative binomial mixed (NB mixed) model, 2) Poisson mixed model, 3) linear mixed model, and 4) pseudobulk linear model (Methods S4). After *B* simulations, a critical threshold is determined from the null estimates (e.g., the (1 − 𝛼_adj_)th quantile for a one-sided 𝛼_adj_ test), and power is calculated as the proportion of alternative estimates that exceed (or fall below, depending on the sign) this threshold. Detailed steps are introduced in the Methods S3 and pseudocode provided in Supplementary Fig. S25.

#### Multiple testing correction

To control the family-wise error rate (FWER) due to multiple testing across genes and SNPs in eQTL analysis, we followed the approach used by Lonsdale et al. and Schmid et al. [15, 93]. Given *N*_gene_ genes and *N*_SNP_ independent (uncorrelated) SNPs tested per gene, the significance threshold 𝛼 will be corrected by 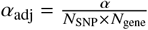.

### 10.9 Power analysis comparison

We compared scDesignPop’s power analysis performance to powerEQTL and scPower using the same classical monocytes from OneK1K dataset. Input parameters for powerEQTL and scPower were estimated empirically from the data. Power analysis with powerEQTL was conducted under the two modeling options provided by its package: a zero-inflated negative binomial (ZINB) mixed model for raw count data, and a linear mixed model for normalized expression data. In contrast, scPower performs power analysis using a simple linear model at the pseudobulk level to estimate eQTL effects.

For the ZINB mixed model in powerEQTL, required inputs include the slope, the number of individuals, the number of cells per individual, the mean and standard deviation of the individual random intercept, the dispersion parameter of the ZINB distribution, the probability that an excess zero occurs in the ZINB distribution, the minor allele frequency (MAF), the family-wise error rate (FWER) threshold, and the number of tests. Using the *glmmTMB* (v1.1.9) R package, we fitted a ZINB mixed model on the given data (classical monocyte in OneK1K), to obtain the slope and zero probability since these were needed as inputs for powerEQTL. The fixed intercept and the variance of the individual-level random intercept were estimated from the fitted model, with the latter derived from the standard deviation of the random intercept term. The dispersion parameter was estimated under the ZINB mixed model. Minor allele frequency (MAF) was calculated from the observed genotype data (classical monocytes from OneK1K), whereas the family-wise error rate (FWER) threshold and the total number of tests were fixed at 0.05 and 8,000, respectively.

For the linear mixed model in powerEQTL, required inputs include the slope, the number of individuals, the number of cells per individual, the standard deviation of the log-normalized counts, intra-class correlation (ICC), MAF, FWER threshold, and the number of tests. We fitted a linear mixed model using the *lme4* (v1.1-35.3) R package on the same given data (classical monocyte in OneK1K) to obtain the slope while the ICC was calculated based on the formula 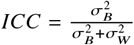, where 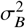 is the between-subject variance and 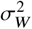 is the within-subject variance. Additionally, the standard deviation of the log-normalized counts was also calculated. Similarly, MAF was calculated based on the same given data (classical monocyte in OneK1K), while the FWER threshold and the number of tests were fixed as 0.05 and 8,000, respectively.

For scPower, required inputs include the R-squared value, and default values were used for other parameters. After log-normalizing the mean gene expression across cells for each individual (classical monocyte in OneK1K), we fitted a simple linear model between the log-normalized mean expression and genotype across individuals. The resulting coefficient of determination (*R*^2^) was supplied to scPower for power calculations across varying numbers of individuals and cells per individual. A significance threshold of 0.05 was supplied to scPower for Bonferroni correction, while the number of tests was internally determined and could not be specified by the user.

### 10.10 Benchmarking eQTL mapping methods

#### Simulating scRNA-seq data

We first trained scDesignPop on 93,376 cells from 70 subsampled individuals using the OneK1K cohort. For the marginal modeling, a negative binomial model with single-SNP mode was used, with genotype and cell type as main effects, and an interaction effect between cell type and genotype. For ease of parameter interpretability, no additional covariates were included in the marginal model. For ground truth, we used the cts-eGenes empirically found by Yazar et al. [5] under an FDR threshold ≤ 0.05 out of 810 highly variable genes as true cts-eGenes, with the remaining as true cts-non-eGenes. In CD4_NC_ T cells, there were 417 cts-eGenes and 393 cts-non-eGenes; in memory B cells, there were 156 cts-eGenes and 654 cts-non-eGenes.

Using scDesignPop, we generated approximately 1.2 million cells from 981 individuals under three simulation settings designed to progressively increase the separation between the eQTL genes (eGenes) and non-eGenes: (1) a `fitted` scenario using the effect sizes estimated from the reference data, (2) a trueH0 scenario in which non-eGenes were assigned zero eQTL effect sizes, and (3) a `trueH0_higher_eqtl` scenario where non-eGenes were assigned zero eQTL effect sizes and eGenes’ eQTL effects were amplified by 0.5 log2-fold-change relative to their fitted parameter estimates (Supplementary Table S5). For each simulation setting, we either used the fitted parameter estimates or modified the estimated cell-type main effects and interaction effects between cell-type and genotype in the marginal model via scDesignPop’s `ModifyMarginalModels`. Then, using the fitted (or modified) marginal and copula models, we simulated 1,267,768 cells for 981 individuals using the provided cell type labels and real genotypes from OneK1K cohort for 1 replicate under each simulation setting.

#### Data preprocessing

To ensure fair comparison of the eQTL mapping methods, we used biallelic SNPs from a 1 mega-basepair (Mbp) window flanking both ends of the gene body (i.e., its transcription start and end sites), and filtered for genetic variants with ≥ 0.05 MAF or ≥ 10 minor allele sample threshold. We used GRCh37 gene annotations from GENCODE v19 to prepare the input gene annotation files for each method. For all methods, we included age and sex as covariates.

#### FastQTL

To prepare the input data for FastQTL, we first normalized the scRNA-seq data by applying log(count + 1), then subsetted by the corresponding cell type, followed by aggregating the mean to obtain the pseudobulk expression for each individual. We ran FastQTL in adaptive permutation mode with 1000 and 100,000 as the lower and upper permutation limits (`–permute 1000 100000`) to obtain the beta-approximation adjusted *p*-values. Using FastQTL’s output, we then computed the gene-level q-values (using Storey’s procedure) from the adjusted *p*-values to call cts-eGenes or non-cts-eGenes.

#### SAIGE-QTL

For SAIGE-QTL, we perform *cis*-eQTL mapping following its three-step workflow. In step 1, for each gene and cell type from the synthetic scRNA-seq data, gene expression was modeled by SAIGE-QTL using a Poisson generalized linear mixed model as count data (`–traitType=count`) with age and sex included as fixed covariates (`–covarColList=sex,age`) and specified the individual IDs (`–sampleIDColinphenoFile=IND_ID`). In step 2, variant-level cis association testing was conducted using score tests, conditioning on the fitted null model and variance ratio estimates (`–GMMATmodelFile,–varianceRatioFile`) from step 1, with *p*-values calibrated using saddlepoint approximation (`–SPAcutoff=2 as default`) and without leave-one-chromosome-out (LOCO) correction (–LOCO=FALSE). Following data preprocessing, we used a minor allele frequency ≥ 0.05 (–minMAF=0.05) and minor allele count ≥ 10 (–minMAC=10) with predefined cis regions using gene annotations (–rangestoIncludeFile). The computed ACAT *p*-values were then corrected for multiple testing at the gene-level using Storey’s procedure. In step 3, variant-level *p*-values within each cis region were aggregated to obtain gene-level q-values using the Cauchy combination test to call cts-eGenes, as implemented in SAIGE-QTL.

#### jaxQTL

For jaxQTL, we considered both *p*-value calibration modes – ACAT-V test without permutation (jaxQTL_ACAT) and beta-approximation with permutations (jaxQTL_perm) – to correct for multiple testing due to many SNPs tested per gene. For the genotype data, we used PLINK to filter out biallelic SNPs with < 0.05 minor allele frequency or < 10 minor allele counts since jaxQTL does not currently have internal options and we wanted all methods to test for a comparable number of SNPs. For the simulated scRNA-seq data, we summed the synthetic counts within each cell type per individual to obtain the pseudobulk expression. Under both modes, we used default or recommended settings: negative binomial model (–model NB) with score method (`–test-method score`), and 2 expression PCs (–addpc 2), and –platform cpu. Age and sex were included as covariates. For the jaxQTL_perm mode (–mode cis), we used the default 1000 permutations (–nperm 1000).

#### Performance metrics

We ranked the performance of eQTL mapping methods across two cell types and three simulation settings based on their ability to detect cts-eGenes using five different metrics: 1) area under the receiver operating characteristic (AUROC) curve, 2) area under the precision-recall (AUPRC) curve, 3) Matthew’s correlation coefficient (MCC), 4) total computational time, and 5) peak memory usage.

For each method, we first computed the gene-level q-values using Storey’s ST procedure as implemented in the R package `qvalue` (v2.30.0). Within each cell type and simulation setting, we then compared the set of cts-eGenes identified by each method against the ground-truth cts-eGenes and cts-non-eGenes to categorize them as either true positives (TP), false positive (FP), true negatives (TN), or false negatives (FN). We used 1 − q-values as prediction scores to construct the ROC and PR curves and their areas under the curve using the R package `precrec` (v0.14.4). To account for class imbalance, we also computed the relative improvement over baseline AUPRC, defined as 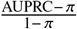, where 𝜋 denotes the proportion of true cts-eGenes in the given cell type.

We computed the Matthew’s correlation coefficient (MCC) [94] at a target false discovery rate (FDR) threshold of 5%, since this is commonly used to call cts-eGenes. MCC is less sensitive to class imbalance since it uses all four classification outcomes, and is defined as

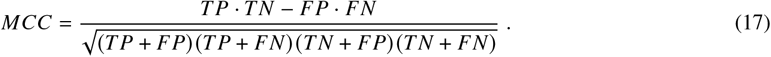

We also measured the peak memory usage in GiB and total computational time in CPU hours for each eQTL mapping method. This was done once under the “fitted” simulation setting in CD4_NC_ T cells by using `/usr/bin/time -v` for each method.

Finally, we ranked the methods within each simulation setting and cell type, then averaged them before assigning a final rank using `min_rank` from the R package `dplyr` (v1.1.4). For AUROC, AUPRC, and MCC metrics, higher values indicate better performance, while for peak memory usage and total computational time, lower values are better.

### 10.11 eQTL-based linking attacks

For the eQTL-based linking attack, we trained scDesignPop on 100 individuals from the OneK1K cohort and generated synthetic scRNA-seq data using either the original reference genotypes or synthetic genotypes simulated by HAPGEN2 for 981 individuals (including the training individuals). After aggregating the reference scRNA-seq or simulated scRNA-seq data into pseudobulk profiles for CD4_NC_ T cells (most common cell type), we followed PrivaSeq’s pipeline to predict genotypes. Specifically, adapting the create_pseudobulk.ipynb script from Walker et al. [13], raw count values were averaged per cell type and individual and normalized using log_2_(TMM + 1). Ten expression principal components (PCs) were regressed out from the normalized pseudobulk values to obtain the residuals. The residuals were transformed using a rank-based inverse normal transformation (RINT) to generate normalized pseudobulk gene expression as inputs for PrivaSeq’s linking attack [12]. In the linking attack, PrivaSeq identifies the first distance gap statistic for all 981 query individuals. The first distance gap statistic [12] is defined as the difference between the smallest and second smallest genotype distances (denoted as 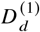 and 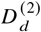, respectively) between query individual *d*’s genotype and all target individuals’ genotypes in the reference data:

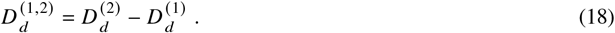

The linking attack results for each of the 981 query individuals were then classified as either predicted positive or predicted negative based on a pre-specified gap statistic threshold. As the threshold varied, precision was defined as the proportion of correctly linked individuals among those exceeding the threshold. The proportion of true positives was defined as the proportion of correctly linked individuals exceeding the threshold among all 981 links.

For visualization, gap statistics within each query individual group were compared using a two-sided Wilcoxon rank-sum test. Using the normalized pseudobulk gene expression matrices described above together with the corresponding genotype data, Spearman correlations were computed for all putative eQTL pairs within each query group. Reference Spearman correlations were obtained from the original OneK1K study. *R*^2^ values were then calculated between the estimated Spearman correlation for each query group and the same reference Spearman correlations from the original OneK1K study.

### 10.12 Computation time and peak memory

We assessed scDesignPop’s performance from loading input data to generating synthetic data using the OneK1K cohort. For training and data generation, we subsampled 20, 30, 50, 100, 150, 200 individuals with 25,904, 39,004, 66,319, 130,999, 194,124, 260,889 cells on average, respectively, for 810 highly variable genes. For each setting, we measured from repeated triplicates the peak memory in KiB (maximum resident set size), elapsed (wall clock) time in seconds, and percentage of CPU used with /usr/bin/time -v. The average peak memory was converted to GiB using 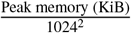 and the average total CPU hours were computed as 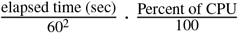.

scDesignPop currently supports four CPU parallelization backend options via the `parallel, BiocParallel`, `future.apply`, and `pbmcapply` R packages in marginal model fitting, copula fitting, parameter estimation, synthetic data generation, and power analysis modules. Parallelization was not implemented in cell type proportion modeling and simulation due to the relatively small number of observations for individual-level data.

All our analyses in this study were performed on a Linux server system (Ubuntu v22.04) equipped with two AMD EPYC 9754 CPUs, each with 128 cores, and 2304 GB of total RAM.

## References

[1] Martin Farrall. Quantitative genetic variation: a post-modern view. Human molecular genetics, 13(suppl_1): R1–R7, 2004.

[2] Anna SE Cuomo, Aparna Nathan, Soumya Raychaudhuri, Daniel G MacArthur, and Joseph E Powell. Single-cell genomics meets human genetics. Nature Reviews Genetics, 24(8):535–549, 2023.

[3] Maleeha Maria, Negar Pouyanfar, Tiit Örd, and Minna U Kaikkonen. The power of single-cell rna sequencing in eqtl discovery. Genes, 13(3):502, 2022.

[4] Jingfei Zhang and Hongyu Zhao. eqtl studies: from bulk tissues to single cells. Journal of Genetics and Genomics, 50(12):925–933, 2023.

[5] Seyhan Yazar, Jose Alquicira-Hernandez, Kristof Wing, Anne Senabouth, M Grace Gordon, Stacey Andersen, Qinyi Lu, Antonia Rowson, Thomas RP Taylor, Linda Clarke, et al. Single-cell eqtl mapping identifies cell type–specific genetic control of autoimmune disease. Science, 376(6589):eabf3041, 2022.

[6] Richard K Perez, M Grace Gordon, Meena Subramaniam, Min Cheol Kim, George C Hartoularos, Sasha Targ, Yang Sun, Anton Ogorodnikov, Raymund Bueno, Andrew Lu, et al. Single-cell rna-seq reveals cell type–specific molecular and genetic associations to lupus. Science, 376(6589):eabf1970, 2022.

[7] Heini M Natri, Christina B Del Azodi, Lance Peter, Chase J Taylor, Sagrika Chugh, Robert Kendle, Mei-i Chung, David K Flaherty, Brittany K Matlock, Carla L Calvi, et al. Cell-type-specific and disease-associated expression quantitative trait loci in the human lung. Nature Genetics, 56(4):595–604, 2024.

[8] Aparna Nathan, Samira Asgari, Kazuyoshi Ishigaki, Cristian Valencia, Tiffany Amariuta, Yang Luo, Jessica I Beynor, Yuriy Baglaenko, Sara Suliman, Alkes L Price, et al. Single-cell eqtl models reveal dynamic t cell state dependence of disease loci. Nature, 606(7912):120–128, 2022.

[9] Benjamin J Schmiedel, Divya Singh, Ariel Madrigal, Alan G Valdovino-Gonzalez, Brandie M White, Jose Zapardiel-Gonzalo, Brendan Ha, Gokmen Altay, Jason A Greenbaum, Graham McVicker, et al. Impact of genetic polymorphisms on human immune cell gene expression. Cell, 175(6):1701–1715, 2018.

[10] Julien Bryois, Daniela Calini, Will Macnair, Lynette Foo, Eduard Urich, Ward Ortmann, Victor Alejandro Iglesias, Suresh Selvaraj, Erik Nutma, Manuel Marzin, et al. Cell-type-specific cis-eqtls in eight human brain cell types identify novel risk genes for psychiatric and neurological disorders. Nature neuroscience, 25(8):1104–1112, 2022.

[11] Roy Oelen, Dylan H de Vries, Harm Brugge, M Grace Gordon, Martijn Vochteloo, single-cell eQTLGen consortium, BIOS Consortium, Chun J Ye, Harm-Jan Westra, Lude Franke, et al. Single-cell rna-sequencing of peripheral blood mononuclear cells reveals widespread, context-specific gene expression regulation upon pathogenic exposure. Nature communications, 13(1):3267, 2022.

[12] Arif Harmanci and Mark Gerstein. Quantification of private information leakage from phenotype-genotype data: linking attacks. Nature methods, 13(3):251–256, 2016.

[13] Conor R Walker, Xiaoting Li, Manav Chakravarthy, William Lounsbery-Scaife, Yoolim A Choi, Ritambhara Singh, and Gamze Gürsoy. Private information leakage from single-cell count matrices. Cell, 187(23):6537–6549, 2024.

[14] Xianjun Dong, Xiaoqi Li, Tzuu-Wang Chang, Clemens R Scherzer, Scott T Weiss, and Weiliang Qiu. powereqtl: an r package and shiny application for sample size and power calculation of bulk tissue and single-cell eqtl analysis. Bioinformatics, 37(22):4269–4271, 2021.

[15] Katharina T Schmid, Barbara Höllbacher, Cristiana Cruceanu, Anika Böttcher, Heiko Lickert, Elisabeth B Binder, Fabian J Theis, and Matthias Heinig. scpower accelerates and optimizes the design of multi-sample single cell transcriptomic studies. Nature communications, 12(1):6625, 2021.

[16] Hyun Min Kang, Meena Subramaniam, Sasha Targ, Michelle Nguyen, Lenka Maliskova, Elizabeth McCarthy, Eunice Wan, Simon Wong, Lauren Byrnes, Cristina M Lanata, et al. Multiplexed droplet single-cell rna-sequencing using natural genetic variation. Nature biotechnology, 36(1):89–94, 2018.

[17] Drew Neavin, Anne Senabouth, Himanshi Arora, Jimmy Tsz Hang Lee, Aida Ripoll-Cladellas, sc-eQTLGen Consortium, Lude Franke, Shyam Prabhakar, Chun Jimmie Ye, Davis J McCarthy, et al. Demuxafy: improvement in droplet assignment by integrating multiple single-cell demultiplexing and doublet detection methods. Genome Biology, 25(1):94, 2024.

[18] George C Hartoularos, Yichen Si, Fan Zhang, Pooja Kathail, David S Lee, Anton Ogorodnikov, Yang Sun, Yun S Song, Hyun Min Kang, and Chun Jimmie Ye. Reference-free multiplexed single-cell sequencing identifies genetic modifiers of the human immune response. bioRxiv, pages 2023–05, 2023.

[19] Samuel L Wolock, Romain Lopez, and Allon M Klein. Scrublet: computational identification of cell doublets in single-cell transcriptomic data. Cell systems, 8(4):281–291, 2019.

[20] Christopher S McGinnis, Lyndsay M Murrow, and Zev J Gartner. Doubletfinder: doublet detection in single-cell rna sequencing data using artificial nearest neighbors. Cell systems, 8(4):329–337, 2019.

[21] Aaron TL Lun, Davis J McCarthy, and John C Marioni. A step-by-step workflow for low-level analysis of single-cell rna-seq data with bioconductor. F1000Research, 5:2122, 2016.

[22] W Evan Johnson, Cheng Li, and Ariel Rabinovic. Adjusting batch effects in microarray expression data using empirical bayes methods. Biostatistics, 8(1):118–127, 2007.

[23] Aaron T L. Lun, Karsten Bach, and John C Marioni. Pooling across cells to normalize single-cell rna sequencing data with many zero counts. Genome biology, 17:1–14, 2016.

[24] Christoph Hafemeister and Rahul Satija. Normalization and variance stabilization of single-cell rna-seq data using regularized negative binomial regression. Genome biology, 20(1):296, 2019.

[25] Andrew Butler, Paul Hoffman, Peter Smibert, Efthymia Papalexi, and Rahul Satija. Integrating single-cell transcriptomic data across different conditions, technologies, and species. Nature biotechnology, 36(5):411–420, 2018.

[26] Mark D Robinson and Alicia Oshlack. A scaling normalization method for differential expression analysis of rna-seq data. Genome biology, 11(3):1–9, 2010.

[27] Mark D Robinson and Alicia Oshlack. A scaling normalization method for differential expression analysis of rna-seq data. Genome biology, 11(3):1–9, 2010.

[28] T Mark Beasley, Stephen Erickson, and David B Allison. Rank-based inverse normal transformations are increasingly used, but are they merited? Behavior genetics, 39:580–595, 2009.

[29] Halit Ongen, Alfonso Buil, Andrew Anand Brown, Emmanouil T Dermitzakis, and Olivier Delaneau. Fast and efficient qtl mapper for thousands of molecular phenotypes. Bioinformatics, 32(10):1479–1485, 2016.

[30] Andrey A Shabalin. Matrix eqtl: ultra fast eqtl analysis via large matrix operations. Bioinformatics, 28(10): 1353–1358, 2012.

[31] Francesco Paolo Casale, Barbara Rakitsch, Christoph Lippert, and Oliver Stegle. Efficient set tests for the genetic analysis of correlated traits. Nature methods, 12(8):755–758, 2015.

[32] Zixuan Eleanor Zhang, Artem Kim, Noah Suboc, Nicholas Mancuso, and Steven Gazal. Efficient count-based models improve power and robustness for large-scale single-cell eqtl mapping. medRxiv, pages 2025–01, 2025.

[33] Yoav Benjamini and Yosef Hochberg. Controlling the false discovery rate: a practical and powerful approach to multiple testing. Journal of the Royal statistical society: series B (Methodological), 57(1):289–300, 1995.

[34] John D Storey and Robert Tibshirani. Statistical significance for genomewide studies. Proceedings of the National Academy of Sciences, 100(16):9440–9445, 2003.

[35] James Liley and Chris Wallace. Accurate error control in high-dimensional association testing using conditional false discovery rates. Biometrical Journal, 63(5):1096–1130, 2021.

[36] Igor Mandric, Tommer Schwarz, Arunabha Majumdar, Kangcheng Hou, Leah Briscoe, Richard Perez, Meena Subramaniam, Christoph Hafemeister, Rahul Satija, Chun Jimmie Ye, et al. Optimized design of single-cell rna sequencing experiments for cell-type-specific eqtl analysis. Nature communications, 11(1):5504, 2020.

[37] Anna S. E. Cuomo, Giordano Alvari, Christina B. Azodi, Davis J. McCarthy, Marc Jan Bonder, and single-cell eQTLGen consortium. Optimizing expression quantitative trait locus mapping workflows for single-cell studies. Genome Biology, 22(1):188, 2021.

[38] Wei Zhou, Anna Cuomo, Angli Xue, Masahiro Kanai, Grant Chau, Chirag Krishna, Ramnik J Xavier, Daniel G MacArthur, Joseph E Powell, Mark J Daly, et al. Efficient and accurate mixed model association tool for single-cell eqtl analysis. medRxiv, pages 2024–05, 2024.

[39] Yue Hu, Xi Xi, Qian Yang, and Xuegong Zhang. Sceqtl: an r package for identifying eqtl from single-cell parallel sequencing data. BMC bioinformatics, 21:1–12, 2020.

[40] Ariel DH Gewirtz, F William Townes, and Barbara E Engelhardt. Expression qtls in single-cell sequencing data. bioRxiv, pages 2022–08, 2022.

[41] David Lähnemann, Johannes Köster, Ewa Szczurek, Davis J McCarthy, Stephanie C Hicks, Mark D Robinson, Catalina A Vallejos, Kieran R Campbell, Niko Beerenwinkel, Ahmed Mahfouz, et al. Eleven grand challenges in single-cell data science. Genome biology, 21:1–35, 2020.

[42] Davide Risso, Fanny Perraudeau, Svetlana Gribkova, Sandrine Dudoit, and Jean-Philippe Vert. A general and flexible method for signal extraction from single-cell rna-seq data. Nature communications, 9(1):284, 2018.

[43] Giacomo Baruzzo, Ilaria Patuzzi, and Barbara Di Camillo. Sparsim single cell: a count data simulator for scrna-seq data. Bioinformatics, 36(5):1468–1475, 2020.

[44] Luke Zappia, Belinda Phipson, and Alicia Oshlack. Splatter: simulation of single-cell rna sequencing data. Genome biology, 18(1):174, 2017.

[45] Matteo Marouf, Pierre Machart, Vikas Bansal, Christoph Kilian, Daniel S Magruder, Christian F Krebs, and Stefan Bonn. Realistic in silico generation and augmentation of single-cell rna-seq data using generative adversarial networks. Nature communications, 11(1):166, 2020.

[46] Helena L Crowell, Charlotte Soneson, Pierre-Luc Germain, Daniela Calini, Ludovic Collin, Catarina Raposo, Dheeraj Malhotra, and Mark D Robinson. Muscat detects subpopulation-specific state transitions from multi-sample multi-condition single-cell transcriptomics data. Nature communications, 11(1):6077, 2020.

[47] Elizabeth A Wynn, Kara J Mould, Brian E Vestal, and Camille M Moore. Simulating paired and longitudinal single-cell rna sequencing data with rescuesim. Bioinformatics, 41(8):btaf442, 2025.

[48] Christina B Azodi, Luke Zappia, Alicia Oshlack, and Davis J McCarthy. splatpop: simulating population scale single-cell rna sequencing data. Genome biology, 22(1):341, 2021.

[49] Luca Bonomi, Yingxiang Huang, and Lucila Ohno-Machado. Privacy challenges and research opportunities for genomic data sharing. Nature genetics, 52(7):646–654, 2020.

[50] Nils Homer, Szabolcs Szelinger, Margot Redman, David Duggan, Waibhav Tembe, Jill Muehling, John V Pearson, Dietrich A Stephan, Stanley F Nelson, and David W Craig. Resolving individuals contributing trace amounts of dna to highly complex mixtures using high-density snp genotyping microarrays. PLoS genetics, 4(8):e1000167, 2008.

[51] Mara Thomas, Nuria Mackes, Asad Preuss-Dodhy, Thomas Wieland, Markus Bundschus, et al. Assessing privacy vulnerabilities in genetic data sets: Scoping review. JMIR Bioinformatics and Biotechnology, 5(1):e54332, 2024.

[52] Shuvom Sadhuka, Daniel Fridman, Bonnie Berger, and Hyunghoon Cho. Assessing transcriptomic reidentification risks using discriminative sequence models. Genome research, 33(7):1101–1112, 2023.

[53] Mayo Blegen Ashley L. 18 Wirkus Samantha J. 18 Wagner Victoria A. 18 Meyer Jeffrey G. 18 Cicek Mine S. 10 18 Biobank and All of Us Research Demonstration Project Teams Choi Seung Hoan 14 http://orcid.org/0000-0002-0322-8970 Wang Xin 14 http://orcid.org/00000001-6042-4487 Rosenthal Elisabeth A. 15. Genomic data in the all of us research program. Nature, 627(8003):340–346, 2024.

[54] Rafael Kramann, Christoph Kuppe, Valerie Luyckx, Wim Van Biesen, and Stefanie Steiger. Unveiling the risks: protecting privacy in single-cell genomics data. Nephrology Dialysis Transplantation, 40(6):1077–1080, 2025.

[55] Cynthia Dwork. Differential privacy. In International colloquium on automata, languages, and programming, pages 1–12. Springer, 2006.

[56] Bristena Oprisanu, Georgi Ganev, and Emiliano De Cristofaro. On utility and privacy in synthetic genomic data. arXiv preprint arXiv:2102.03314, 2021.

[57] Gidan Min and Junhyoung Oh. Can synthetic data protect privacy? IEEE Access, 2025.

[58] Dongyuan Song, Qingyang Wang, Guanao Yan, Tianyang Liu, Tianyi Sun, and Jingyi Jessica Li. scdesign3 generates realistic in silico data for multimodal single-cell and spatial omics. Nature biotechnology, pages 1–6, 2023.

[59] Wenan Chen, Yan Li, John Easton, David Finkelstein, Gang Wu, and Xiang Chen. Umi-count modeling and differential expression analysis for single-cell rna sequencing. Genome biology, 19(1):70, 2018.

[60] Jan Lause, Philipp Berens, and Dmitry Kobak. Analytic pearson residuals for normalization of single-cell rna-seq umi data. Genome biology, 22(1):258, 2021.

[61] Ruofan Ding, Qixuan Wang, Lihai Gong, Ting Zhang, Xudong Zou, Kewei Xiong, Qi Liao, Mireya Plass, and Lei Li. scqtlbase: an integrated human single-cell eqtl database. Nucleic Acids Research, 52(D1):D1010–D1017, 2024.

[62] Etienne Becht, Leland McInnes, John Healy, Charles-Antoine Dutertre, Immanuel WH Kwok, Lai Guan Ng, Florent Ginhoux, and Evan W Newell. Dimensionality reduction for visualizing single-cell data using umap. Nature biotechnology, 37(1):38–44, 2019.

[63] Nona Farbehi, Drew R Neavin, Anna SE Cuomo, Lorenz Studer, Daniel G MacArthur, and Joseph E Powell. Integrating population genetics, stem cell biology and cellular genomics to study complex human diseases. Nature Genetics, 56(5):758–766, 2024.

[64] BJ Strober, Reem Elorbany, Katherine Rhodes, Nirmal Krishnan, Karl Tayeb, Alexis Battle, and Yoav Gilad. Dynamic genetic regulation of gene expression during cellular differentiation. Science, 364(6447):1287–1290, 2019.

[65] Kevin R Moon, David Van Dijk, Zheng Wang, Scott Gigante, Daniel B Burkhardt, William S Chen, Kristina Yim, Antonia van den Elzen, Matthew J Hirn, Ronald R Coifman, et al. Visualizing structure and transitions in high-dimensional biological data. Nature biotechnology, 37(12):1482–1492, 2019.

[66] Zhan Su, Jonathan Marchini, and Peter Donnelly. Hapgen2: simulation of multiple disease snps. Bioinformatics, 27(16):2304–2305, 2011.

[67] Sebastian Ueckert, Mats O Karlsson, and Andrew C Hooker. Accelerating monte carlo power studies through parametric power estimation. Journal of pharmacokinetics and pharmacodynamics, 43:223–234, 2016.

[68] Peter Green and Catriona J MacLeod. Simr: An r package for power analysis of generalized linear mixed models by simulation. Methods in Ecology and Evolution, 7(4):493–498, 2016.

[69] Levi Kumle, Melissa L-H Võ, and Dejan Draschkow. Estimating power in (generalized) linear mixed models: An open introduction and tutorial in r. Behavior research methods, 53(6):2528–2543, 2021.

[70] Martin Jinye Zhang, Vasilis Ntranos, and David Tse. Determining sequencing depth in a single-cell rna-seq experiment. Nature communications, 11(1):774, 2020.

[71] Tom Meyvis and Stijn MJ Van Osselaer. Increasing the power of your study by increasing the effect size. Journal of Consumer Research, 44(5):1157–1173, 2018.

[72] Pak C Sham and Shaun M Purcell. Statistical power and significance testing in large-scale genetic studies. Nature Reviews Genetics, 15(5):335–346, 2014.

[73] Ceyhan Ceran Serdar, Murat Cihan, Doğan Yücel, and Muhittin A Serdar. Sample size, power and effect size revisited: simplified and practical approaches in pre-clinical, clinical and laboratory studies. Biochemia medica, 31(1):27–53, 2021.

[74] Ruochen Jiang, Tianyi Sun, Dongyuan Song, and Jingyi Jessica Li. Statistics or biology: the zero-inflation controversy about scrna-seq data. Genome biology, 23(1):31, 2022.

[75] Anna SE Cuomo, Eleanor Spenceley, Hope A Tanudisastro, Blake Bowen, Albert Henry, Hao Lawrence Huang, Angli Xue, Wei Zhou, Matthew J Welland, Arthur S Lee, et al. Impact of rare and common genetic variation on cell type-specific gene expression. medRxiv, pages 2025–03, 2025.

[76] Sung Eun Hong, Seon Ju Mun, Young Joo Lee, Taekyeong Yoo, Kyung-Suk Suh, Keon Wook Kang, Myung Jin Son, Won Kim, and Murim Choi. Single-cell eqtl analysis identifies genetic variation underlying metabolic dysfunction-associated steatohepatitis. Nature Genetics, pages 1–11, 2025.

[77] Kian Hong Kock, Kyung Yeon Han, Yoshinari Ando, Damita Jevapatarakul, Ankita Chatterjee, Quy Xiao Xuan Lin, Eliora Violain Buyamin, Radhika Sonthalia, Deepa Rajagopalan, Yoshihiko Tomofuji, et al. Asian diversity in human immune cells. Cell, 188(8):2288–2306, 2025.

[78] Zepeng Mu, Haley E. Randolph, Raúl Aguirre-Gamboa, Ellen Ketter, Anne Dumaine, Veronica Locher, Cary Brandolino, Xuanyao Liu, Daniel E. Kaufmann, Luis B. Barreiro, and Yang I. Li. Impact of disease-associated chromatin accessibility qtls across immune cell types and contexts. Cell Genomics, 6(1), 2026. ISSN 2666-979X. doi:10.1016/j.xgen.2025.101061. URL https://doi.org/10.1016/j.xgen.2025.101061.

[79] Adam W Turner, Shengen Shawn Hu, Jose Verdezoto Mosquera, Wei Feng Ma, Chani J Hodonsky, Doris Wong, Gaëlle Auguste, Yipei Song, Katia Sol-Church, Emily Farber, et al. Single-nucleus chromatin accessibility profiling highlights regulatory mechanisms of coronary artery disease risk. Nature Genetics, 54(6):804–816, 2022.

[80] Paola Benaglio, Jacklyn Newsome, Jee Yun Han, Joshua Chiou, Anthony Aylward, Sierra Corban, Michael Miller, Mei-Lin Okino, Jaspreet Kaur, Sebastian Preissl, et al. Mapping genetic effects on cell type-specific chromatin accessibility and annotating complex immune trait variants using single nucleus atac-seq in peripheral blood. PLoS genetics, 19(6):e1010759, 2023.

[81] Jun Wang, Xuesen Cheng, Qingnan Liang, Leah A Owen, Jiaxiong Lu, Yiqiao Zheng, Meng Wang, Shiming Chen, Margaret M DeAngelis, Yumei Li, et al. Single-cell multiomics of the human retina reveals hierarchical transcription factor collaboration in mediating cell type-specific effects of genetic variants on gene regulation. Genome Biology, 24(1):269, 2023.

[82] Rushdy Ahmad and Bogdan Budnik. A review of the current state of single-cell proteomics and future perspective. Analytical and Bioanalytical Chemistry, 415(28):6889–6899, 2023.

[83] Apostolos Dimitromanolakis, Jingxiong Xu, Agnieszka Krol, and Laurent Briollais. sim1000g: a user-friendly genetic variant simulator in r for unrelated individuals and family-based designs. BMC bioinformatics, 20:1–9, 2019.

[84] Chris CA Spencer, Zhan Su, Peter Donnelly, and Jonathan Marchini. Designing genome-wide association studies: sample size, power, imputation, and the choice of genotyping chip. PLoS genetics, 5(5):e1000477, 2009.

[85] Sophie Wharrie, Zhiyu Yang, Vishnu Raj, Remo Monti, Rahul Gupta, Ying Wang, Alicia Martin, Luke J O’Connor, Samuel Kaski, Pekka Marttinen, et al. Hapnest: efficient, large-scale generation and evaluation of synthetic datasets for genotypes and phenotypes. Bioinformatics, 39(9):btad535, 2023.

[86] Tianyi Sun, Dongyuan Song, Wei Vivian Li, and Jingyi Jessica Li. scdesign2: a transparent simulator that generates high-fidelity single-cell gene expression count data with gene correlations captured. Genome biology, 22(1):163, 2021.

[87] Shaun Purcell, Benjamin Neale, Kathe Todd-Brown, Lori Thomas, Manuel AR Ferreira, David Bender, Julian Maller, Pamela Sklar, Paul IW De Bakker, Mark J Daly, et al. Plink: a tool set for whole-genome association and population-based linkage analyses. The American journal of human genetics, 81(3):559–575, 2007.

[88] Petr Danecek, James K Bonfield, Jennifer Liddle, John Marshall, Valeriu Ohan, Martin O Pollard, Andrew Whitwham, Thomas Keane, Shane A McCarthy, Robert M Davies, et al. Twelve years of samtools and bcftools. Gigascience, 10(2):giab008, 2021.

[89] Adam Frankish, Mark Diekhans, Anne-Maud Ferreira, Rory Johnson, Irwin Jungreis, Jane Loveland, Jonathan M Mudge, Cristina Sisu, James Wright, Joel Armstrong, et al. Gencode reference annotation for the human and mouse genomes. Nucleic acids research, 47(D1):D766–D773, 2019.

[90] Zuguang Gu, Roland Eils, and Matthias Schlesner. Complex heatmaps reveal patterns and correlations in multidimensional genomic data. Bioinformatics, 32(18):2847–2849, 2016.

[91] Ilya Korsunsky, Nghia Millard, Jean Fan, Kamil Slowikowski, Fan Zhang, Kevin Wei, Yuriy Baglaenko, Michael Brenner, Po-ru Loh, and Soumya Raychaudhuri. Fast, sensitive and accurate integration of single-cell data with harmony. Nature methods, 16(12):1289–1296, 2019.

[92] Paul Robert and Yves Escoufier. A unifying tool for linear multivariate statistical methods: the rv-coefficient. Journal of the Royal Statistical Society Series C: Applied Statistics, 25(3):257–265, 1976.

[93] John Lonsdale, Jeffrey Thomas, Mike Salvatore, Rebecca Phillips, Edmund Lo, Saboor Shad, Richard Hasz, Gary Walters, Fernando Garcia, Nancy Young, et al. The genotype-tissue expression (gtex) project. Nature genetics, 45 (6):580–585, 2013.

[94] Davide Chicco and Giuseppe Jurman. The advantages of the matthews correlation coefficient (mcc) over f1 score and accuracy in binary classification evaluation. BMC genomics, 21(1):6, 2020.

