## Supplementary Materials for "scDesignPop generates realistic population-scale single-cell RNA-seq for power analysis, benchmarking, and privacy protection"

### Supplementary Methods

#### S1 Modeling real population-scale scRNA-seq datasets

The marginal model in Equation 1 can be simplified depending on the users' choice of design covariates, assumptions about the interaction effects, and the size of the scRNA-seq dataset. Below, we specify the gene marginal models used to verify scDesignPop as a realistic simulator with OneK1K and CLUES cohorts.

**Modeling OneK1K dataset**

We modeled UMI counts using a negative binomial (NB) mixed distribution, denoted $F^{\mathrm{NB}}$, for the gene marginal model (Equation 1). Here, the location and scale parameters (i.e., $\mu_{ij}$ and $\phi_j$) are the mean and dispersion parameters of the NB distribution, respectively. For the NB distribution, the link function for the mean is $\log(\mu_{ij})$. We evaluated both single-SNP and multi-SNP modes of scDesignPop on the same dataset, using labeled cell types from the OneK1K cohort. For model fitting, we randomly selected 100 individuals and performed stratified sampling to select 70% of cells from each cell type per individual, yielding a training set of 92,304 cells. The remaining 30% of cells served as held-out test data.

For the single-SNP mode, we chose one putative cell-type-specific eQTL (cts-eQTL) per gene (i.e., a SNP with genotype $G_{k,j}$), and did not include any additional design covariates. Interaction effects between cell type and genotype were also omitted. Under this specification, the conditional mean of the gene marginal model is

$$\log(\mu_{ij}) = \alpha_j + u_{kj} + G_{kj}\boldsymbol{\beta}_j + s_i\gamma_{jh} \ . \tag{19}$$

For the multi-SNP mode, we permitted one or more putative cts-eQTLs per gene (i.e., a set of SNPs with genotype vector $\mathbf{G}_{k,l(j)}$). We included the individual random effect, age and gender as design covariates $\mathbf{c}_i$, as well as an interaction effect between cell type and genotype denoted $\mathbf{G}_{k,l(j)} \otimes s_i$. The conditional mean of the gene marginal model is

$$\log(\mu_{ij}) = \alpha_j + u_{kj} + \mathbf{G}_{k,l(j)}^\top\boldsymbol{\beta}_j + s_i\gamma_{jh} + \mathbf{c}_i \cdot \delta_j + (\mathbf{G}_{k,l(j)} \otimes s_i)^\top\boldsymbol{\zeta}_j \ . \tag{20}$$

Since all individuals in the OneK1K cohort were of Northern European genetic ancestry [5], we opted not to include genotype principal components (PCs) as covariates in both single-SNP and multi-SNP modes.

**Modeling CLUES dataset**

We modeled the normalized expression using a linear mixed (LM) distribution, denoted $F^{\mathrm{LM}}$, for the gene marginal model in Equation 1. Here, the location and scale parameters correspond to the mean and variance parameters of the Gaussian distribution, respectively. For the Gaussian distribution, the link function for the mean is the identity $\mu_{ij}$. We evaluated both single-SNP and multi-SNP modes of scDesignPop on the same dataset, using the labeled cell types from the CLUES study. For model fitting, we randomly selected 150 individuals and performed stratified sampling to select

70% of cells from each cell type per individual, yielding a training set of 473,208 cells. The remaining 30% of cells served as held-out test data.

For the single-SNP mode, we have one putative cts-eQTL for each gene. We included an individual random effect $u_{kj}$, and ancestry and disease status as design covariates $\mathbf{c}_i$. We also included interaction effects between cell-types and genotype $G_{k,l(j)} \otimes s_i$, and between each design covariate and genotype $G_{k,l(j)} \otimes \mathbf{c}_i$. The conditional mean of the gene marginal model is

$$\mu_{ij} = \alpha_j + u_{kj} + G_{k,l(j)}\beta_j + s_i\gamma_{jh} + \mathbf{c}_i^\top \boldsymbol{\delta}_j + (G_{k,l(j)} \otimes s_i)^\top \boldsymbol{\zeta}_j + (G_{k,l(j)} \otimes \mathbf{c}_i)^\top \boldsymbol{\eta}_j \ . \tag{21}$$

For the multi-SNP mode, we permitted one or more putative cts-eQTLs per gene. We included an individual random effect $u_{kj}$, ancestry and disease status as design covariates $\mathbf{c}_i$, and an interaction effect between cell type and genotypes $G_{k,l(j)} \otimes \mathbf{c}_i$. The conditional mean of the gene marginal model is

$$\mu_{ij} = \alpha_j + u_{kj} + \mathbf{G}_{k,l(j)}^\top \boldsymbol{\beta}_j + s_i\gamma_{jh} + \mathbf{c}_i^\top \boldsymbol{\delta}_j + (\mathbf{G}_{k,l(j)} \otimes s_i)^\top \boldsymbol{\zeta}_j \ . \tag{22}$$

Motivated by Perez et al.'s findings that ancestry and disease status influence genetic associations in systemic lupus erythematosus (SLE) [6], we included both variables as covariates in single-SNP and multi-SNP modes. Since ancestry was already included as a categorical variable in both our single-SNP and multi-SNP modes, we opted not to include genotype PCs.

**Modeling cell-type proportions in OneK1K and CLUES**

To generate scRNA-seq data for new individuals, scDesignPop additionally models the distribution of cell-type proportions observed in the training data. Specifically, scDesignPop first models the total number of cells per individual using a log-normal distribution. Then, conditional on this, a multinominal logit regression model with individual-level covariates are used as predictors (Methods 10.1.4). This is implemented in `simuCellProportion` function in scDesignPop.

To simulate cell-type proportions in OneK1K data, we used the cell type labels as the response variable and included age, sex, and the top 30 genotype PCs as covariates from a subset of 70 individuals. Next, we used scDesignPop to model the cell-type proportions and simulated the cell-type compositions for 100 individuals (including the original 70 individuals) using the fitted model. Similarly, for CLUES data, we used the cell type labels as the response variable and included ancestry, disease status, and the top 30 genotype PCs from a subset of 105 individuals as covariates. Next, we used scDesignPop to model the cell-type proportions and simulated the cell-type compositions for 150 individuals (including the original 105 individuals) using the fitted model.

Genotype PCs were computed using `PLINK` (v1.90 beta 7.2) using observed genotypes from all individuals in each cohort (981 individuals in OneK1K; 256 individuals in CLUES). After concatenating the VCF files from autosomes (chromosomes $1-22$) and converting them into PLINK format, we then filtered variants with a minor allele frequency

(MAF) threshold of 0.05. Next, we pruned SNPs using PLINK's `-indep-pairwise` mode with a window size of 50, a step size of 10, and a correlation threshold of 0.2. Finally, PCA was then performed on the remaining variants to obtain the top 30 genotype PCs.

#### S2 Parameter estimation stability

We assessed the stability of parameter estimates for cell type effect and cts-eQTL effect under different data regimes by repeatedly subsampling the number of individuals using the OneK1K cohort and fitting scDesignPop's marginal model for every gene. To do this, we used all cells from subsampled individuals from $\{20, 30, 50, 100, 150, 200\}$ and fitted a NB mixed model with cell type, putative eQTLs under single-SNP mode, and a cell-type-genotype interaction effect using scDesignPop for 810 highly variable genes. Then, the fitted parameter estimates for cell type effect and cts-eQTL effect, defined as the per-allele-eQTL effect, were compiled. This process was repeated 20 times for each sample size setting. The mean and standard error across replicates were computed for each gene and cell type group.

#### S3 Power analysis workflow

Using scDesignPop's estimated eQTL effects as ground truth, simulation-based power analysis can be conducted to estimate the statistical power, which is the probability of detecting an eQTL effect. A detailed workflow of power analysis using scDesignPop is shown below (Fig. S25):

1. For the selected gene $j$, we fit the full marginal model with eQTL effect to the training data as described in Methods S1 with Equation 20 for OneK1K and Equation 22 for CLUES. Based on the full model, we record the genotype effect size from the targeted SNP in each cell type $\boldsymbol{\beta}_{jh}$ and then set the genotype effect and all its interaction effects to zero to obtain the null model.

2. For a selected cell type $h$, we resample the genotype of the corresponding SNP and other covariates from $\tilde{K}$ individuals (indexed by $\tilde{k}$) and $n_{\tilde{k}}$ cells per individual for all $\tilde{k} \in \{1, \ldots, \tilde{K}\}$ in the original data in $B$ simulation times ($b \in \{1, \ldots, B\}$). In total, $n_{\tilde{k}}$ cells will be sampled for each individual $\tilde{k}$ in each parameter setting. To maintain the allele frequency, the numbers of individuals in genotype $0, 1, 2$ are calculated based on the proportions of individuals from real data.

3. Based on the full model, we predict new count values $\tilde{\mathbf{y}}_{j,H_1}^{(b)}$ as alternative ($H_1$) data with nonzero genotype effect size in the cell type. Similarly, we predict new count values $\tilde{\mathbf{y}}_{j,H_0}^{(b)}$ as null ($H_0$) data based on the null model with zero genotype effect size.

4. Then we fit the alternative data with the user-specified eQTL model to get $\hat{\beta}_{H_1}^{(b)}$ as an alternative genotype effect size estimate. With the same eQTL model, we fit the null data to get $\hat{\beta}_{H_0}^{(b)}$ as a null genotype effect size estimate.

5. Collect $B$ different $\hat{\beta}_{H_1}^{(b)}$ and $\hat{\beta}_{H_0}^{(b)}$ to obtain a vector $\hat{\boldsymbol{\beta}}_{H_1}$ and a vector $\hat{\boldsymbol{\beta}}_{H_0}$, respectively.

6. Then, we resample both the $\hat{\boldsymbol{\beta}}_{H_1}$ and $\hat{\boldsymbol{\beta}}_{H_0}$ for $x = 1000$ times to generate 1000 groups of the vector $\hat{\boldsymbol{\beta}}_{H_1}$ and vector $\hat{\boldsymbol{\beta}}_{H_0}$. In each group, if $\boldsymbol{\beta}_{jh} > 0$, we then calculate the critical value as the $1 - \alpha_{\text{adj}}$ percentile of $\hat{\boldsymbol{\beta}}_{H_0}$, the proportion of $\hat{\boldsymbol{\beta}}_{H_1}$ that are above the critical value is obtained as the statistical power. If $\boldsymbol{\beta}_{jh} < 0$, we then calculate the critical value as the $\alpha_{\text{adj}}$ percentile of $\hat{\boldsymbol{\beta}}_{H_0}$, the proportion of $\hat{\boldsymbol{\beta}}_{H_1}$ that are below the critical value is obtained as the statistical power. The setting of the $\alpha_{\text{adj}}$ value is introduced in **Multiple testing correction** (Methods 10.8).

7. Lastly, given a particular cell type $h$ with $\tilde{K}$ individuals and $n_{\tilde{k}}$ cells per individual, the average statistical power with the standard deviation is calculated based on the statistical power obtained in each group.

#### S4 eQTL models for power analysis

scDesignPop also allows users to specify multiple different eQTL models to assess the power of eQTL effect detection in each cell type, we currently include four options: 1) negative binomial mixed (NB mixed) model, 2) Poisson mixed model, 3) linear mixed model, and 4) pseudobulk linear model. Among them, NB mixed, Poisson mixed, and linear mixed are fitted directly at the cell-type level. Linear mixed models are capable of fitting normalized, scaled scRNA-seq data. For pseudobulk linear, we first sum the single-cell expressions across cells for each individual, then we fit a linear model without a random effect. The eQTL model is partially inherited from the full marginal model Equation 1. If the marginal full model is the same as in Methods 10.1, in each cell type, we fit a selected model to obtain $\boldsymbol{\beta}'_j$ estimates ($\hat{\boldsymbol{\beta}}_{H_0}$ and $\hat{\boldsymbol{\beta}}_{H_1}$, respectively) for both null and alternative data.

**Negative binomial, Poisson, or linear mixed model**

Here, single-cell level expression $Y_{ij}$ follows the model

$$\begin{cases} Y_{ij}|\mathbf{G}', \mathbf{z}'_i, \mathbf{c}'_i \overset{\text{ind.}}{\sim} F_j(\,\cdot\,|\mathbf{G}', \mathbf{z}'_i, \mathbf{c}'_i; \mu'_{ij}, \phi'_j) \\ \text{if } z'_{ik} = 1 \ \ (\text{cell } i \text{ belongs to individual } k): \\ \log(\mu'_{ij}) = \alpha'_j + u'_{kj} + \mathbf{G}'^{\top}_{k,l(j)}\boldsymbol{\beta}'_j + \mathbf{c}'^{\top}_i\boldsymbol{\delta}'_j + (\mathbf{G}'_{k,l(j)} \otimes \mathbf{c}'_i)^{\top}\boldsymbol{\eta}'_j \\ u'_{kj} \overset{\text{ind.}}{\sim} \mathcal{N}(0, \sigma^{2'}_j) \end{cases}, \tag{23}$$

where $\phi'_j, \alpha'_j, \boldsymbol{\beta}'_j, \boldsymbol{\delta}'_j, \boldsymbol{\eta}'_j, \sigma^{2'}_j$ are new eQTL model parameters estimated based on the simulated data with covariates $\mathbf{G}', \mathbf{z}'_i, \mathbf{c}'_i$ from the marginal full model for each gene $j$. Here, $F_j$ can be negative binomial, Poisson, or linear mixed model. When it's for negative binomial or linear mixed models, $\mu'_{ij}$ and $\phi'_j$ are the corresponding mean and dispersion parameters. When it's for Poisson mixed model, there is no $\phi'_j$ exists and $\mu'_{ij}$ is the mean parameter.

**Pseudobulk linear model**

Define $\mathbf{C}' = [\mathbf{a}' \ \mathbf{b}' \ \mathbf{e}'] = [\mathbf{c}'^{\top}_1 \cdots \mathbf{c}'^{\top}_d] \in \mathbb{R}^{d \times q}$: an (optional) individual-by-covariate design matrix with $d$ individuals, and $q$ total dimensions corresponding to individual-level covariates $\mathbf{a}' = [\mathbf{a}'_1, \ldots, \mathbf{a}'_d]^{\top}$ (such as age, gender, etc), batch covariate $\mathbf{b}' = (b'_1, \ldots, b'_d)^\top$ has $b'_i \in \{1, \ldots, B\}$, and experiment condition covariate where $\mathbf{e}' = (e'_1, \ldots, e'_d)^\top$ has $e'_k \in \{1, \ldots, E\}$, respectively. We refer to $\mathbf{c}'_k$ as the *design covariates* for individual $k$ here. Define $\mu'_{kj}$ and $\phi'_j$ as the mean and variance parameters of Gaussian distributions for pseudobulk expression values. We sum $Y_{ij}$ across cells from each individual $k$ in each cell type to get pseudobulk expression $Y_{kj}$. Here, $Y_{kj}$ follows a linear model

$$\begin{cases} Y_{kj} | \mathbf{G}', \mathbf{c}'_k \overset{\text{ind.}}{\sim} F_j(\,\cdot\, | \mathbf{G}', \mathbf{c}'_k; \mu'_{kj}, \phi'_j) \\ \mu'_{kj} = \alpha'_j + \mathbf{G}'^\top_{k,l(j)} \boldsymbol{\beta}'_j + \mathbf{c}'^\top_k \boldsymbol{\delta}'_j + (\mathbf{G}'_{k,l(j)} \otimes \mathbf{c}'_k)^\top \boldsymbol{\eta}'_j \end{cases}, \tag{24}$$

where $\phi'_j, \alpha'_j, \boldsymbol{\beta}'_j, \boldsymbol{\delta}'_j, \boldsymbol{\eta}'_j$ are new eQTL model parameters estimated based on the simulated data with covariates $\mathbf{G}', \mathbf{c}'_k$ from the marginal full model for each gene $j$. $\mu'_{ij}$ and $\phi'_j$ are the corresponding mean and dispersion parameters

### Supplementary Tables

**Table S1: Comparison of main functionalities between scDesignPop, scDesign3, and multi-individual scRNA-seq simulators.**

| Functionality | scDesignPop | scDesign3 | splatPop | muscat | rescueSim |
|---|---|---|---|---|---|
| New individual simulation | ✓ | ✗ | ✓ | ✗ | ✗ |
| eQTL effects captured | ✓ | ✗ | ✓ | ✗ | ✗ |
| Multiple eSNPs per gene | ✓ | ✗ | ✗ | ✗ | ✗ |
| Dynamic eQTL modeling | ✓ | ✗ | ✗ | ✗ | ✗ |
| Flexible distribution family | ✓ | ✓ | ✗ | ✗ | ✗ |
| Gene cor. preserved | ✓ | ✓ | ✗ | ✓ | ✗ |
| Covariate flexibility | ✓ | ✓ | ✗ | ✗ | ✗ |
| Power analysis | ✓ | ✗ | ✗ | ✗ | ✗ |

**Notes:**
New individual simulation: ability to simulate scRNA-seq data for previously unseen individuals.
eQTL effect modeling: ability to model eQTL effects.
Multiple eSNPs per gene: supports modeling of multiple putative eSNPs per gene.
Dynamic eQTL modeling: supports modeling of dynamic eQTL effects across cell states.
Flexible distribution family: supports multiple parametric families for modeling gene expression.
Gene cor. preserved: preserves pairwise gene-gene correlations.
Covariate flexibility: supports multiple (more than two) user-specified covariates.
Power analysis: includes built-in functionality for eQTL power analysis.

**Table S2: Comparison of main functionalities between scDesignPop, and alternative eQTL power analysis tools.**

| Functionality | scDesignPop | PowerEQTL | scPower |
|---|---|---|---|
| Data-driven power estimation | ✓ | ✗ | ✓ |
| Covariate flexibility | ✓ | ✗ | ✗ |
| eQTL-specific power estimates | ✓ | ✓ | ✗ |
| Cell-type-aware study design | ✓ | ✗ | ✓ |
| Flexible eQTL model choices | ✓ | ✓ | ✗ |
| Expected power trends | ✓ | ✗ | ✗ |

**Notes:**
Data-driven power estimation: estimates power from empirical scRNA-seq data.
Covariate flexibility: ability to incorporate additional covariates beyond the eQTL effect.
eQTL-specific power estimates: reports power for the selected eQTL.
Cell-type-aware study design: supports cell types as input for customizable power analysis designs.
Flexible eQTL model choices: supports multiple eQTL model specifications.
Expected power trends: produces expected power trends with increasing sample size (number of individuals and number of cells per individual) or varying effect size.

**Table S3: Parametric families supported in scDesignPop's gene marginal modeling**

| Distribution | Parameters | Probability density function (PDF) or Probability mass function (PMF) | Link function | Applicable data type |
|---|---|---|---|---|
| Gaussian | $\mu$: mean<br>$\sigma$: standard deviation | $f(x) = \frac{1}{\sigma\sqrt{2\pi}} e^{-\frac{1}{2}(\frac{x-\mu}{2})^2}$ | $g(\mu) = \mu$ | Normalized data |
| Bernoulli | $\mu$: mean | $f(x) = \mu^x (1-\mu)^{1-x}; x \in \{0, 1\}$ | $g(\mu) = \log \frac{\mu}{1-\mu}$ | Binary data |
| Poisson | $\mu$: mean | $f(x) = \frac{\mu^x e^{-\mu}}{x!}; x \in \{0, 1, 2, \cdots\}$ | $g(\mu) = \log \mu$ | Count data without over-dispersion |
| Negative Binomial | $\mu$: mean<br>$\phi$: dispersion | $f(x) = \frac{\Gamma(x+\frac{1}{\phi})}{\Gamma(\frac{1}{\phi})\Gamma(x+1)} (\frac{1}{1+\phi\mu})^{\frac{1}{\phi}} (\frac{\phi\mu}{1+\phi\mu})^x; x \in \{0, 1, 2, \cdots\}$ | $g(\mu) = \log \mu$ | Count data with over-dispersion |

**Notes:** The cell index $i$ and gene index $j$ are dropped here for notation simplicity.

**Table S4: Benchmark results for eQTL mapping methods under different simulation settings.** Each value represents a performance metric for the indicated cell type. **Bolded** value indicates the best performing method for a metric in a cell type and simulation setting. Area under the receiver operating characteristic curve (AUROC) and area under the precision-recall curve (AUPRC) correspond to the curves shown in Fig. 6B and C. We additionally report the Matthew's correlation coefficient (MCC), the relative improvement over baseline AUPRC (Rel. impr. AUPRC), total computational time in CPU hours, and peak memory usage in GiB. For CPU hours and peak memory, lower values indicate better performance, while for the remaining metrics, higher values indicate better performance.

| **Simulation setting** | **Method** | **Memory B cells** | | | | **$CD4_{NC}$ T cells** | | | | | |
|---|---|---|---|---|---|---|---|---|---|---|---|
| | | AUROC | AUPRC | Rel. impr. AUPRC | MCC | AUROC | AUPRC | Rel. impr. AUPRC | MCC | CPU hours | Peak mem. (GiB) |
| Fitted | FastQTL | **70.92** | **48.95** | **36.76** | **0.2982** | **70.33** | **74.48** | **47.39** | **0.2985** | 30.4 | **0.293** |
| | SAIGE-QTL | 70.42 | 44.98 | 31.86 | 0.2306 | 69.16 | 72.59 | 43.51 | 0.2664 | 618.9 | 7.32 |
| | jaxQTL_perm | 67.61 | 44.20 | 30.89 | 0.1870 | 69.63 | 73.52 | 45.43 | 0.2762 | 120.4 | 13.09 |
| | jaxQTL_ACAT | 51.66 | 21.23 | 2.44 | 0.000693 | 59.09 | 60.38 | 18.34 | 0.0864 | **1.49** | 7.23 |
| True H0 | FastQTL | **83.21** | 74.31 | 68.18 | 0.6542 | 82.78 | 87.81 | 74.87 | 0.5963 | – | – |
| | SAIGE-QTL | 82.03 | 76.27 | 70.61 | **0.7308** | **84.26** | **89.19** | **77.71** | **0.6192** | – | – |
| | jaxQTL_perm | 82.91 | **76.75** | **71.21** | 0.7005 | 83.48 | 88.93 | 77.17 | 0.6113 | – | – |
| | jaxQTL_ACAT | 73.15 | 58.93 | 49.14 | 0.4896 | 77.79 | 83.06 | 65.08 | 0.4327 | – | – |
| True H0 + higher eQTL | FastQTL | **88.17** | 81.66 | 77.29 | 0.7721 | 87.93 | 91.71 | 82.92 | 0.7079 | – | – |
| | SAIGE-QTL | 87.29 | **82.84** | **78.75** | **0.7990** | **89.39** | **92.38** | **84.30** | **0.7163** | – | – |
| | jaxQTL_perm | 87.79 | 82.77 | 78.66 | 0.7686 | 87.96 | 92.00 | 83.51 | 0.6901 | – | – |
| | jaxQTL_ACAT | 72.81 | 56.60 | 46.24 | 0.4355 | 72.99 | 78.35 | 55.38 | 0.3994 | – | – |

**Notes:** Rel. impr. AUPRC is computed using $\frac{\text{AUPRC}-\pi}{1-\pi}$, where $\pi$ is the baseline proportion of true cts-eGenes. MCC is computed at a 0.05 target FDR threshold. Dashes (–) indicate metric not measured. *Abbreviations:* central processing unit (CPU); gibibyte (GiB).

**Table S5: Modified parameter settings used to generate synthetic scRNA-seq data under user-specified simulation settings.**

| **Simulation setting** | **Gene set** | **Cell type** | **`neg_ctrl`** | **`mean_log2fc`** | **`eqtl_log2fc`** |
|---|---|---|---|---|---|
| True H0 | 654 non-eGenes | Memory B cells | `TRUE` | 0 | 0 |
| | 393 non-eGenes | $CD4_{NC}$ T cells | `TRUE` | 0 | 0 |
| True H0 + higher eQTL | 156 eGenes | Memory B cells | `FALSE` | 0 | 0.5 |
| | 654 non-eGenes | Memory B cells | `TRUE` | 0 | 0 |
| | 417 eGenes | $CD4_{NC}$ T cells | `FALSE` | 0 | 0.5 |
| | 393 non-eGenes | $CD4_{NC}$ T cells | `TRUE` | 0 | 0 |

**Notes:** For the "Fitted" simulation setting, no parameter modifications were made, and hence not listed in the table. The `neg_ctrl` option sets the per-allele eQTL effect size to 0 on the link scale for genes within a cell type. The `mean_log2fc` option modifies the conditional mean on link scale via log2-fold-changes for genes in a cell type. The `eqtl_log2fc` option modifies the per-allele eQTL effect size on link scale via log2-fold-changes for genes within a cell type. All other parameters used were set to their default in scDesignPop's `modifyMarginalModels` function. *Abbreviations*: memory B cells (bmem); $CD4_{NC}$ T cells (cd4nc).

**Table S6: Computational time for marginal gene model fitting in scDesignPop**. The average computation time in seconds per gene across different datasets and model specifications. Averages and standard deviations are computed using 817 genes for OneK1K cohort and 798 genes for CLUES cohort.

| Dataset | # of covariates | SNP mode | Distribution | # of cells | # of individuals | Time (sec) |
|---|---|---|---|---|---|---|
| OneK1K | 2 | single-SNP | Negative Binomial | 92,304 | 100 | 6.4 ± 5.9 |
| | 5 | multi-SNP | Negative Binomial | 92,304 | 100 | 11.5 ± 10.1 |
| CLUES | 7 | single-SNP | Gaussian | 473,208 | 150 | 17.0 ± 7.6 |
| | 5 | multi-SNP | Gaussian | 473,208 | 150 | 17.9 ± 8.8 |

**Table S7: Computational costs under simulation scenarios to generate millions of synthetic cells.** An scDesignPop pipeline consisting of data input construction, marginal model fitting, copula model fitting, and synthetic data generation was used to evaluate the total computation time in CPU hours and peak memory usage in gibibyte. With scDesignPop's parallelization, 20 CPU cores were utilized to train the marginal models, and 2 CPU cores were used to train the copula and for generating synthetic data. All simulations were trained on 70 individuals from the OneK1K cohort across 810 genes, with each simulation scenario performed once. *Abbreviations:* central processing unit (CPU); gibibyte (GiB).

| Training cells | Training individuals | Synthetic cells | Synthetic individuals | CPU hours | Peak memory (GiB) |
|---|---|---|---|---|---|
| 93,376 | 70 | 3,803,304 | 2,943 | 13.2 | 419 |
| | | 5,071,072 | 3,924 | 15.6 | 461 |
| | | 6,338,840 | 4,905 | 18.9 | 597 |

#### Supplementary Figures

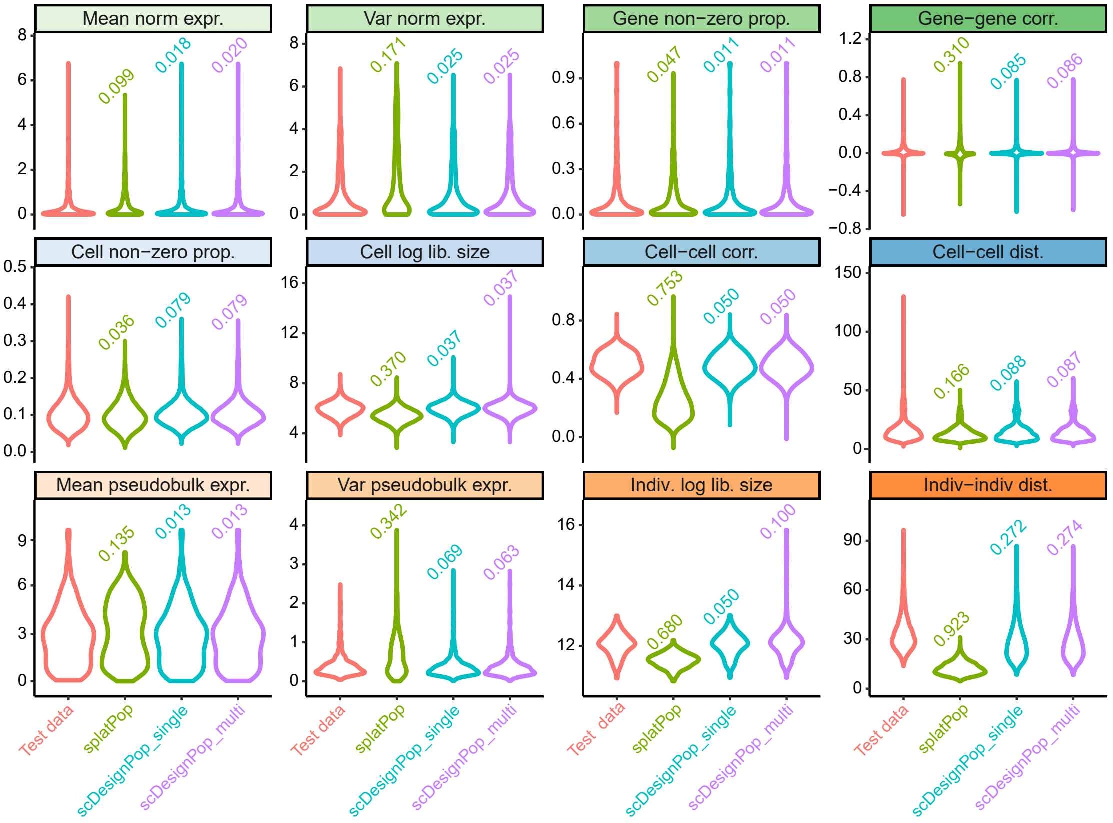


**Fig S1:** Summary statistics of the OneK1K data from four groups: test data, splatPop-simulated data, and scDesignPop-simulated data with both single-SNP (scDesignPop_single) and multi-SNP modes (scDesignPop_multi; details in Methods 10.2)). **(Top row)** Gene-level statistics: mean normalized expression, variance of normalized expression, proportion of non-zero counts per gene, and gene-gene correlation. **(Middle row)** Cell-level statistics: proportion of non-zero counts per cell, cell log library size, cell-cell correlation, and cell-cell distance. **(Bottom row)** Individual-level statistics: mean pseudobulk expression, variance of pseudobulk expression, individual log library size, and individual-individual distance. One cell in both scDesignPop-simulated data was excluded due to extreme outlier behavior. Values above each violin plot indicate the Kolmogorov-Smirnov (KS) statistic comparing simulated and test distributions; a lower KS value indicates closer resemblance between the two empirical distributions.

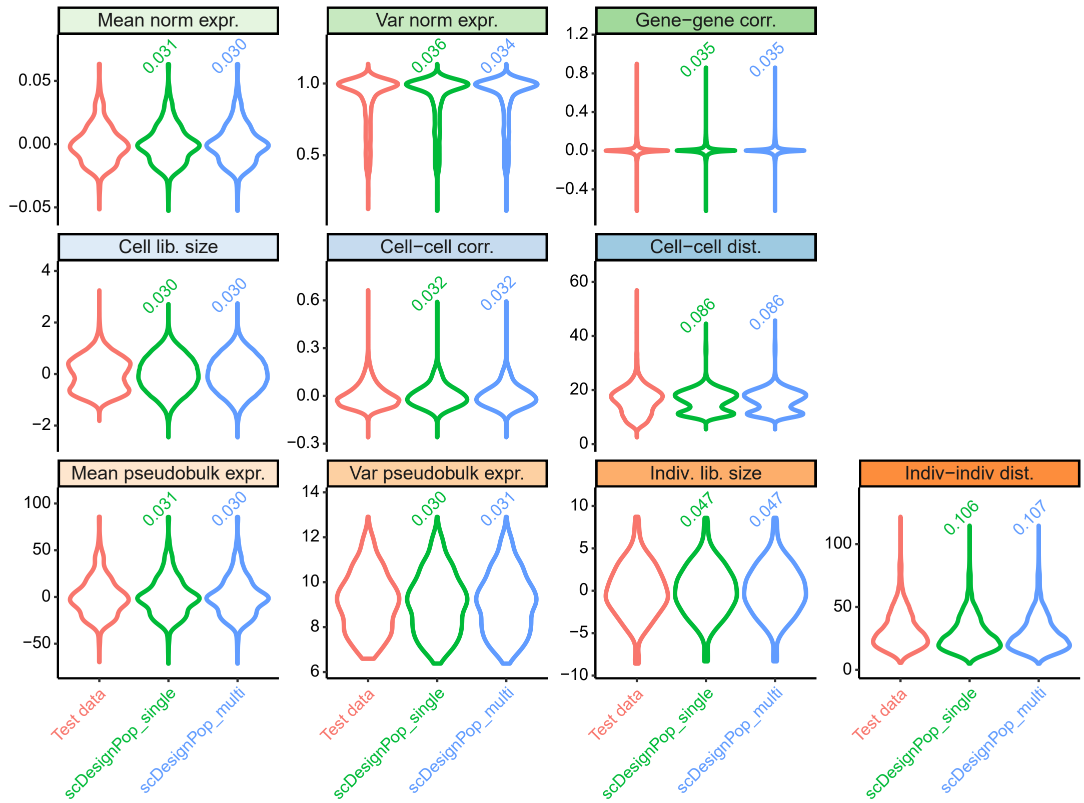


**Fig S2:** Summary statistics of the CLUES data from three groups: test data, and scDesignPop-simulated data with both single-SNP (scDesignPop_single) and multi-SNP modes (scDesignPop_multi; details in Methods 10.2). **(Top row)** Gene-level statistics: mean normalized expression, variance of normalized expression, and gene-gene correlation. **(Middle row)** Cell-level statistics: cell library size, cell-cell correlation, and cell-cell distance. **(Bottom row)** Individual-level statistics: mean pseudobulk expression, variance of pseudobulk expression, individual library size, and individual-individual distance. Values above each violin plot indicate the Kolmogorov–Smirnov (KS) statistic comparing simulated and test distributions; a lower KS value indicates closer resemblance between the two empirical distributions.

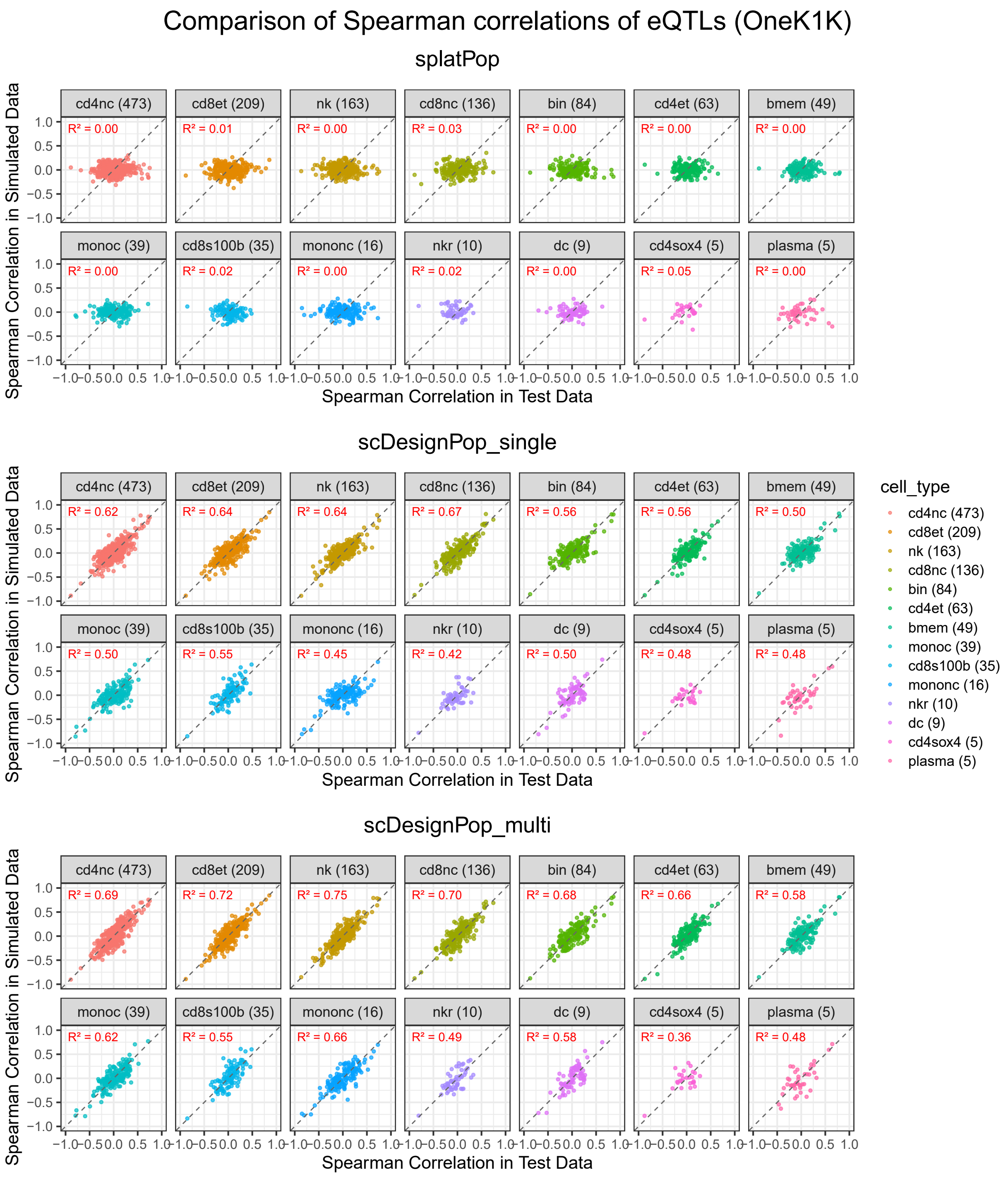


**Fig S3:** Scatterplots of estimated Spearman correlations in the test and simulated data for all cts-eQTLs for splatPop, scDesignPop_single, and scDesignPop_multi across 14 cell types. Each point corresponds to a Spearman correlation calculated between a gene's pseudobulk expression (aggregated via sum) in the simulated or test data and the corresponding eSNP genotypes in 100 individuals. $R^2$ was obtained from the Pearson correlation computed between the simulated and test data Spearman correlations. An average number of cells per individual (rounded) is shown for each cell type based on the OneK1K reference data (981 individuals).

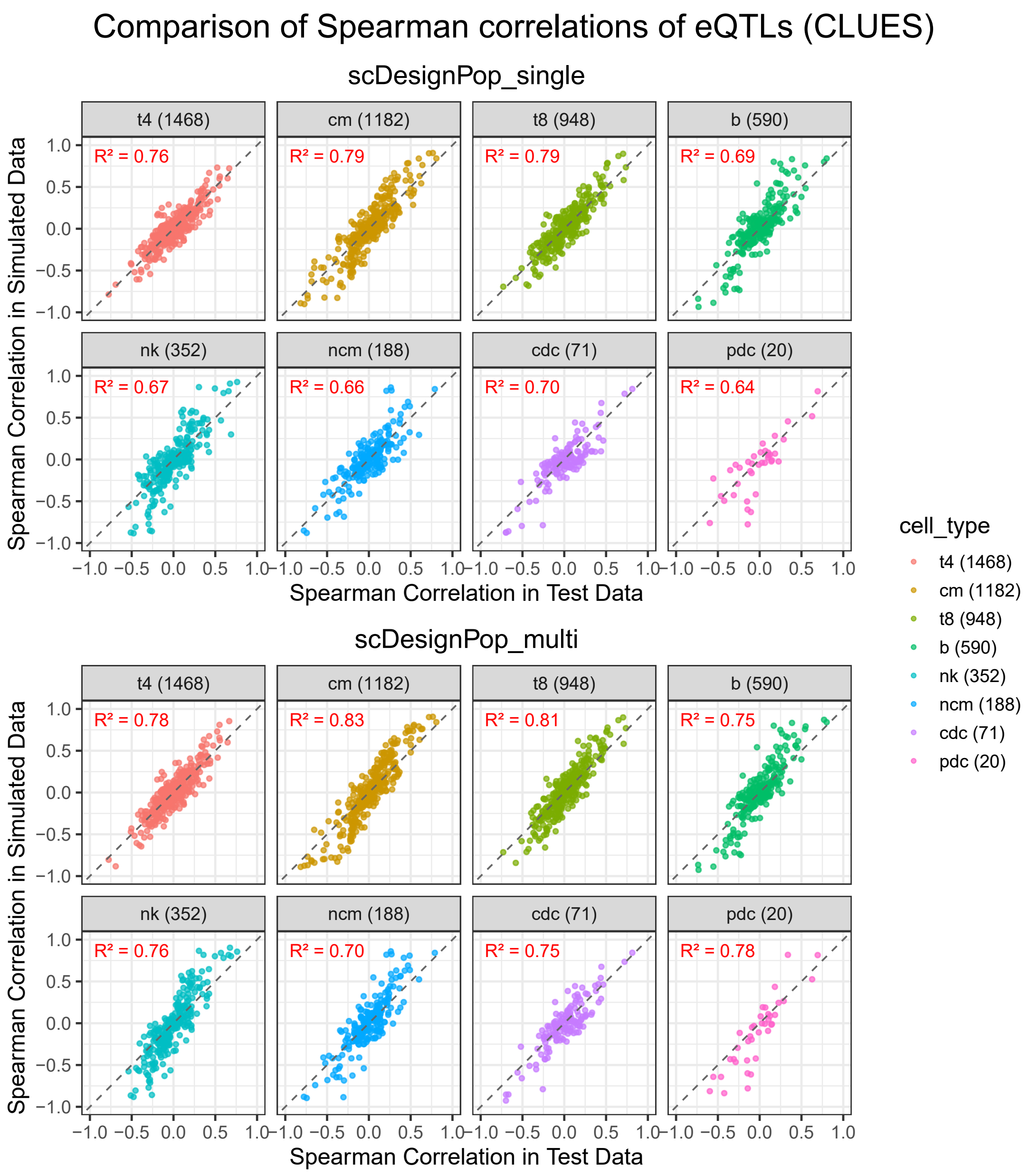


**Fig S4:** Scatterplots of estimated Spearman correlations in the test and simulated datasets for all cts-eQTLs for scDesignPop_single and scDesignPop_multi across 8 cell types. Each point corresponds to a Spearman correlation calculated between a gene's pseudobulk expression (aggregated via sum) in the simulated or test data and the corresponding eSNP genotypes in 150 individuals. $R^2$ was obtained from the Pearson correlation computed between the simulated and test data Spearman correlations. An average number of cells per individual (rounded) is shown for each cell type based on the CLUES reference data (256 individuals).

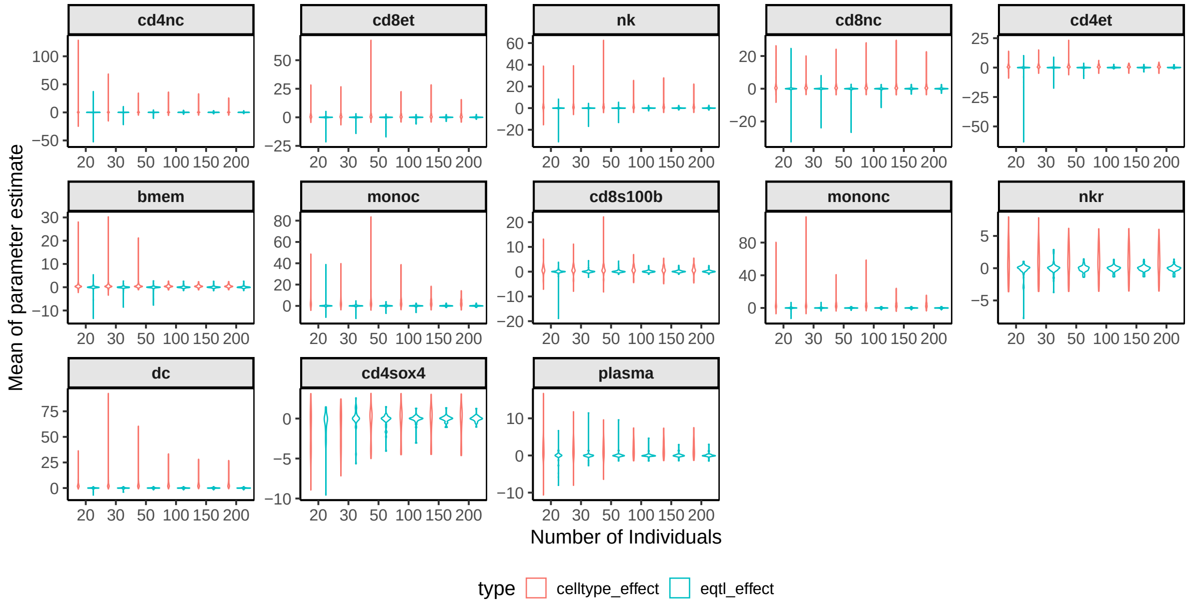


**Fig S5:** Mean of parameter estimates for cell type effect and cts-eQTL effect across varying numbers of individuals in training data in 13 cell types. The number of individuals at each setting was subsampled from OneK1K cohort 20 times and fitted using the same marginal model in scDesignPop. Means were computed across replicates for each gene-SNP pair and cell type. Immature/naive B cells (bin) were used as the baseline cell type in the marginal modeling, and are therefore not shown.

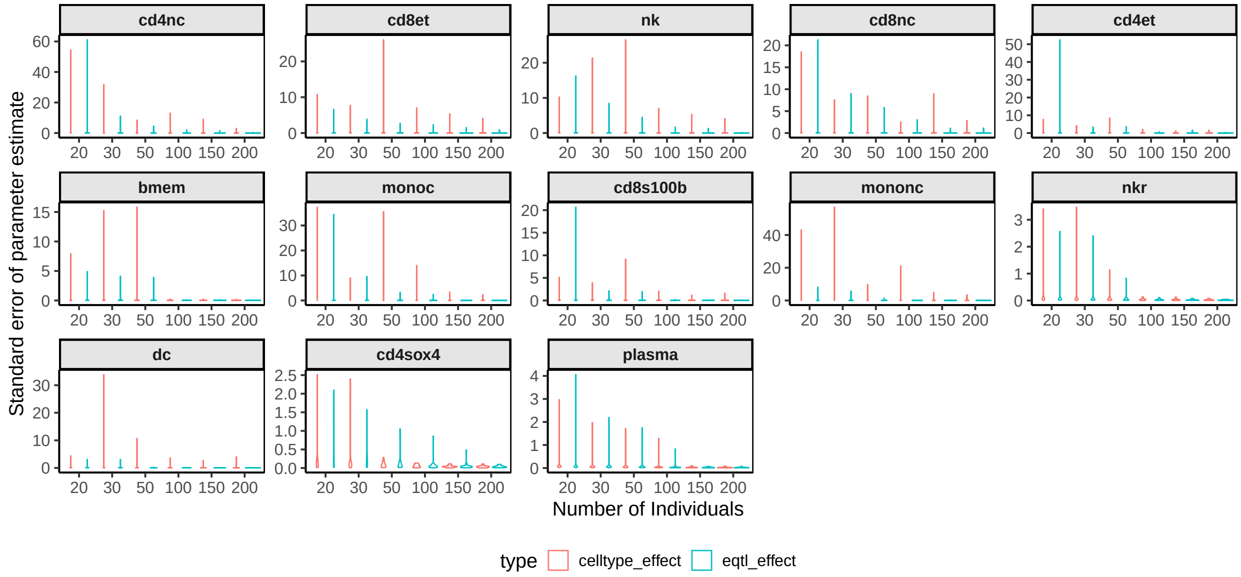


**Fig S6:** Standard error of parameter estimates for cell type effect and cts-eQTL effect across varying numbers of individuals in training data in 13 cell types. The number of individuals at each setting was subsampled from OneK1K cohort 20 times and fitted using the same marginal model in scDesignPop. Means were computed across replicates for each gene-SNP pair and cell type. Immature/naive B cells (bin) were used as the baseline cell type in the marginal modeling, and are therefore not shown.

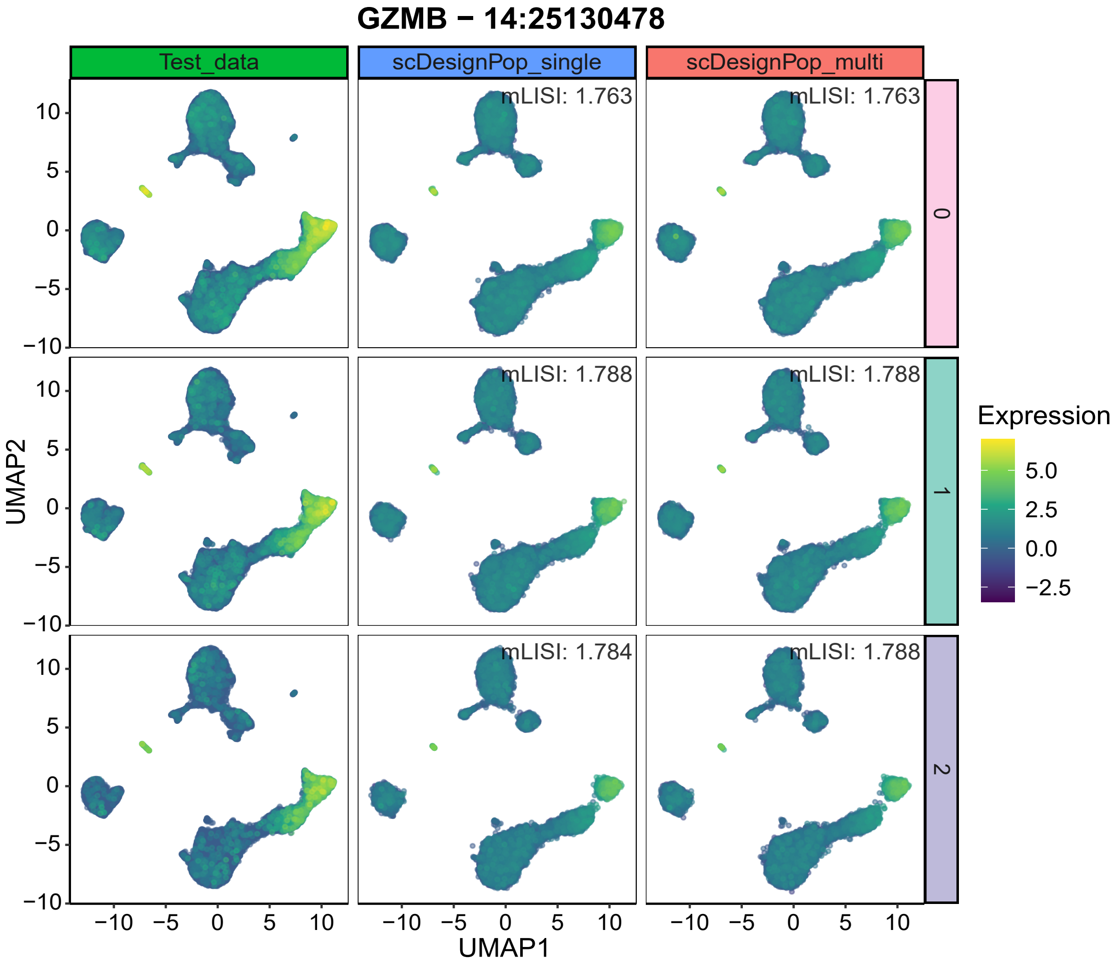


**Fig S7:** UMAP plot of CLUES cohort with *GZMB* gene's expression across 0, 1, 2 genotypes of SNP at locus 14:25130478 in three groups: test data, and scDesignPop-simulated data with both single-SNP (scDesignPop_single) and multi-SNP modes (scDesignPop_multi; details in Methods 10.2). mLISI values quantify similarity between simulated and test data within each genotype.

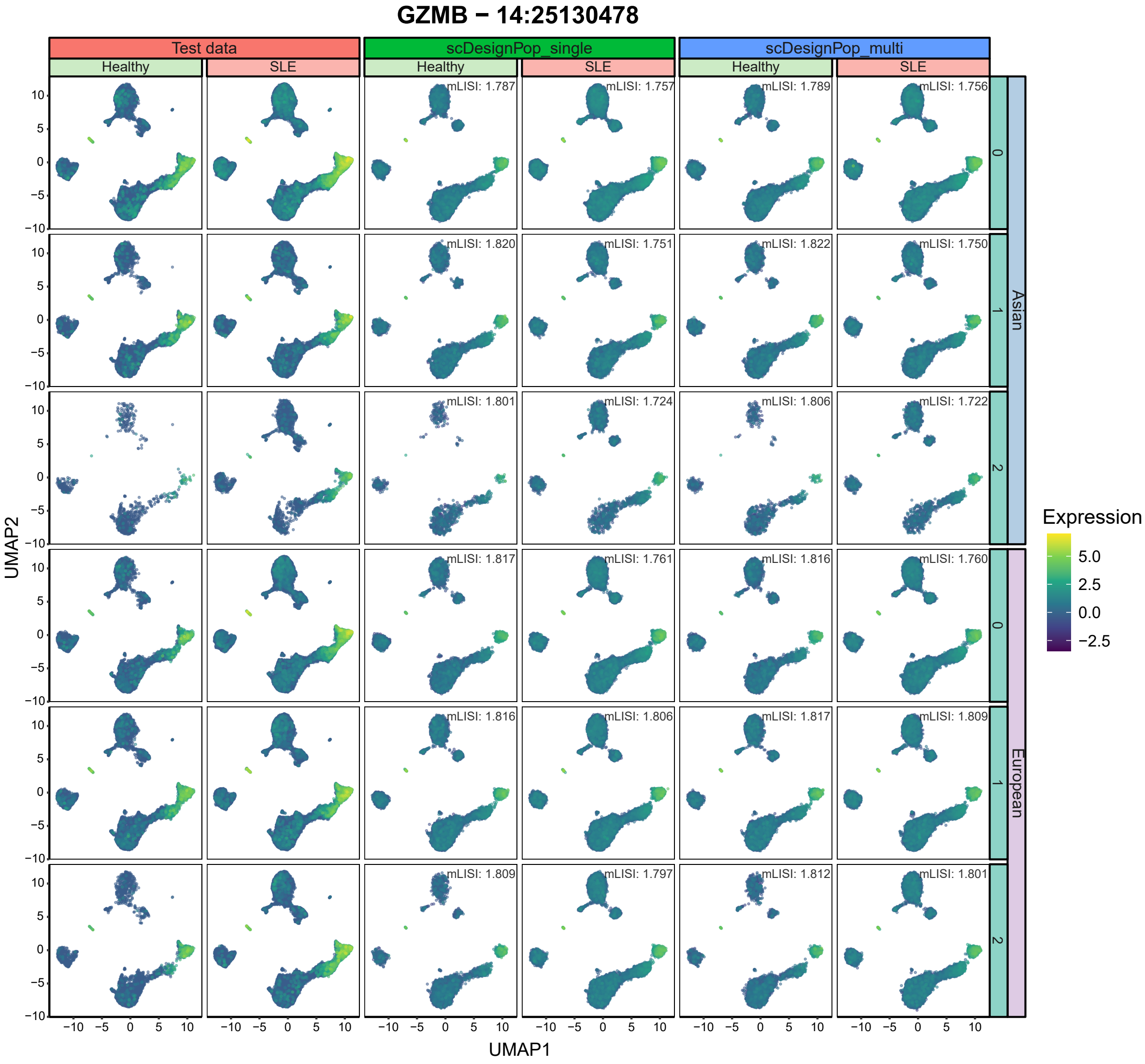


**Fig S8:** UMAP plot of CLUES data for *GZMB* gene's expression across 0, 1, 2 genotypes of SNP at locus 14:25130478 under different disease statuses and ancestries in three groups: test data, and scDesignPop-simulated data with both single-SNP (scDesignPop_single) and multi-SNP modes (scDesignPop_multi; details in Methods 10.2). mLISI values quantify similarity between simulated and test data within each genotype, disease status, and ancestry group.

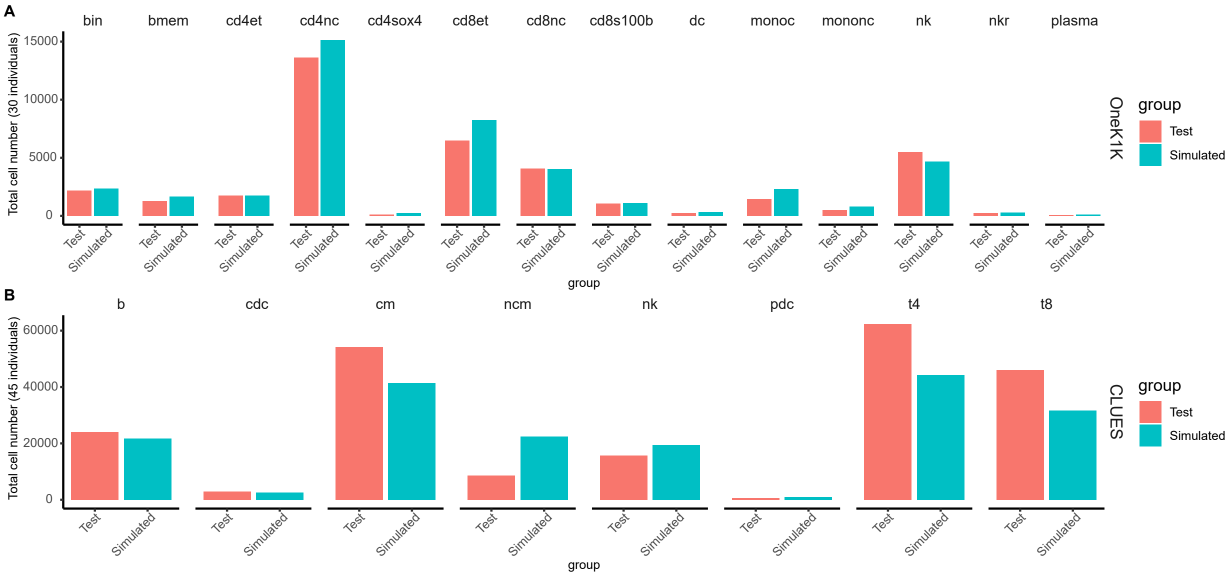


**Fig S9:** Bar charts of cell-type proportions in test and scDesignPop-simulated for the OneK1K and CLUES cohorts. **(A)** Total cell numbers across 14 cell types for 30 individuals in the OneK1K cohort, comparing test data (salmon) and scDesignPop-simulated data (teal). **(B)** Total cell numbers across 8 cell types for 45 individuals in the CLUES cohort, comparing test data (salmon) and scDesignPop-simulated data (teal).

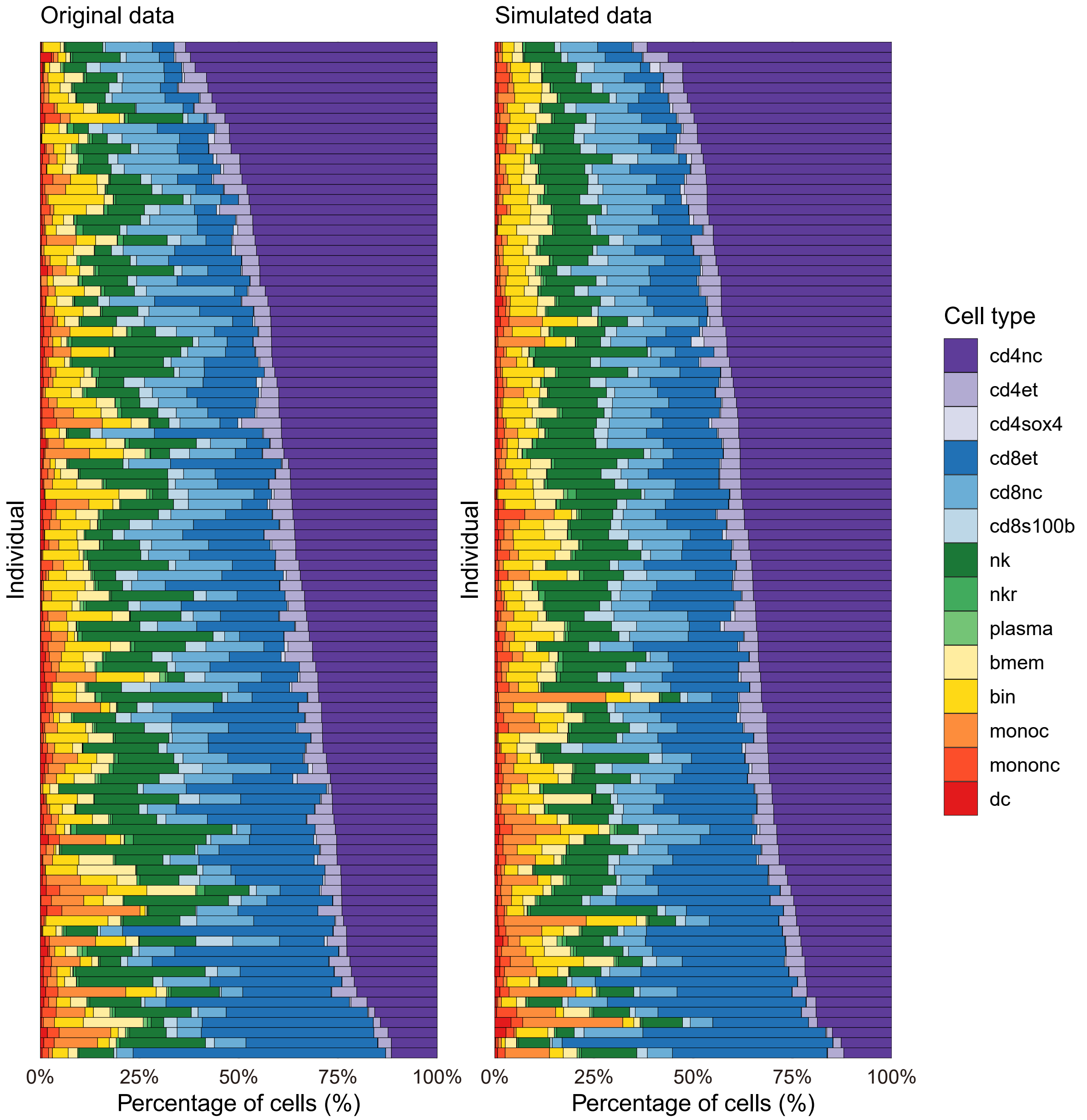


**Fig S10:** Comparison of cell-type proportions between reference and scDesignPop-simulated data for 100 randomly selected individuals (70 train and 30 test data) from the OneK1K cohort. Each horizontally stacked bar represents one individual. Individuals are ordered by decreasing percentage of $CD4_{NC}$ T cells (cd4nc).

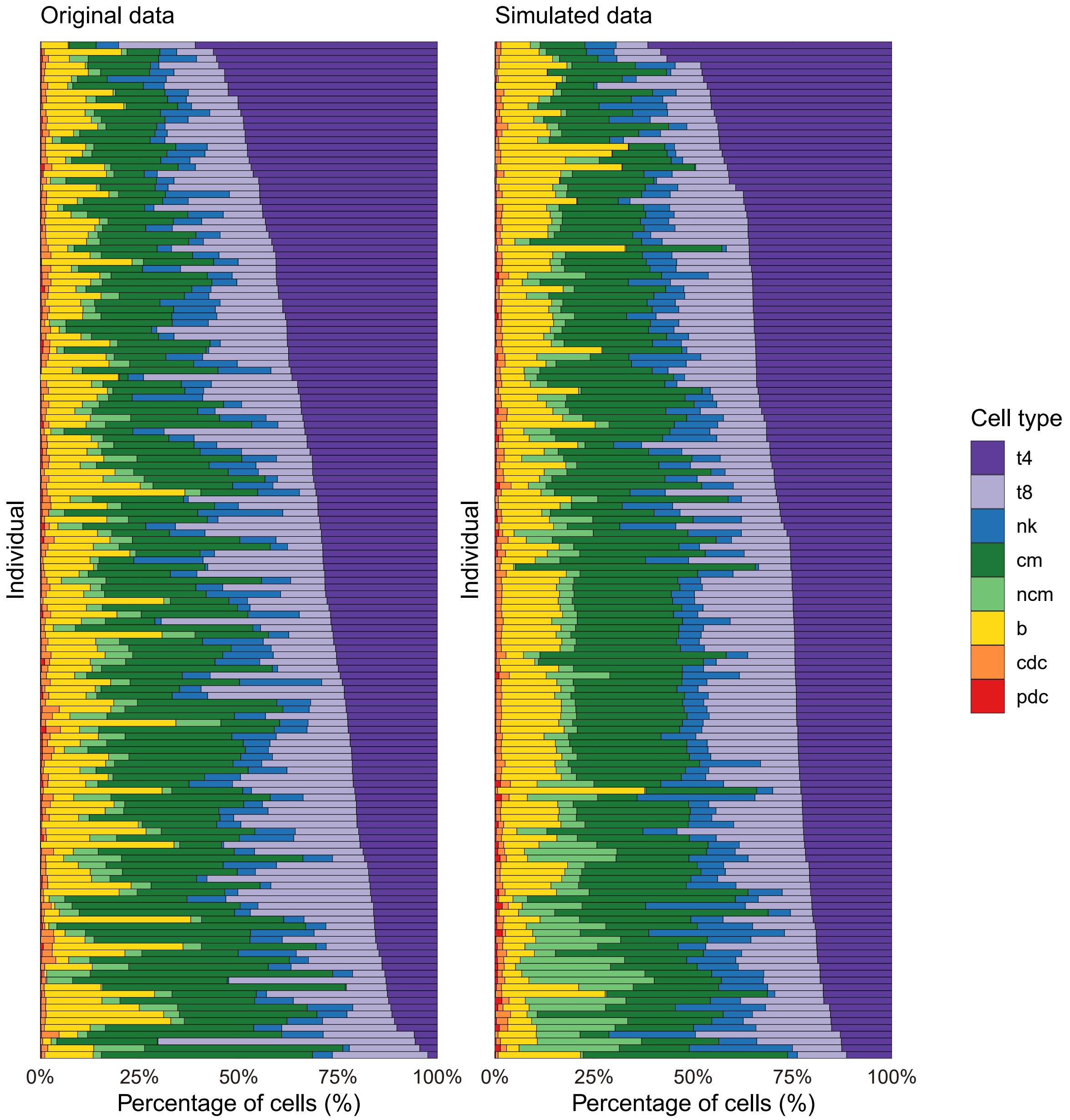


**Fig S11:** Comparison of cell-type proportions between reference and scDesignPop-simulated data for 150 randomly selected individuals (105 train and 45 test data) from the CLUES cohort. Each horizontally stacked bar represents one individual. Individuals are ordered by decreasing percentage of CD4 T cells (t4).

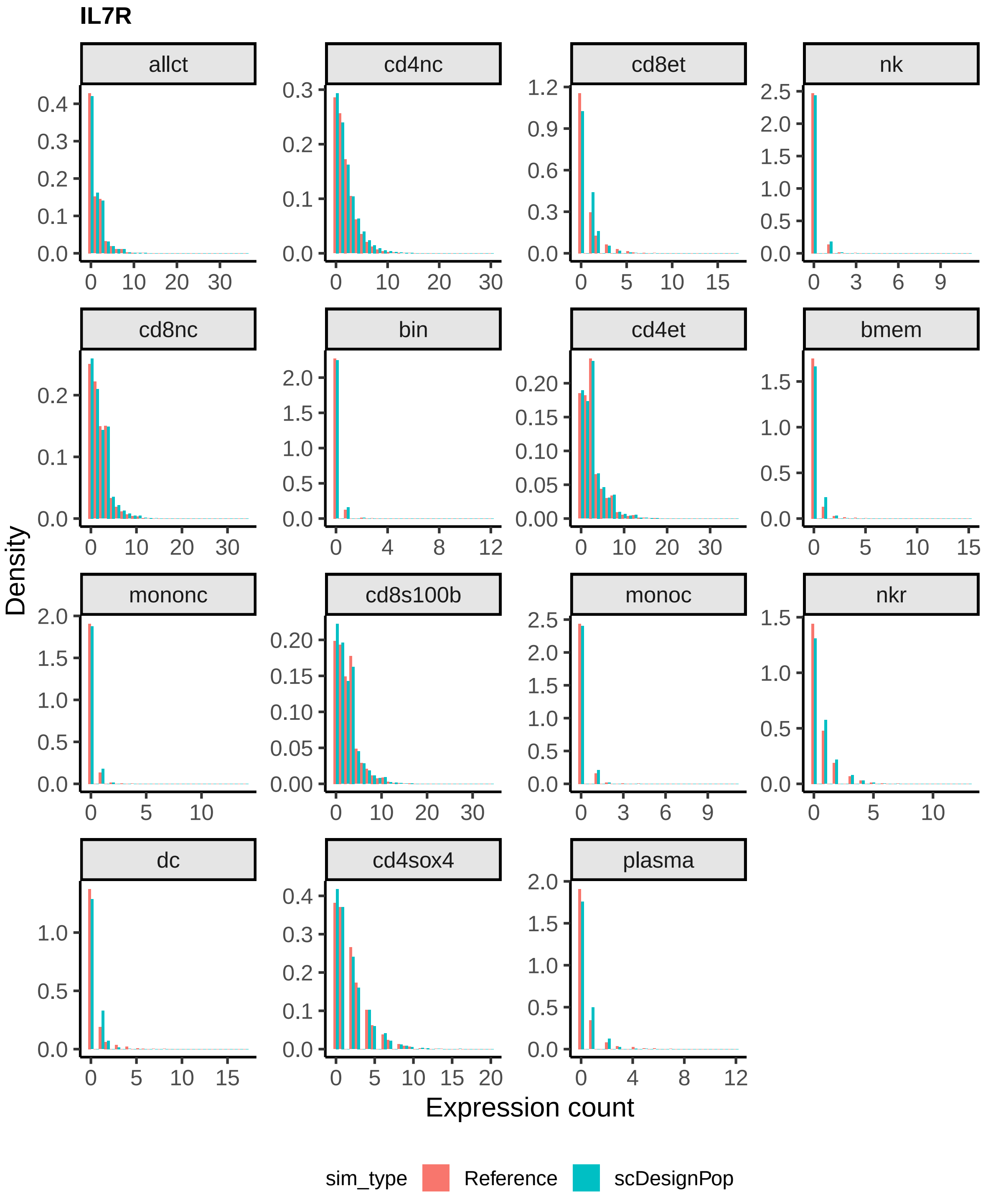


**Fig S12:** Density histograms of *IL7R* expression counts across all cells (allct) and 14 cell types, comparing the reference OneK1K cohort (salmon) and scDesignPop-simulated scRNA-seq data (teal). A total of 1,267,768 cells from 981 individuals in the reference dataset were compared with 1,316,454 synthetic cells generated by scDesignPop for 982 individuals

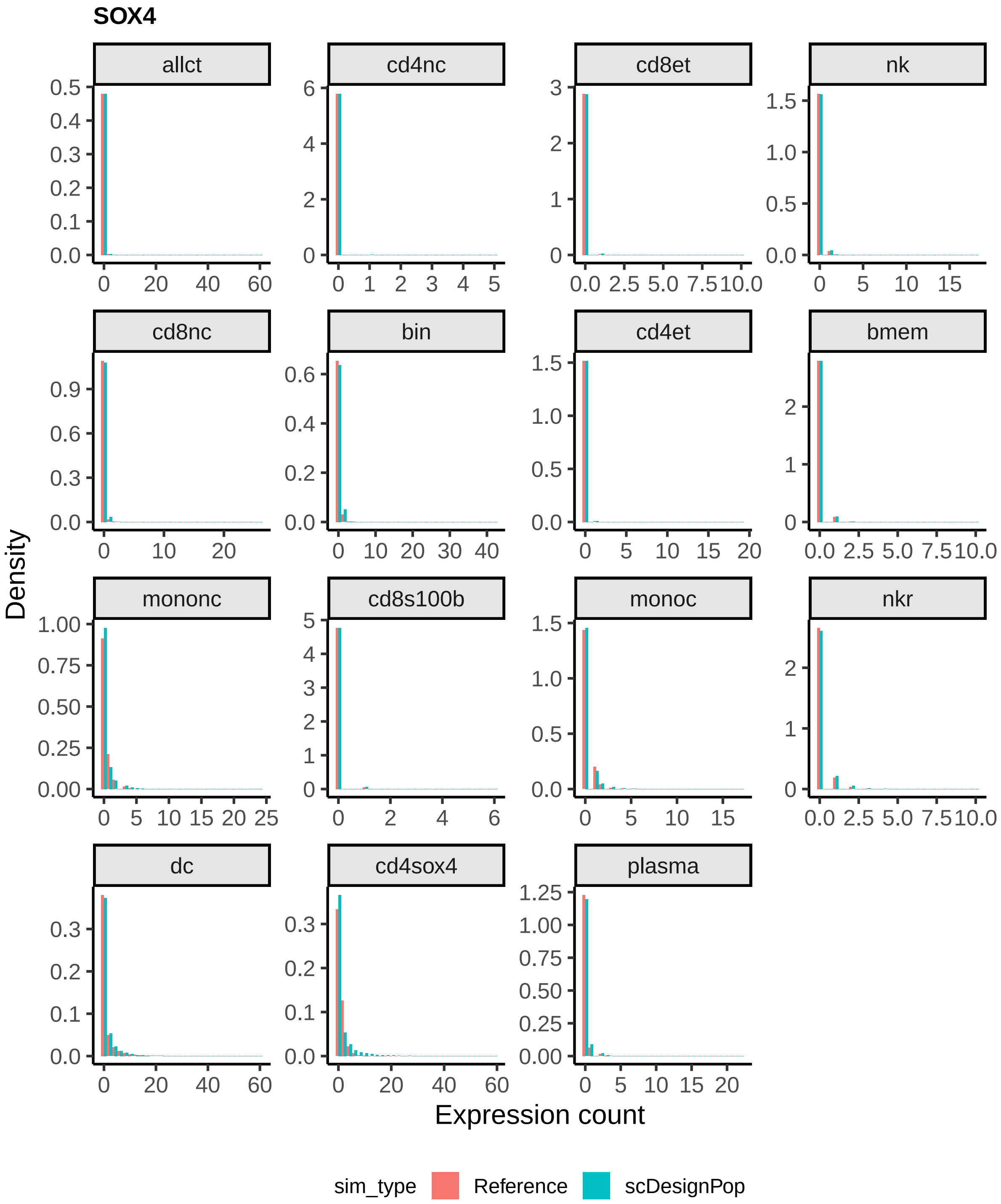


**Fig S13:** Density histograms of *SOX4* expression counts across all cells (allct) and 14 cell types, comparing the reference OneK1K cohort (salmon) and scDesignPop-simulated scRNA-seq data (teal). A total of 1,267,768 cells from 981 individuals in the reference dataset were compared with 1,316,454 synthetic cells generated by scDesignPop for 982 individuals.

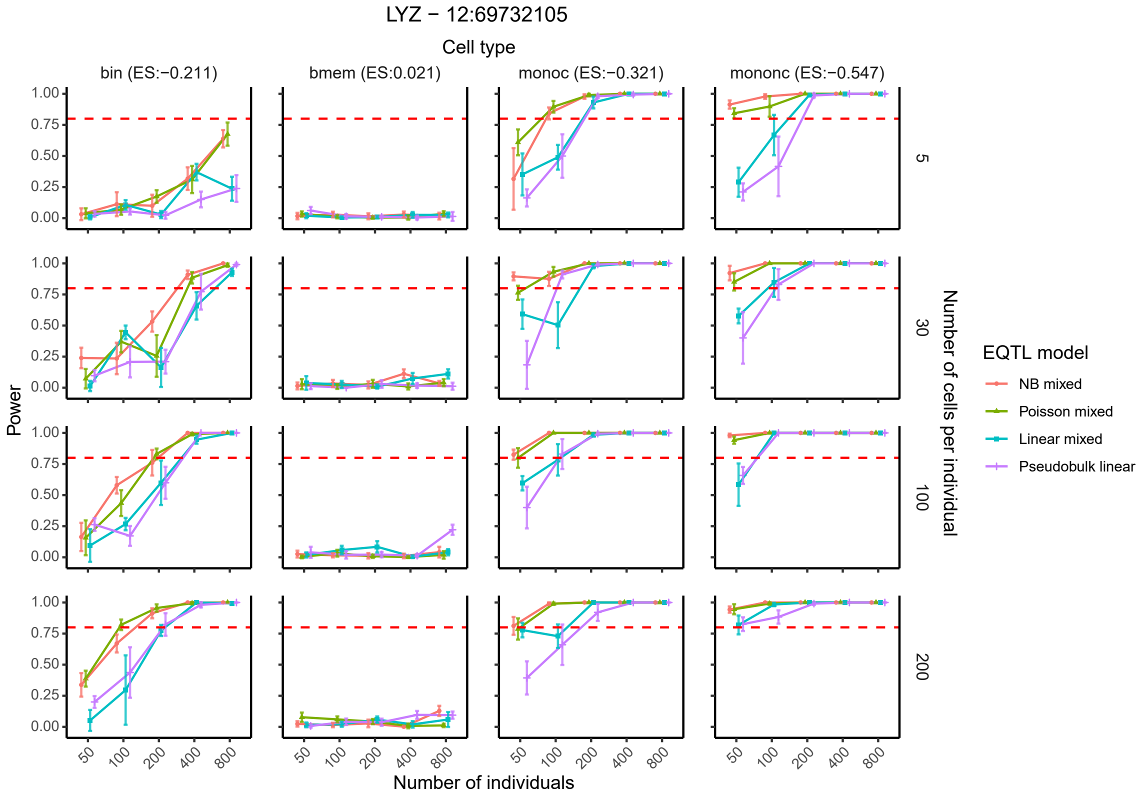


**Fig S14:** Power curves for detecting the cts-eQTL effect between *LYZ* gene and SNP locus 12:69732105 across varying numbers of individuals and cells per individual in four representative cell types. Four eQTL models (negative binomial mixed, Poisson mixed, linear mixed, and pseudobulk linear) were used. Error bars represent standard deviations. Effect sizes estimated from the marginal full model are shown above each panel. The red dashed line indicates 80% power. *Abbreviations:* immature/naive B cells (bin); memory B cells (bmem); classical monocytes (monoc); nonclassical monocytes (mononc); effect size (ES); negative binomial (NB).

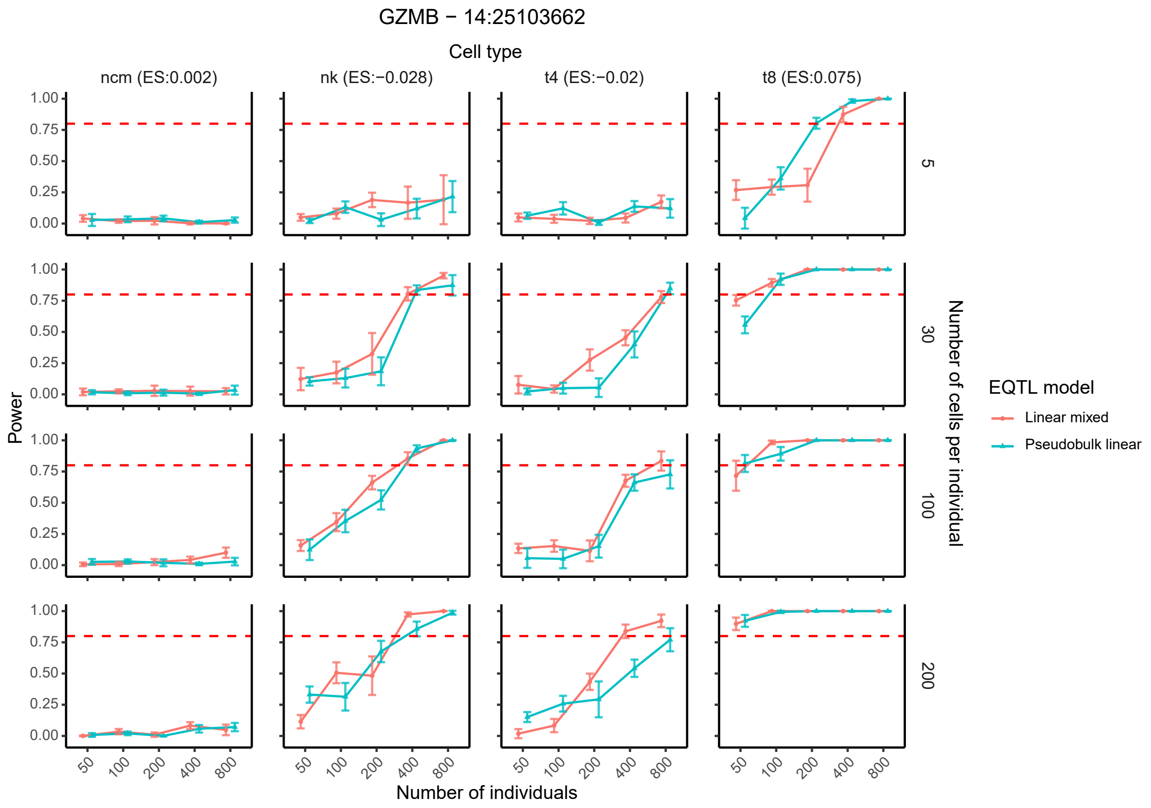


**Fig S15:** Power curves for detecting the cts-eQTL effect between *GZMB* gene and SNP locus 14:25103662 across varying numbers of individuals and cells per individual in four representative cell types. Two eQTL models (linear mixed and pseudobulk linear) were used. Negative binomial and Poisson mixed eQTL models were excluded since CLUES data is normalized (e.g. not count-based). Error bars denote standard deviations. Effect sizes estimated from the marginal full model are shown above each panel. The red dashed line indicates 80% power. *Abbreviations:* nonclassical monocytes (ncm); natural killer cells (nk); CD4+ T cells (t4); and CD8+ T cells (t8); effect size (ES).

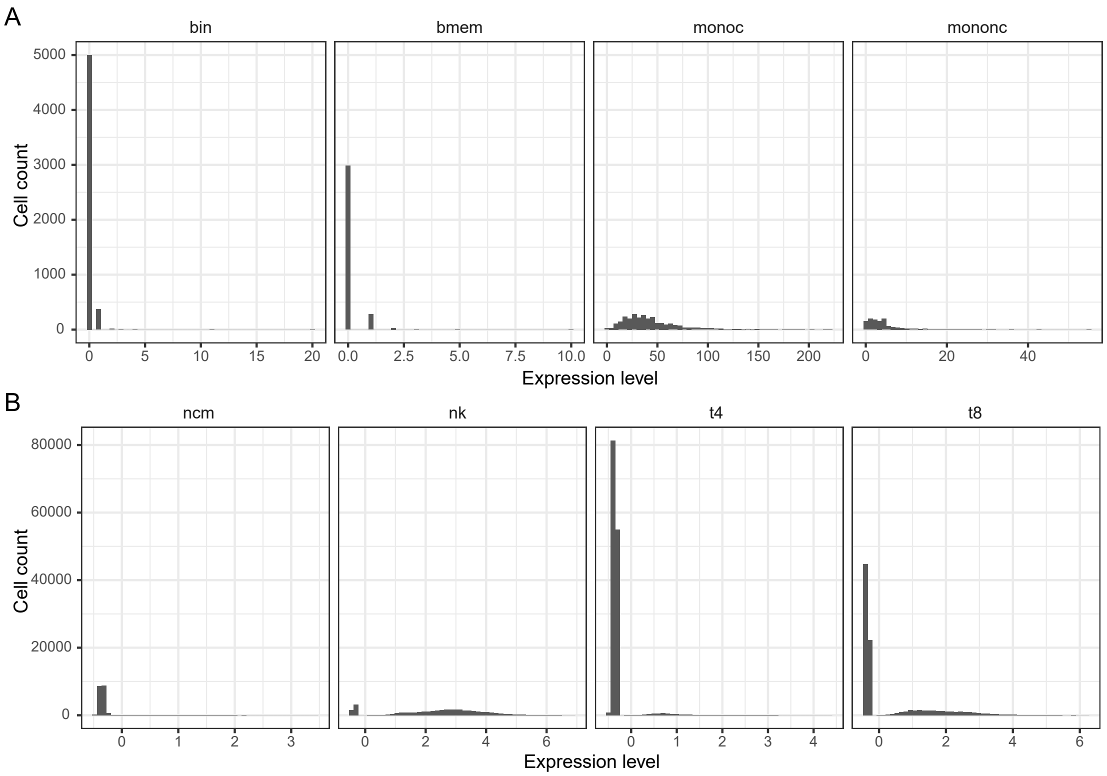


**Fig S16:** Histograms of gene expression levels across cell types showing **(A)** distribution of *LYZ*'s expression across four cell types in OneK1K training data; and **(B)** distribution of *GZMB*'s expression across four cell types in the CLUES training data. *Abbreviations:* immature/naive B cells (bin); memory B cells (bmem); classical monocytes (monoc); nonclassical monocytes (mononc, ncm); natural killer cells (nk); CD4+ T cells (t4); CD8+ T cells (t8).

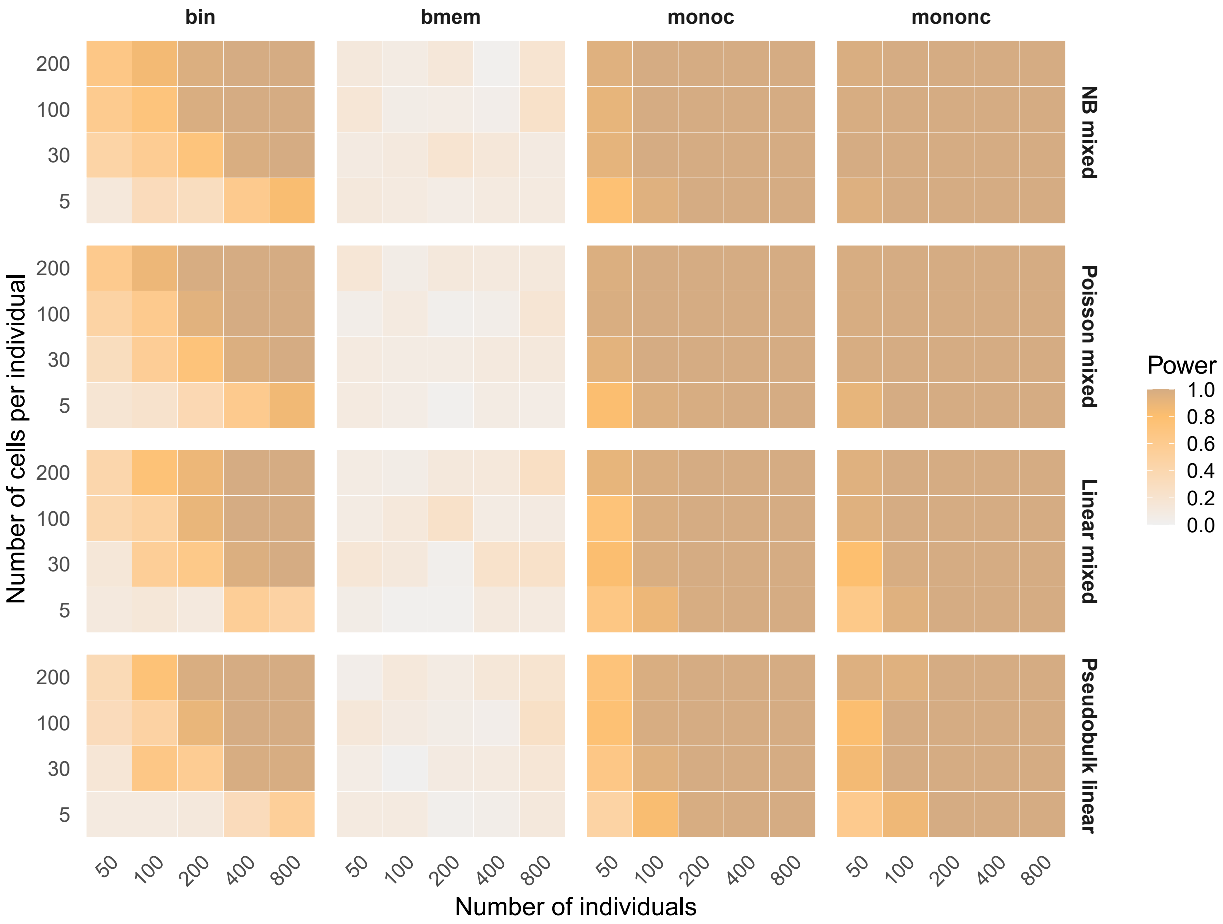


**Fig S17:** Heatmaps showing power of detecting the cts-eQTL effect between *LYZ* gene and SNP locus 12:69732105 using four different eQTL models (Negative Binomial mixed, Poisson mixed, linear mixed and pseudobulk linear) in immature/naive B cells (bin), memory B cells (bmem), classical monocytes (monoc), and nonclassical monocytes (mononc) from the OneK1K data.

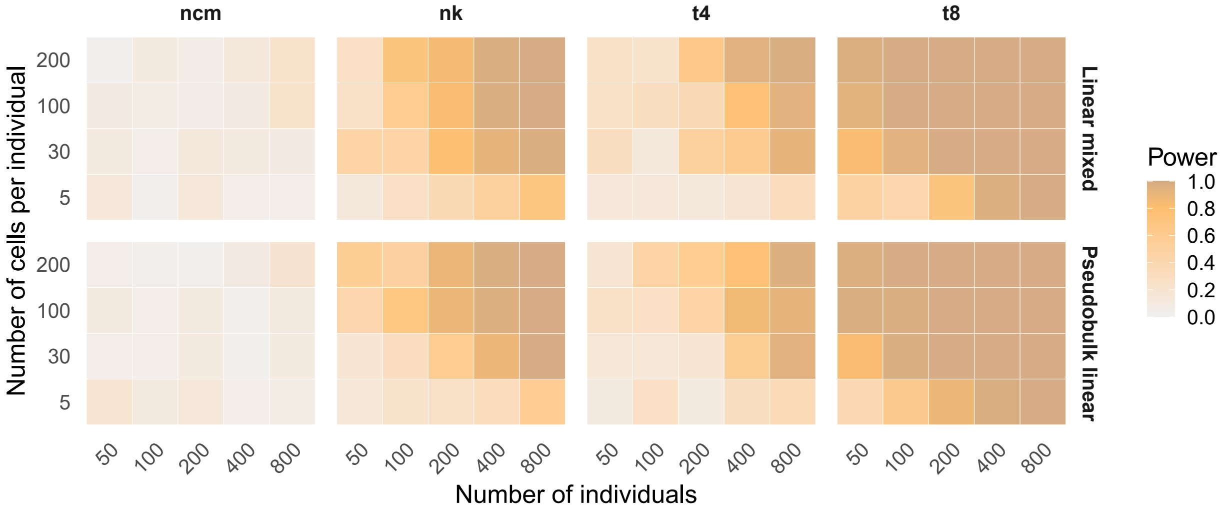


**Fig S18:** Heatmaps showing power of detecting the cts-eQTL effect between *GZMB* gene and SNP locus 14:25103662 using two different eQTL models (linear mixed and pseudobulk linear) in nonclassical monocytes (ncm), natural killer cells (nk), CD4+ T cells (t4), and CD8+ T cells (t8) from the CLUES data.

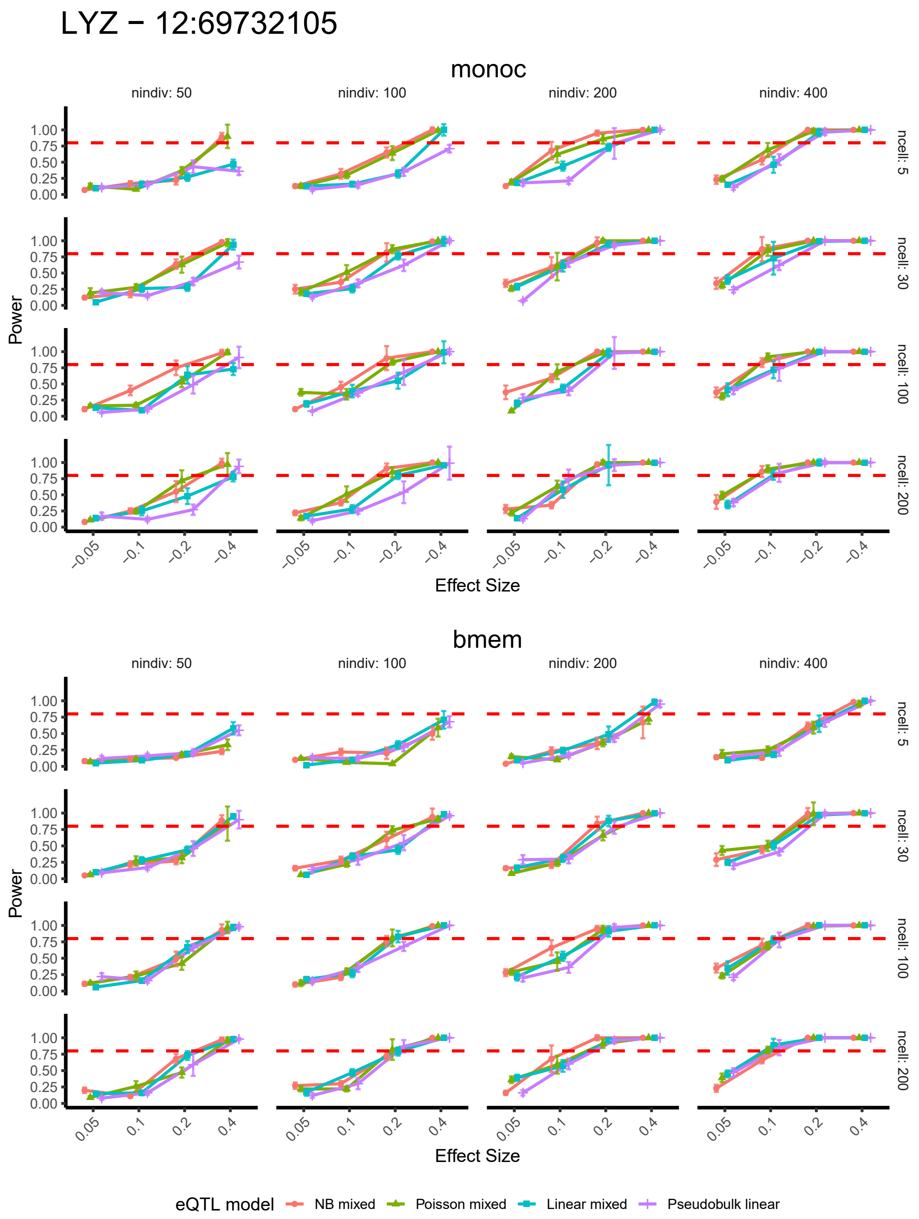


**Fig S19:** Power curves for detecting the cts-eQTL effect between *LYZ* gene and SNP locus 12:69732105 under user-specified cts-eQTL effect sizes in classical monocytes (monoc) and memory B cells (bmem) from the OneK1K data. Four eQTL models (Negative Binomial mixed, Poisson mixed, linear mixed and pseudobulk linear) are used. Error bars represent standard deviations. scDesignPop's estimated Power will increase with a higher absolute value of user-specified effect sizes. The red dashed line indicates 80% power.

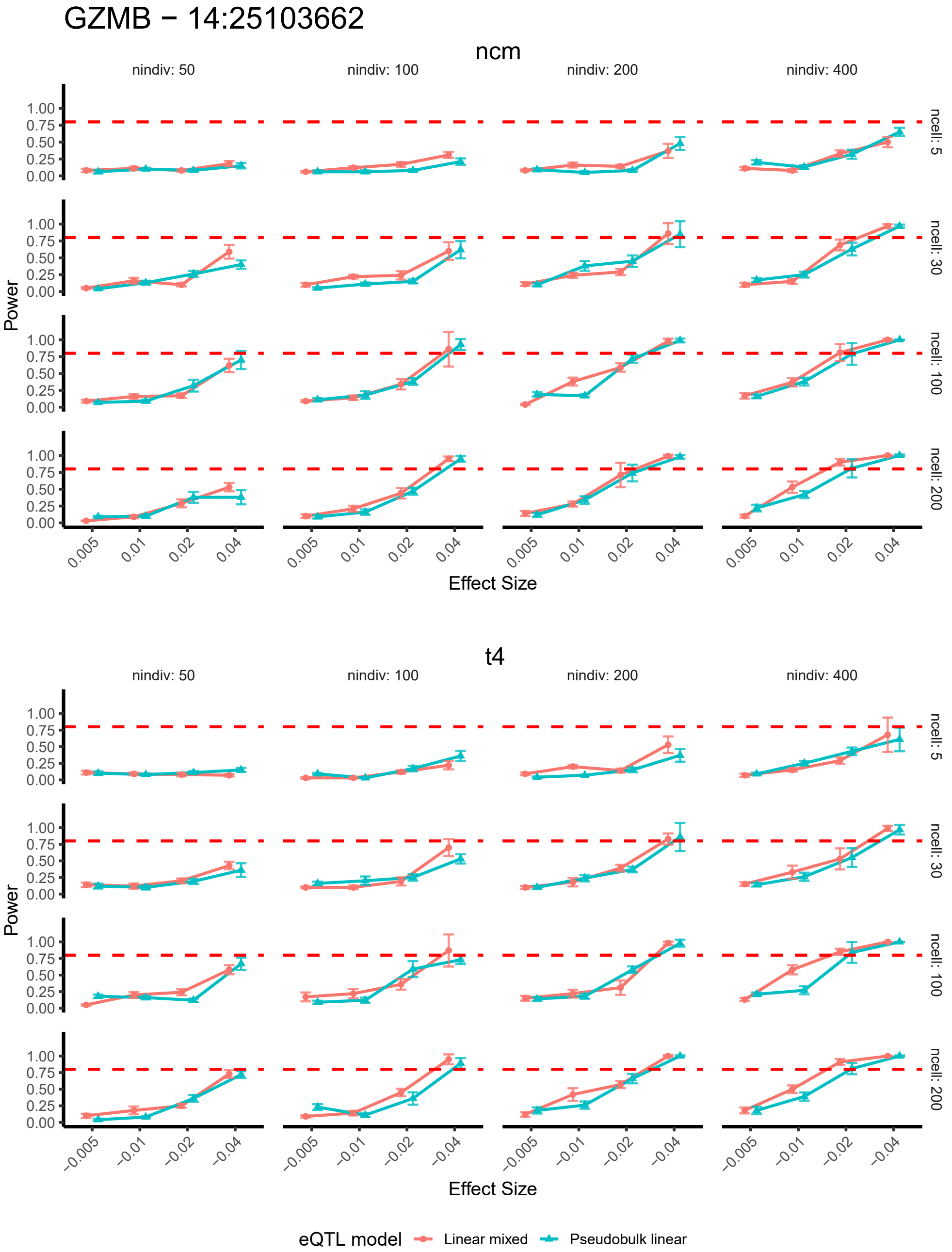


**Fig S20:** Power curves for detecting the eQTL effect between *GZMB* gene and SNP locus 14:25103662 under user-specified cts-eQTL effect sizes in nonclassical monocytes (ncm) and CD4+ T cells (t4) from the CLUES data. Two eQTL models (linear mixed and pseudobulk linear) are used. Error bars represent standard deviations. scDesignPop's estimated Power will increase with a higher absolute value of user-specified effect sizes. The red dashed line indicates 80% power.

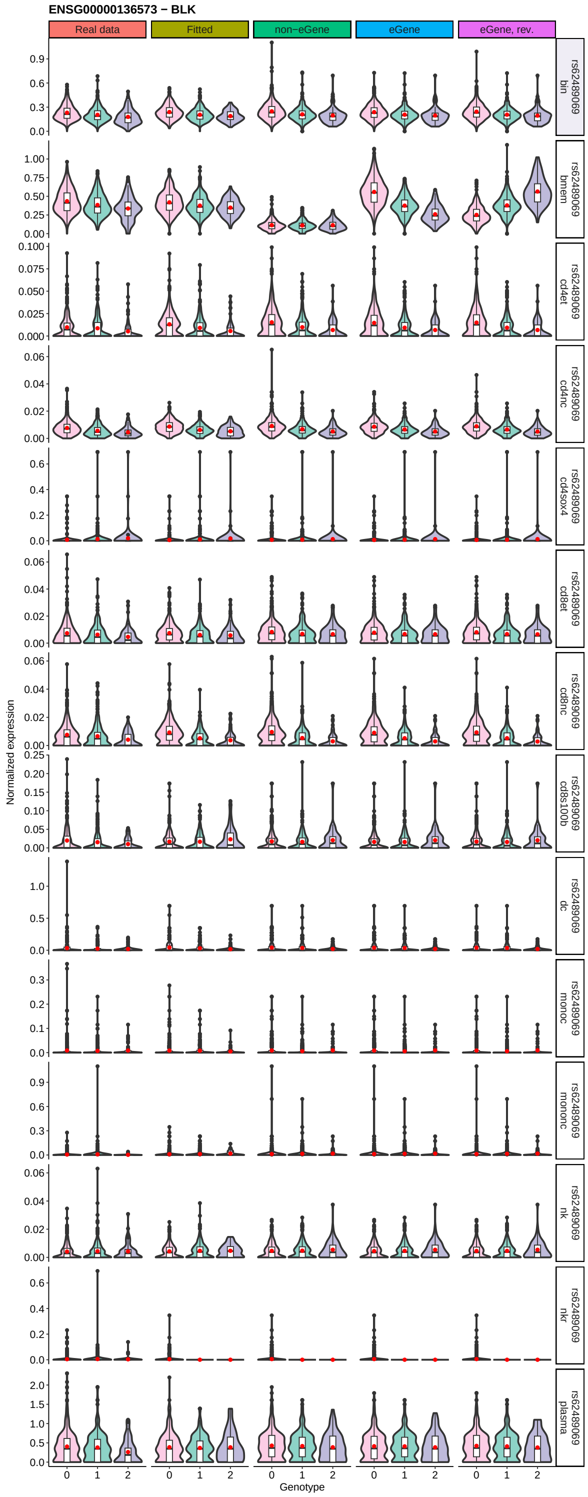


**Fig S21:** Violin plots of pseudobulk expression for *BLK* gene across SNP (rs62489069) genotypes in 14 cell types comparing real OneK1K data with scDesignPop-simulated data generated under fitted parameters or modified settings defining either a non-eGene or eGene in memory B cells. In the non-eGene setting, the eQTL effect size was set to zero and the gene's mean expression was decreased by 2 $\log_2$ fold-change. In the eGene settings, the eQTL effect size was increased by 2 $\log_2$ fold-change, with its direction either matching the same sign as the fitted or reversed (eGene, rev.).

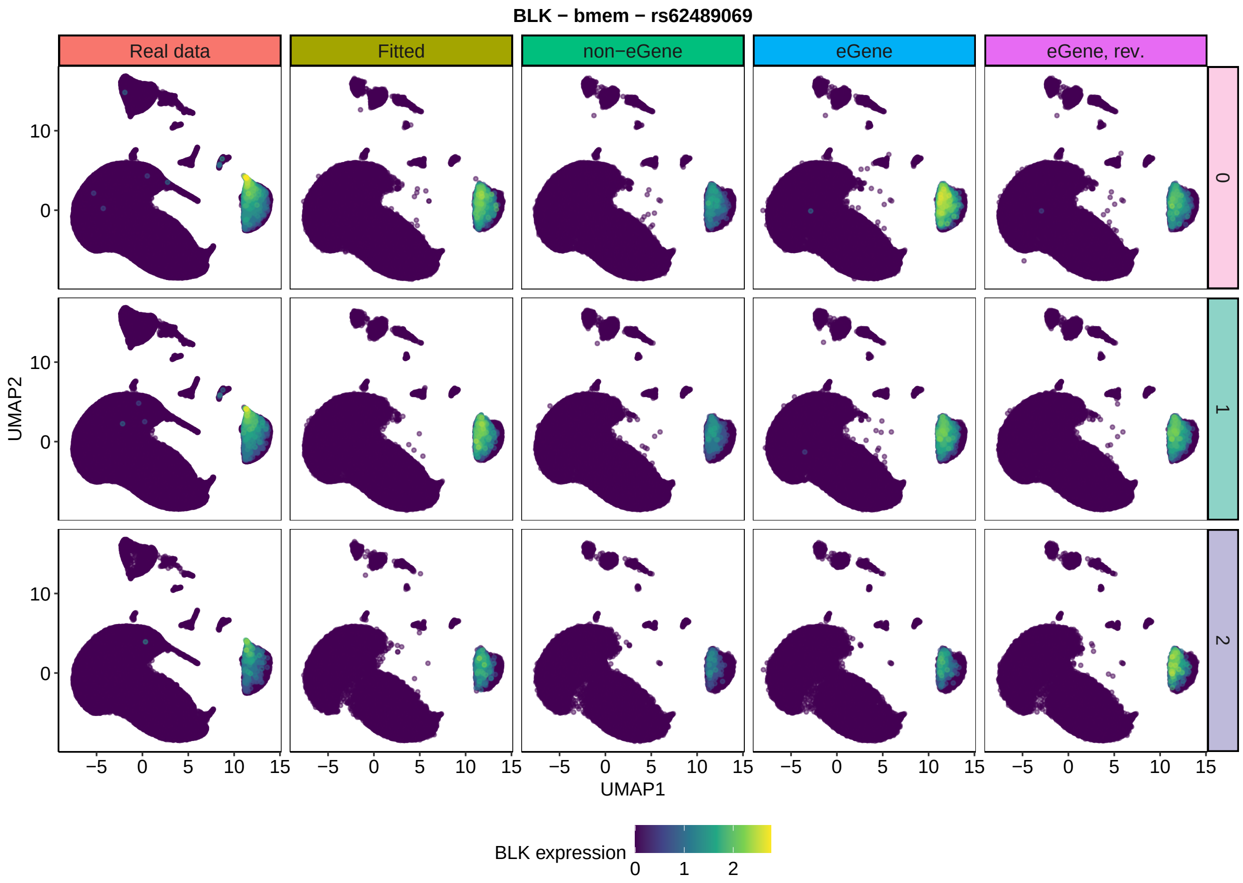


**Fig S22:** UMAP plots of *BLK* gene expression across genotypes at SNP locus *rs62489069*, comparing real OneK1K data with scDesignPop-simulated data generated under fitted parameters or modified settings defining either a non-eGene or eGene. In the non-eGene setting, the eQTL effect size was set to zero and the gene's mean expression was decreased by 2 $\log_2$ fold-change. In the eGene settings, the eQTL effect size was increased by 2 $\log_2$ fold-change, with its direction either matching the same sign as the fitted or reversed (eGene, rev.). Only *BLK* expression in 48,023 memory B cells out of 1,267,768 total cells per setting is highlighted. *Abbreviations*: memory B cells (bmem)

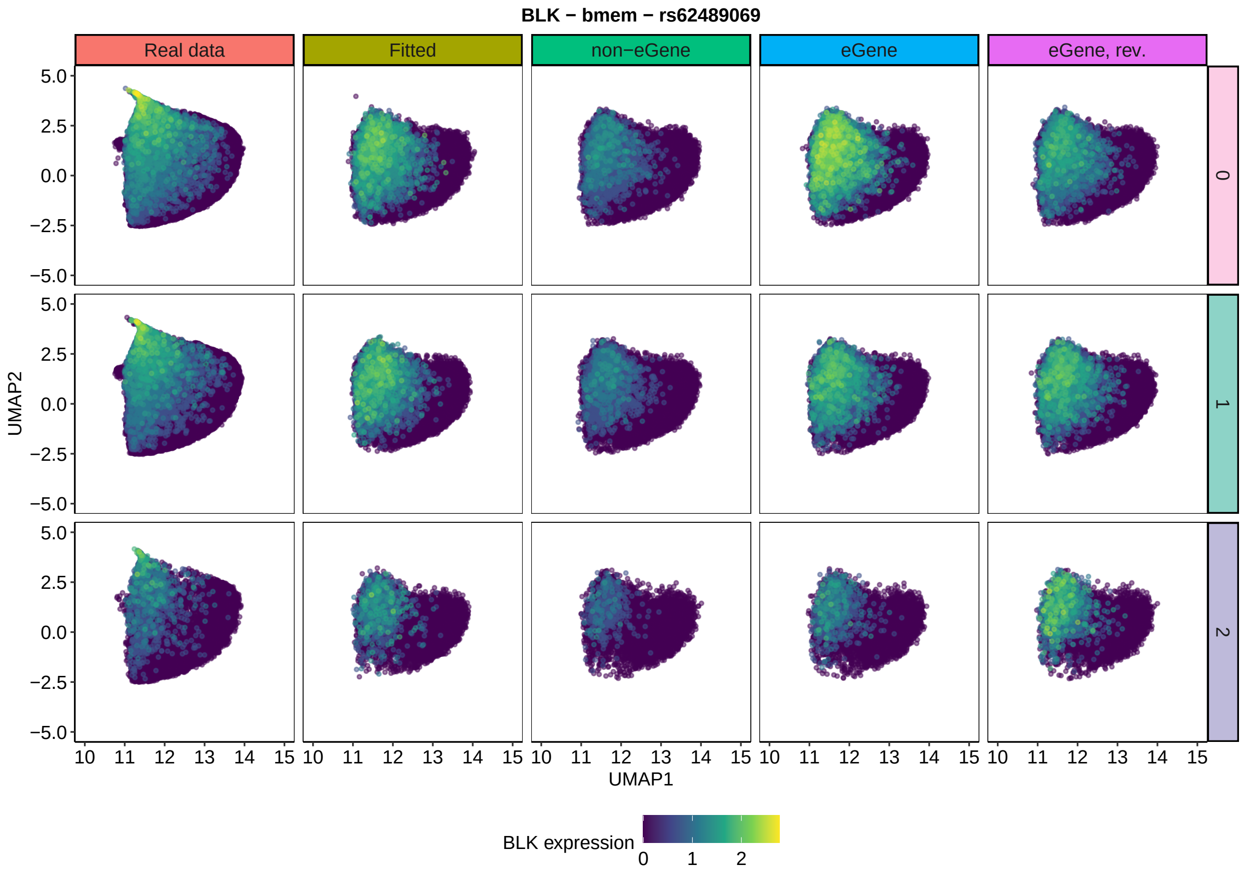


**Fig S23:** Zoomed-in UMAP plots of B cells highlighting *BLK* gene expression across genotypes at SNP locus *rs62489069*, comparing real OneK1K data with scDesignPop-simulated data generated under fitted parameters or modified settings defining either a non-eGene or eGene (full UMAP shown in S22). In the non-eGene setting, the eQTL effect size was set to zero and the gene's mean expression was decreased by 2 $\log_2$-fold-change. In the eGene settings, the eQTL effect size was increased by 2 $\log_2$ fold-change, with its direction either matching the same sign as the fitted or reversed (eGene, rev.). Only *BLK* expression in 48,023 memory B cells out of 130,091 total B cells per setting is highlighted. *Abbreviations*: memory B cell (bmem).

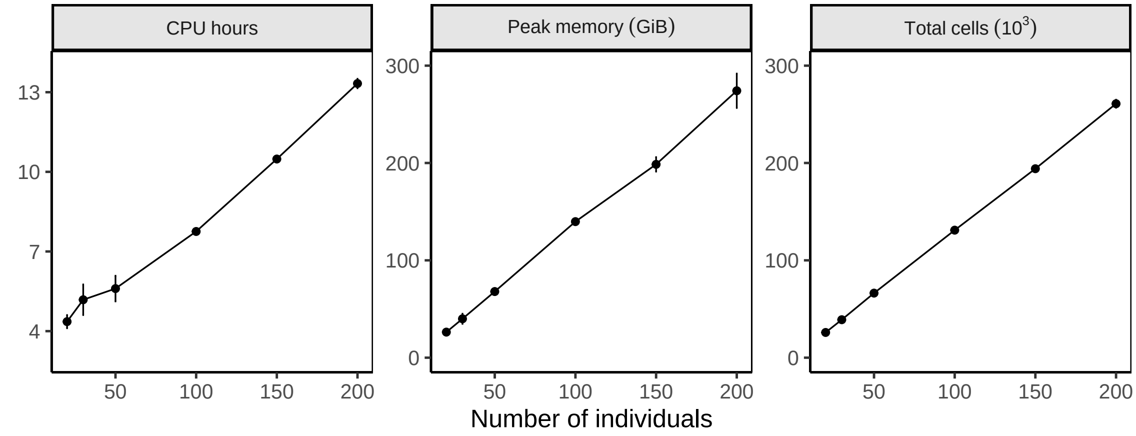


**Fig S24:** Overall computation time and peak memory usage of scDesignPop scales linearly. An end-to-end pipeline consisting of data input construction, marginal model fitting, copula model fitting, and synthetic data generation was used to evaluate scDesignPop's total computation time in CPU hours and peak memory usage in GiB. With scDesignPop's parallelization, 20 CPU cores were used to train the marginal models, and 2 CPU cores were used to train the copula and for generating synthetic data for 810 genes. After subsampling the number of individuals from OneK1K cohort, the same number of cells was used for both model training and synthetic data generation. Each point represents the mean of three replicates, with error bars indicating the standard deviation. *Abbreviations:* central processing unit (CPU); gibibyte (GiB).

**Algorithm 1:** Power calculation for single-cell eQTL analysis

**Input Parameters:** Gene $j$, SNP index $l$, eQTL model method, cell-type index $h \in \{1, \ldots, H\}$, number of individuals $\tilde{K}$ indexed by $\tilde{k}$, number of cells per individual $n_{\tilde{k}}$ constant for all $\tilde{k} \in \{1, \ldots, \tilde{K}\}$, significance threshold $\alpha$, number of SNPs $N_{\text{SNP}}$, number of genes $N_{\text{gene}}$, and simulation iterations $B$

**Input Data:** A cell-by-gene matrix $\mathbf{Y} = \{Y_{ij}\}$, a cell-by-covariate design matrix $\mathbf{C}$, a cell-state matrix $\mathbf{S}$ with $H$ discrete cell types, a cell-by-individual indicator matrix $\mathbf{Z}$, and an individual-by-SNP genotype matrix $\mathbf{G}$ with $K$ individuals and $L$ SNPs

**Output:** A vector of average statistical powers $\mathbf{P} = \{P_h\}_{h=1}^{H}$ across cell types

Extract the individual-level genotype $\mathbf{G}_{l(j)} = [\mathbf{G}_{1,l(j)}, \ldots, \mathbf{G}_{K,l(j)}]^\top$ of the input SNP $l$ from $\mathbf{G}$ for gene $j$, and map it to the cell level using indicator matrix $\mathbf{Z}$ ;
Extract the gene $j$'s expression vector $\mathbf{y}_j = (Y_{1j}, \ldots, Y_{Ij})^\top$ from the cell-by-gene matrix $\mathbf{Y}$ ;
Fit the marginal model introduced in Equation 1 for gene $j$ across cell types to obtain the full model with the true genotype effect size $\beta_{jh}$ (genotype main effect and its interaction effect) for each cell type $h$ ;
For each cell type $h$, change the $\beta_{jh}$ to zero to obtain the null model ;
Let $\mathbf{C}_h$ denote the submatrix of $\mathbf{C}$ whose cells belong to cell type $h$ in the $\mathbf{S}$;
**for** $b = 1$ **to** $B$ **do**
    $\tilde{\mathbf{Z}}^{(b)} \leftarrow$ sample $\tilde{K}$ individuals from $\mathbf{Z}$ ;
    $\tilde{\mathbf{G}}_{l(j)}^{(b)} \leftarrow$ sample $\tilde{K}$ individual-level genotype covariate from $\mathbf{G}_{l(j)}$ ;
    $\tilde{\mathbf{C}}_h^{(b)} \leftarrow$ sample $n_{\tilde{k}}$ cell design covariates from $\mathbf{C}_h$ corresponding to index $\tilde{k}$ for all $1, \ldots, \tilde{K}$ using $\tilde{\mathbf{Z}}^{(b)}$ for individual-to-cell mapping ;
    $\tilde{\mathbf{y}}_{j,H_1}^{(b)} \leftarrow$ simulate from the full model with covariates $\tilde{\mathbf{C}}_h^{(b)}, \tilde{\mathbf{G}}_{l(j)}^{(b)}, \tilde{\mathbf{Z}}^{(b)}$ ;
    $\tilde{\mathbf{y}}_{j,H_0}^{(b)} \leftarrow$ simulate from the null model with covariates $\tilde{\mathbf{C}}_h^{(b)}, \tilde{\mathbf{G}}_{l(j)}^{(b)}, \tilde{\mathbf{Z}}^{(b)}$ ;
    Fit the selected eQTL model method on $\tilde{\mathbf{y}}_{j,H_1}^{(b)}$ to obtain alternative genotype effect size estimate $\hat{\beta}_{H_1}^{(b)}$ ;
    Fit the selected eQTL model method on $\tilde{\mathbf{y}}_{j,H_0}^{(b)}$ to obtain null genotype effect size estimate $\hat{\beta}_{H_0}^{(b)}$ ;
**end**
Collect $\{\hat{\beta}_{H_1}^{(b)}\}_{b=1}^{B}$ into a vector $\hat{\boldsymbol{\beta}}_{H_1}$ ;
Collect $\{\hat{\beta}_{H_0}^{(b)}\}_{b=1}^{B}$ into a vector $\hat{\boldsymbol{\beta}}_{H_0}$ ;
**for** $x = 1$ **to** $1000$ **do**
    Resample $\hat{\boldsymbol{\beta}}_{H_1}$ with replacement to obtain $\hat{\boldsymbol{\beta}}_{H_1,x}$ ;
    Resample $\hat{\boldsymbol{\beta}}_{H_0}$ with replacement to obtain $\hat{\boldsymbol{\beta}}_{H_0,x}$ ;
    **if** $\beta_{jh} > 0$ **then**
        $p_x \leftarrow$ proportion of $\hat{\boldsymbol{\beta}}_{H_1,x}$ exceeding the $1 - \frac{\alpha}{N_{\text{SNP}} N_{\text{gene}}}$ quantile of $\hat{\boldsymbol{\beta}}_{H_0,x}$ ;
    **else**
        $p_x \leftarrow$ proportion of $\hat{\boldsymbol{\beta}}_{H_1,x}$ below the $\frac{\alpha}{N_{\text{SNP}} N_{\text{gene}}}$ quantile of $\hat{\boldsymbol{\beta}}_{H_0,x}$ ;
    **end**
**end**
Calculate and record the average statistical power $P_h = \frac{1}{1000} \sum_{x=1}^{1000} p_x$ ;
Repeat the above steps for each cell type $h \in \{1, \ldots, H\}$ and record the vector of average statistical powers $\mathbf{P} = \{P_h\}_{h=1}^{H}$ ;

**Fig S25:** Algorithm pseudocode for scDesignPop's eQTL power analysis.
